# Causes of variability in estimates of mutational variance from mutation accumulation experiments

**DOI:** 10.1101/2021.07.07.451460

**Authors:** Cara Conradsen, Mark W. Blows, Katrina McGuigan

**Affiliations:** School of Biological Sciences; The University of Queensland; St. Lucia, Queensland, Australia 4072

**Author notes:** Corresponding Author: Cara Conradsen, School of Biological Sciences, The University of Queensland; St. Lucia, QLD, 4072 AUSTRALIA.

**Keywords:** meta-analysis, Drosophila, life history, morphology, heritability, coefficient of variance, wing, sampling error

## Abstract

Characteristics of the new phenotypic variation introduced via mutation have broad implications in evolutionary and medical genetics. Standardised estimates of this mutational variance, *V_M_*, span two orders of magnitude, but the causes of this remain poorly resolved. We investigated estimate heterogeneity using two approaches. First, meta-analyses of ~150 estimates from 37 mutation accumulation (MA) studies did not support a difference among taxa (which differ in mutation rate) in standardised *V_M_*, but provided equivocal support for standardised *V_M_* to vary with trait type (life history versus morphology, predicted to differ in mutation rate). Notably, several experimental factors were confounded with taxon and trait, and further empirical data are required to resolve their influences. Second, we analysed morphological data from an experiment in *Drosophila serrata* to determine the potential for unintentional heterogeneity among environments in which phenotypes were measured (i.e., among laboratories or time points) or transient segregation of mutations within MA lines to affect standardised *V_M_*. Approximating the size of an average MA experiment, variability among repeated estimates of (accumulated) mutational variance was comparable to variation among published estimates of standardised *V_M_* for morphological traits. This heterogeneity was (partially) attributable to unintended environmental variation or within line segregation of mutations only for wing size, not wing shape traits. We conclude that sampling error contributed substantial variation within this experiment, and infer that it will also contribute substantially to differences among published estimates. We suggest a logistically permissive approach to improve the precision of estimates, and consequently our understanding of the dynamics of mutational variance of quantitative traits.

## INTRODUCTION

The magnitude of per-generational increase in genetic variance due to spontaneous mutations (*V_M_*) is important for a wide range of genetic and evolutionary phenomena, including the maintenance of quantitative genetic variance (Lynch 1988; Barton and Turelli 1989; Johnson and Barton 2005). Much of our understanding of *V*_M_ comes from mutation accumulation (MA) experiments, where populations diverge phenotypically due solely to the neutral fixation of new mutations (Mukai 1964; Halligan and Keightley 2009). Reviews of MA experiments in a range of traits and taxa have reported that mutation increases phenotypic variance in quantitative traits by 10^-4^ to 10^-2^ times the environmental variance of the trait, or 0.02 to 5.1 percent of the trait mean per generation (Houle *et al.* 1996; Lynch *et al.* 1999; Halligan and Keightley 2009). Differences in *V_M_* may cause differences in the magnitude of standing quantitative genetic variation and, ultimately, in rates of phenotypic evolution (Houle 1998; Lynch *et al.* 1999; Houle *et al.* 2017; Walsh and Lynch 2018). However, the causes of variation among estimates of *V*_M_, and thus the evolutionary interpretation of this variability, are not well resolved.

Mutation rate is known to vary widely among species (reviewed in Katju AND Bergthorsson 2019), with further opportunity for differences in per-generation mutation number arising through differences in ploidy, genome size, and/or effective population size (Lynch *et al.* 1999; Lynch 2010; Sung *et al.* 2012). Marked variation in mutation rate has also been observed within species, both among replicated MA experiments (i.e., different founder genotypes: Ness *et al.* 2012; Sung *et al.* 2012; Schrider *et al.* 2013; Ho *et al.* 2020) and among lines within a single MA panel (Huang *et al.* 2016; Ho *et al.* 2020). Resulting differences in mutation number may explain variation in *V_M_* estimates, such as, for example, the fourfold range of 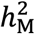 of body size estimated for different MA in *Caenorhabditis elegans* (Azevedo *et al.* 2002; Estes *et al.* 2005; Ostrow *et al.* 2007).

Traits have also been hypothesised to differ in magnitude of *V_M_* due to differences in mutation rate, arising due to differences in the number of contributing loci. Specifically, life history traits are hypothesised to be affected by more loci than morphological traits (Houle 1991; Houle 1992; Houle *et al.* 1996; Houle 1998; Merilä and Sheldon 1999). The magnitude of *V_M_* depends not only on the rate of mutation, but also on their effects, and the relationship between rate and effect size is not well characterised. Besnard *et al.* (2020) demonstrated that the high mutational variance (and relatively rapid evolution) of a vulval phenotype in nematodes was due to a broad mutational target size, rather than large-effect mutation. Whether trait types differ systematically in mutational target size is difficult to assess, as a full catalogue of causal loci is unknown for most quantitative traits (Barton and Keightley 2002; Mackay *et al.* 2009; Yang *et al.* 2010; Rockman 2012). Indeed, emerging evidence that diverse traits, including morphology, are all highly polygenic (Yang *et al.* 2010; Boyle *et al.* 2017) suggests that differences in the distribution of mutational effect sizes (Simons *et al.* 2018), rather than simply in number of contributing loci, might cause heterogeneity in estimates of *V_M_* among traits.

Comparison among trait types is complicated by differences among them in variability and measurement scale, which may influence standardised values. Low mutational heritability (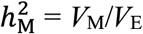, where *V*_E_ is the environmental variance) of life history traits relative to morphological traits has been attributed to greater environmental variance (larger *V_E_*) for life history traits, rather than lower *V*_M_ (Houle *et al.* 1996). Thus, comparison on the coefficient of mutational variance scale (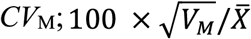, where 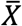 is the trait mean) reveals a different picture, one of greater mutational variance in life history than morphological traits, consistent with the prediction of greater mutational target size (Houle *et al.* 1996).

Other contributions to variation in magnitude of *V_M_* might be revealed by consideration of the MA experimental design itself. The timeframe over which mutations accumulate might influence estimates of *V_M_*. When MA lines are established from a homozygous (heterozygous) base population, estimates of *V*_M_ will be downwardly (upwardly) biased before 6*N_e_* generations (Lynch and Hill 1986). However, *V_M_* is typically estimated after >6*N_e_* generations, suggesting limited contribution of ancestral variation to variability of *V_M_*. Conversely, long-running MA experiments might under-estimate *V_M_* when the cumulative effect of low fitness mutation causes line extinction, or within-line selection against further accumulation (Lynch *et al.* 1999; Estes *et al.* 2004; McGuigan and Blows 2013). A decline in *V*_M_ over time has been observed in some studies (Mackay *et al.* 1995), but not in others (GarcÍa-Dorado *et al.* 2000; Hall *et al.* 2008).

Stochastic sampling from the distribution of mutational effects could also introduce temporal heterogeneity among estimates of *V*_M_. For example, among-line variance estimated before versus after a line(s) fixed a large effect mutation(s) could result in inference of a much larger pergeneration increase in variance at the second time-point relative to the first. Transient within-line segregation of mutations might generate variability in estimates, for example causing temporary inflation of within-line variance (*V_E_*), impacting power to detect among-line variance, and potentially biasing estimates of 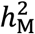 (downward) and *CV*_M_ (upward; see Hoffmann *et al.* 2016). Notably, several studies in nematode have suggested that within-line variance increased over the duration of the MA experiment (Baer 2008; Baer *et al.* 2010; Braendle *et al.* 2010), which may contribute to a pattern of lower estimated *V*_M_ in longer-running MA experiments.

Environmental context within which MA lines are assayed could also contribute to variation among *V_M_* estimates. Several studies have considered the effect of replicable, experimenter-imposed, changes in the environment, including in temperature (Wayne and Mackay 1998), light (Kavanaugh and Shaw 2005) and density (Fry *et al.* 1996). Although the magnitude of *V*_M_ often varies under such environmental manipulations, there is only weak evidence for predicable patterns, such as novel or stressful environments increasing the magnitude of *V*_M_ (Kondrashov and Houle 1994; Martin and Lenormand 2006). Even in carefully controlled laboratory experiments, factors such as food quality or quantity, light, humidity and diurnal timing of collection will vary among individuals or lines within a phenotyping assay, and among assays conducted in different laboratories or at different times within the same laboratory. Such variation may impact estimates of standardized *V_M_* through inflation of within-line variance, similar to the effect of transient, within-line segregation of new mutations. Furthermore, if MA lines differ in their response to this unintended environmental heterogeneity, then genotype by environment (GxE) variance could contribute variation among MA lines, and variability among estimates of *V*_M_, a potential source of variation that has received little attention (but see GarcÍa-Dorado *et al.* 2000).

Here, we combined two approaches to investigate causes of variability in estimates of mutational variance. Given that it has been over twenty years since this variability has been broadly documented and investigated (Houle *et al.* 1996; Houle 1998; Lynch *et al.* 1999), we first conducted a meta-analysis to update tests of the previously implicated causal factors of taxon (Lynch and Walsh 1998; Lynch *et al.* 1999; Halligan and Keightley 2009) and trait type (Houle *et al.* 1996; Houle 1998). We had intended to examine how the number of generations affected *V_M_* (Lynch and Hill 1986; Mackay *et al.* 1995), but MA duration was strongly confounded with taxon (detailed in the Results). Second, we conducted a new empirical experiment in *Drosophila serrata,* in which we repeatedly estimate the among-line (mutational) variance to investigate whether unintended environmental heterogeneity, or transient within-line segregation of mutations can contribute variation among estimates. After accounting for these effects within the data, we finally quantify the magnitude of variation among estimates from a set of 10 wing shape traits to characterise the magnitude of variation among estimates within a trait category.

## METHODS

### Meta-analysis of empirical estimates of mutational variance

#### Literature search

We extracted all papers in seven reviews of mutational variance: Lynch (1988); Keightley *et al.* (1993); Houle *et al.* (1996); Lynch and Walsh (1998); Lynch *et al.* (1999); Halligan and Keightley (2009); Walsh and Lynch (2018). We then searched the Web of Science database on the 11/12/2019 at 4:38 PM AEST for *journal article* document types meeting the topic criteria of “mutation accumulation” and (varia* or “mutat* coefficient*”) and published between 1998 and 2019. These years overlapped Halligan and Keightley (2009) (fitness traits only) and Walsh and Lynch (2018) (brief update on Lynch and Walsh 1998), allowing us to capture papers that may have been excluded from those reviews, as well as those published subsequently.

Further details on the papers identified and preliminary handling steps can be found in Figure S1. For 473 unique papers identified, we screened titles and abstracts, then the full text, for relevance, applying four strict criteria, retaining only studies where the estimates of mutational variance were: 1) quantitative; 2) from spontaneous MA; 3) from MA environmental conditions and; 4) not rereporting of previously published estimates. We excluded six studies of transcriptomic data as the number of traits was much larger than for other trait categories.

#### Meta-analysis data collection

For each of the 65 papers retained after applying the above criteria, mutational parameter estimates were extracted (as described in Table S1), associated with taxon and trait identifiers, and details of the experimental design. Twenty papers not reporting error for the mutational parameters were excluded (Table S1.C). Following initial qualitative assessments of data, we excluded five studies (15 traits) due to low representation of taxon type (one vertebrate, *Mus musculus*; one alga, *Chlamydomonas reinhardtii,* and; two *non-Drosophila* insects: *Daktulosphaira vitifoliae* and *Nasonia vitripennis*), and one study due to low representation of trait type: mitotic cell division traits (Table S1.B).

Where possible, we extracted (or calculated from provided information) both the coefficient of variance (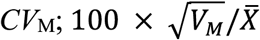, where 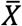 is the trait mean) and mutational heritability (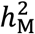; *V_M_*/*V*_E_, where *V*_E_ is the environmental variance) for each trait. As detailed below, estimates were weighted by the inverse of their standard error (SE) in the meta-analysis. Where these were not reported for 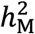 or *CV*_M_, but were for *V_M_* and *V_E_* or 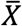, we used a sampling approach to obtain estimates. We sampled from 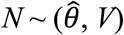 10,000 times, using the *rnorm* function in R [v.3.6.1], where 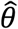 and *V* were respectively the reported parameter value and its standard error. We then calculated *CV*_M_ or 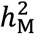 for each of these simulated samples, and obtained the standard error of this sample of estimates. Samples with negative values of *V_M_* are undefined for *CV*_M_; to ensure unbiased estimates of the magnitude of error we calculated *CV_M_* as: 100 x (sign of *V_M_*) x 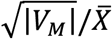. This sampling approach was used to estimate the error for 28% of the 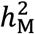 estimates and 61% of the *CV*_M_ estimates analysed (Table S1.A). Two studies (17 estimates) were excluded due to nonsensically large standard error estimates, while a further three estimates (from three studies) were excluded due nonsensical scaled parameter estimates (detailed in Table S1.B). Two extreme values (> 3 SD) of 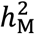 and two of *CV*_M_ were excluded from the analyses (Table S1.B). There were 11 cases with extremely small SE (> 5 IQR below the median); notably, for six of these came from studies where confidence intervals were constrained to be positive, suggesting that this boundary condition had reduced the SE estimate, inflating meta-analysis weights for traits where the mutational variance was not supported. These outliers were excluded from analyses, although results and conclusions were qualitatively consistent when they were included.

#### Predictor variables for the meta-analysis

Estimates came from 11 species, and based on the distribution of estimates, we defined five taxon categories (Figure 1A): Daphnia (*Daphnia pulex* only); Drosophila (*Drosophila melanogaster* [*n* = 68] and *D. serrata* [*n* = 5]); Plant (*Arabidopsis thaliana* [*n* = 12], *Amsinckia douglasiana* [*n*=2] and *Amsinckia gloriosa* [*n* = 1]) and; Nematode (*Caenorhabditis elegans* [*n*=62], *C. brenneri* [*n*=4], *C. briggsae* [*n*=8], *C. remanei* [*n*=5] and *Oscheius myriophila* [*n*=5]). We differ from a previous study (Houle *et al.* 1996; 1998) in considering size of juveniles as morphological (not growth) traits. Reflecting more recent publications, we defined a physiology category (33% of estimates; Figure 1B), which included locomotive, enzymatic and metabolic activity traits (Table S2), which may differ from life-history or morphological traits in mutational target size or environmental sensitivity. We assigned the relatively well-represented life history traits (52% of estimates; Figure 1B) into more narrowly defined sub-categories: total fitness, survival, productivity, and a miscellaneous category (capturing traits such as development time, phenology, longevity and mating success, which were individually less well represented) (Figure 1B; Table S2).

**Figure 1.**
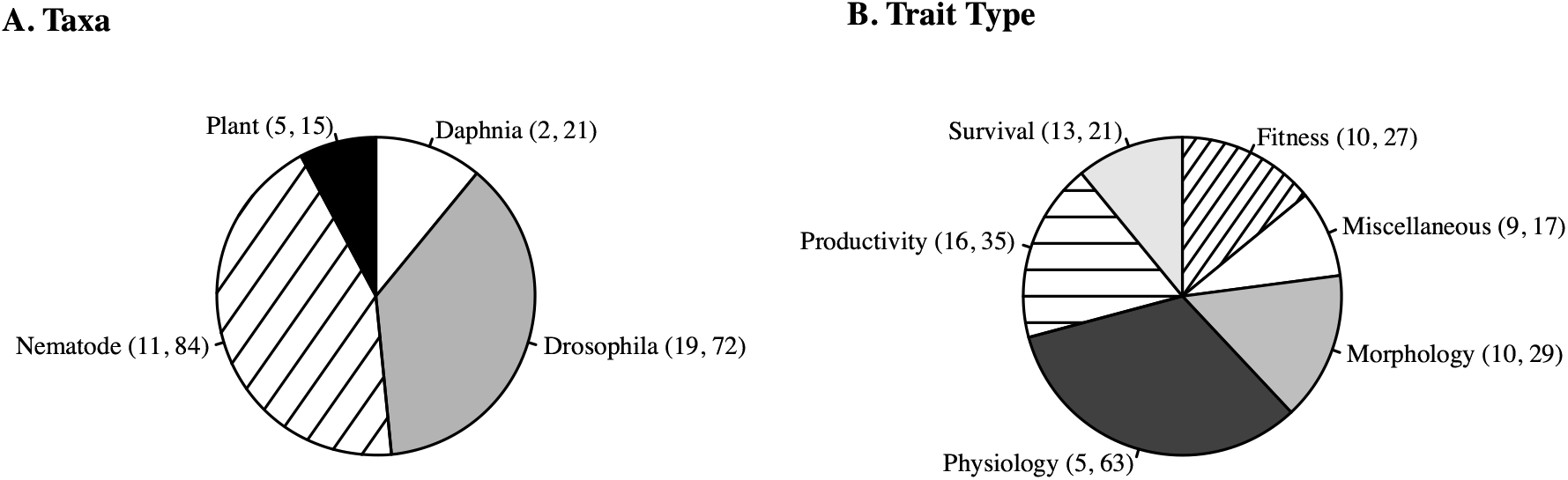
The distribution of published mutational variance estimates by taxon (A) and trait (B) categories. The number of studies (first value in brackets) and estimates (second value) per category are shown. See Methods (and Table S2) for details of the categories and Table S1 for the studies.

#### Meta-analyses of mutational variance estimates

We implemented a mixed model analyses via PROC MIXED in SAS v.9.4 (SAS Institute Inc., Cary, N.C.), using restricted-maximum likelihood (REML) and applying the Satterthwaite approximation to correct the denominator degrees of freedom, to fit the model:

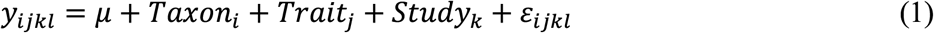

where *y* is the vector of published estimates (either 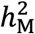 or *CV*_M_), and *μ* was the grand mean; the categorical predicators of taxon and trait (defined above) were fit as fixed effects. Estimates were weighted by the inverse of the standard error (SE) of 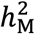 or *CV*_M_, obtained as detailed above. Study was fit as a random effect, accounting for non-independence among estimates within a paper (1-29 estimates per study; median = 3). Studies reporting multiple estimates varied widely in whether these were estimates from different trait types, species (strains), sexes, or time points. Likely reflecting this, most variation not accounted for by the fixed effects was observed at the residual, not study, level (99% for 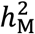; 85% for *CV*_M_). Similarly, while some studies shared the same MA lines (Table S3) fitting a further random effect to account for this non-independence did not explain any variation, a likely consequence of both the unbalanced design (only some studies share lines), and the relative variation of estimates. We investigated different options for fitting heterogenous residuals (e.g., allowing separate estimation of residuals for studies grouped depending on the number of estimates reported), but interpretation of the fixed effects (taxon and trait) were consistent across all investigated models, and we report results only from model (1) above.

### Variation in estimates of among-line variance within the same taxon and trait type: An experiment in *Drosophila serrata*

To what extent do differences in magnitude among estimates of *V_M_* reflect differences in mutation number and/or effect sizes (correlated with the above-investigated proxies of taxon and trait), versus factors such as mutations segregating within MA lines or environmental dependency of mutational effects? We conducted a further experiment to address this question. *Drosophila serrata* is a member of the *montium* species group, endemic to Australia and Papua New Guinea, which has been extensively used in quantitative genetic research, including study of mutational variance (e.g., McGuigan and Blows 2013; Hine *et al.* 2018; Dugand *et al.* 2021). A panel of 200 MA lines were founded from one of the *D. serrata* Reference Genome Panel (DsRGP) lines described in Reddiex *et al.* (2018). These MA lines were each maintained by brother-sister inbreeding for 20 generations, following protocols established by McGuigan *et al.* (2011) to minimise selection. Genome-wide heterozygosity was very low (0.3%) in the DsRGP line that founded the MA lines, and among-line variance for wing traits (defined below) was not statistically supported in the first generation of the MA (S. Chenoweth, pers. comm.).

As detailed below, we applied an experimental design to this MA panel that allowed us to generate repeated estimates of the magnitude of among-line variance over six sequential generations, and characterise the relative contribution to differences among these sequential estimates of i) mutations segregating within the MA lines or ii) unintentional variation in environment. We randomly chose 42 of the MA lines for this investigation based on the median number of MA lines in the reviewed published studies (see Results). The number of MA lines is the relevant degrees of freedom for the among-line variance, and this value (42) allows us to consider the other two effects against a relevant level of sampling error. Quantitative genetic parameters are associated with large sampling errors (Klein *et al.* 1973; Klein 1974; Lynch and Walsh 1998), and the relatively low signal (i.e., few genetic differences) among MA lines will make mutational variance particularly vulnerable to “noisy” estimation, and as such, it is important to document the potential for statistical sampling error to contribute to the observed variation among published estimates.

There were three key aspects of the experimental design that allowed us to test whether segregating variation or unintended environmental variation could explain differences among repeated estimates of among-line variance. First, we increased the population size within each MA line to a minimum of 12 males and 12 females (Figure 2A). Empirical evidence suggests that population sizes as low as ten may be sufficient to prevent fixation of mutations (Estes *et al.* 2004; Katju *et al.* 2015; Luijckx *et al.* 2018). Therefore, we expect no ongoing fixation of mutations among lines during this experiment, and for the repeated estimates of among-line variance to be true replicate sampling of the same mutations (but also test this assumption, as detailed below). We note that these changes in census population size complicate calculation of a per-generation rate of increase in phenotypic variance (Lynch and Hill 1986; Lynch and Walsh 1998); here, we instead focus on the among-line variance, *V_L_*, and do not interpret a per-generation rate of change.

**Figure 2.**
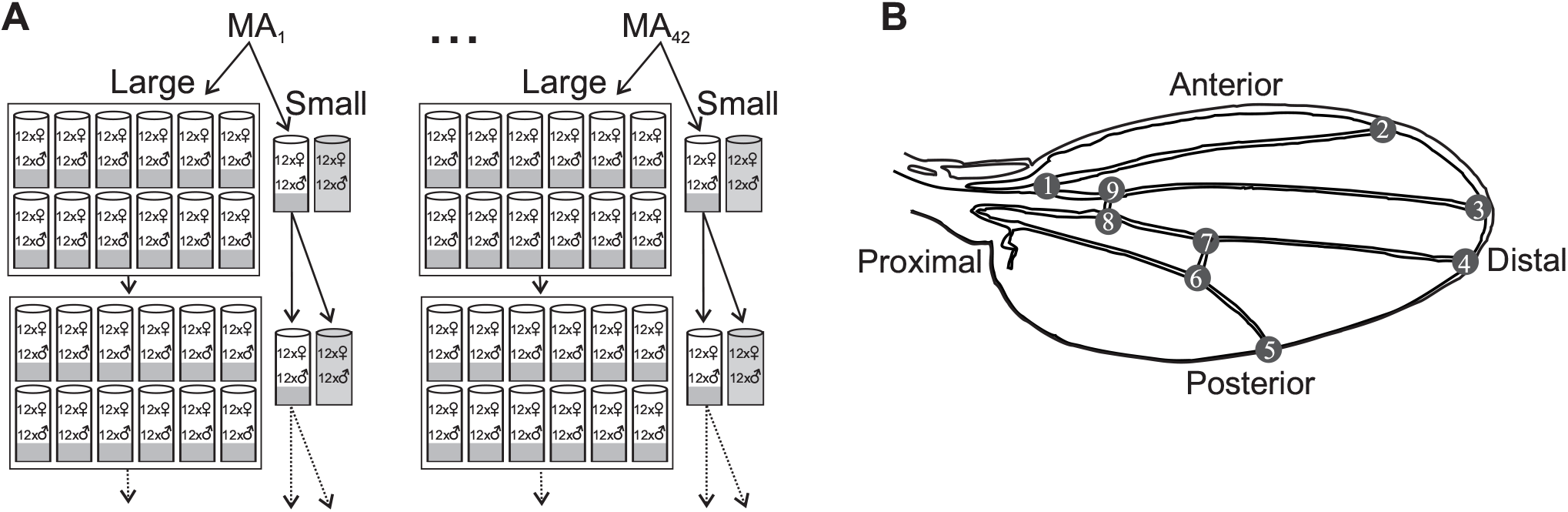
Schematic of design (A) and phenotypes (B) from a manipulative experiment in *Drosophila serrata.* (A) 42 MA lines (evolved through 20 generations of brother-sister mating) each founded two sublines: Small (S; 12 virgin males and 12 virgin females) and Large (L; 144 virgin males and 144 virgin females, distributed evenly among 12 vials). These 84 lines (S and L subline per 42 MA lines) were maintained at these census population sizes for six generations (only two shown here). Each generation, all emergent flies from the 12 vials per L subline were pooled prior to virgin collection. For S sublines, two vials were established each generation; the focal vial contributed offspring to the next generation, while the replicate vial (grey shaded) did not. Each generation, one wing was sampled from each of six randomly chosen males from each of two vials per line (focal and replicate vials for S; randomly chosen two for L). (B) Wing size and shape were characterised from landmarks recorded on an image of each wing: proximal (1) and distal intersections of the radial vein (2); distal intersections of medial (3), cubital vein (4), and distal veins (5) and; the posterior (6, 7) and anterior (8, 9) cross-veins. Inter-landmark distance traits were described by their end-point landmarks (e.g., ILD1.2 was the distance between landmark1 and landmark 2).

The second key aspect of the experimental design was to manipulate the mutation-selection-drift dynamics within an MA line; this was achieved by imposing two, substantially different, population sizes on sublines of each of the 42 MA lines (*N* = 24 versus 288 flies, referred to hereafter as small, S, and large, L, population size treatments: Figure 2A). Segregating variants within MA lines (i.e., mutations that have not yet been fixed or lost) could cause transient inflation of among and/or within line variance (*V_E_*), impacting on both the estimation and scaling of *V_L_*, and this manipulation allowed us to determine the magnitude of this effect. The treatments contrast deterministic evolution of mutations with relatively strong (*s* > ~0.038: *N_e_* ~ 13), versus weak (*s* > 0.003: *N*_e_ ~ 158) fitness effects, based on *s* = 1/2*N*_e_ (Wright 1931; Kimura 1983) and genomic estimates of *N_e_* in MA lines of *D. melanogaster* maintained similarly to our small population size treatment (10 males and 10 females: Huang *et al.* 2016). The S and L treatments therefore had different opportunities for new mutations to increase in frequency within a line, and thus for the magnitude of within-line variance.

The final key aspect of the experimental design was the repeated measures themselves, allowing us to observe the effect of environmental variation on among-line variance. If the phenotypic effects of a mutation are context-dependent (i.e., exhibit genotype by environment [GxE] variance), then unintended differences in assay conditions could contribute heterogeneity among estimates when phenotypic data is collected at different timepoints (or in different laboratories). We randomly sampled the average environmental conditions present within our laboratory by repeatedly sampling the lines (genotypes) over six consecutive generations. Thus, our experiment consisted of applying two population size treatments (S, L) to each of 42 lines (derived from a classical MA experiment, with low among-line variation), where these 84 lines were maintained under the same conditions (12 flies per sex per vial founding each generation, with S and L differing in the number of vials) for six generations (Figure 2A). As detailed below, we consider 11 wing shape and size traits. This allows us to understand the general influences of segregating variation, environment and sampling error for a set of related morphological traits. After accounting for those three factors that are the main focus of the investigation, we also determine whether the magnitude of *V_L_* varies among these traits, allowing insight into potential magnitude of differences in mutational variance among traits within the same category (morphology).

#### Data collection

Each generation, 12 males (six from each of two rearing vials) from each of the 84 sublines were randomly sampled for wing phenotypes (Figure 2). Wings were mounted on microscope slides and photographed using a Leica MZ6 microscope camera and the software LAS EZ v2.0.0 (Leica Microsystems Ltd, Switzerland). A total of 5,135 wings were landmarked for nine positions, defined by wing vein and margin intersections (Figure 2B), using the software tpsDIG2 (Rohlf 2013). The number of wings were evenly distributed across treatments (2,583 in S and 2,552 in L) and generations (~425 per generation, per treatment). Landmarks were aligned using a General Procrustes fit in tpsRelw (Rohlf 2007). Centroid size (CS), the square root of the sum of squared deviations of the coordinates from the centroid (Rohlf 1999), was recorded as a metric of wing size. The aligned X-Y coordinates for each landmark were then used to calculate ten inter-landmark distances (ILDs) (Figure 2B). ILDs scores were re-scaled prior to analysis (multiplied by 100) to aid model convergence. Outliers >3.0 SD from the mean were removed for each of the 11 traits (10 ILD and size) (329 measures across the 56,485 total measures).

#### Analyses of variation in among-line variance estimates

Our experimental design allows us to repeatedly estimate variance among mutation accumulation lines under conditions where we expect the number of mutations fixed among the lines, and their phenotypic effects, to be constant, and thus to investigate other potential causes of variability in estimates. We first treat the data from each generation and population size treatment as independent experiments of similar size (number of lines and individuals sampled per line) to typical MA experiments. To estimate among line variance from these 12 “experiments” for each of the 11 traits we fit the following model using REML in PROC MIXED in SAS v9.4 (SAS Institute Inc., Cary, NC.):

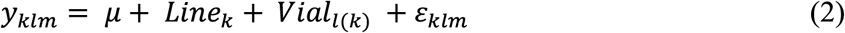

where *y_klm_* was the trait value for the *m*th wing (individual), from the *l*th vial, within the *k*th line, *μ* was the trait mean; Line and replicate rearing Vial (nested within line) were fit as random effects, along with the among-individual variation (residual error, *ε*). We used REML-MVN sampling (Meyer and Houle 2013; Houle and Meyer 2015; Sztepanacz and Blows 2017) to estimate confidence intervals, sampling 10,000 times from 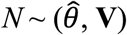 using the rnorm in R [v. 3.6.1], where 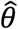 was the vector of REML random effect parameter estimates, and **V** was their inverse Fisher information matrix, 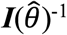. We similarly estimated the confidence intervals for the trait mean, sampling based on least-squares mean and standard error estimates output from model (2). The samples of random effect variances were not constrained to the parameter space (i.e., could be negative), allowing inference of statistical support when the lower 5% CI did not encompass zero (a one-tailed test); this approach is equivalent to a Log Likelihood Ratio test (LRT) (Dugand *et al.* 2021). Here we are interested in general patterns of variability among these 12 “experiments”, and thus do not correct for multiple testing.

As detailed in the Results, substantial heterogeneity in magnitude was observed among the 12 replicate estimates of *V_L_* per trait. We considered the potential contribution to this heterogeneity from unintentional heterogeneity in the culture conditions among sampling time points (generations) or between replicate measures of the same MA line within a generation (the S and L treatments). First, to determine if simple effects of variability in culture conditions on trait scale could account for the variability of among-line (mutational) variation, we placed estimates on a heritability (*V_L_* / *V_E_* where *V_E_* was the sum of among and within vial variances) or coefficient of variance 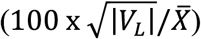 scales, and calculated confidences intervals by applying these equations to each of the 10,000 samples described above (and applying the sign correction for coefficients of variance as detailed in the meta-analysis methods). We further explored the relationship between *V_L_* and the scaling parameters by regressing the 12 estimates of *V_L_* on the corresponding estimates of *V_E_* or trait mean.

Second, we determined whether mutational effects changed in response to the unintended changes in culture conditions, with such genotype by environment causing differences among sequential estimates of *V_L_*. Therefore, we extended this investigation, following GarcÍa-Dorado *et al.* (2000) in treating different generations as different environments to formally test the null hypothesis that there was no GxE variance within the S or L treatments, using PROC MIXED and REML to fit:

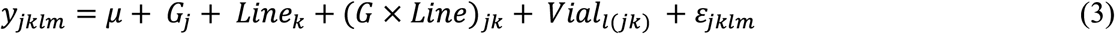

where the fixed effect of generation (G) accounted for differences in trait mean among generations and the random effect of G(eneration) × Line estimated the variation in genetic effects among generations (where generations represent different local environments). For the component of *V_E_* (i.e., Vial and residual), generation-specific effects were modelled (using the GROUP statement) to account for among-generation heterogeneity in the magnitude of *V_E_*. This model was applied to each trait within each population size treatment, and statistical support for G × Line (and for Line) was determined using log-likelihood ratio tests (LRT, 0.5 d.f.: Self and Liang 1987; Littell *et al.* 2006) to compare model (3) to reduced models that did not fit G x Line (or did not fit Line). We applied the Benjamini-Hochberg method (Benjamini and Hochberg 1995) to correct for multiple hypothesis testing (within each random effect), using a conservative 5% false discovery rate (FDR). Sampling based on the REML variance estimate and the Fisher information matrix, as detailed above, was used to estimate confidence intervals for plotting.

While non-zero generation by line variance could reveal the presence of environment-specific mutational effects, it could alternatively be explained by changes in the frequency at which mutations were segregating within or among lines. In contrast to environmental heterogeneity, we expect these evolutionary processes to systematically differ between the two population size treatments due to the different efficacy of selection in the S versus L sublines, and the independent sampling of mutations in the sublines after they were established. Differences between S and L are predicted to increase with increasing time since divergence. For each of the 11 traits, we analysed all data (from both L and S) collected within a single generation, using PROC MIXED and REML to fit:

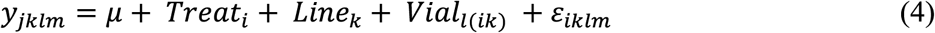

where Treatment was fit as a fixed effect to account for differences in trait mean between L and S panels of sublines within that generation. Vial and residual are as described for model (2). At the among-line level, we took advantage of the paired subline design to model the between treatment variance-covariance matrix. We employed LRT to test two hypotheses. First, we determined whether, for these analyses within a generation, there was support for differences between treatments in the magnitude of *V_E_*. Mutations that are segregating (*i.e.,* occur at frequencies other than 0 or 1) within an MA line will contribute to variation both among-vials and the residual. We compared a model in which one (common to both Treatments) among Vial variance and one residual variance were estimated to a model in which Treatment-specific variances were modelled at both levels (fit using the GROUP statement). Second, we tested whether the two copies of the MA lines had diverged from one another by testing the hypothesis that the correlation between the paired sublines was <1.00 (implemented using the PARMS statement). To correct for multiple hypothesis testing (within each hypothesis), we employed a FDR correction as described above.

Finally, as there was little support for varying mutational effects (no GxE) or number (no divergence between S and L) contributing to the apparent heterogeneity among repeated estimates per trait (detailed in Results), we use our data to revisit the question of whether traits inherently differ from one another in the magnitude of mutational variance. We obtained a single estimate of *V_L_* per trait by using PROC MIXED in SAS to fit:

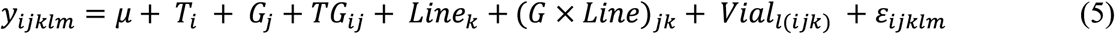

where all effects are as described above, including the fixed effects of population size Treatment (T), generation (G) and their interaction (TG), as well as the random effects of Line, Generation by among-Line, Vial and residual. We obtained REML-MVN confidence intervals for each parameter, as described above. To test whether observed differences in *V_L_* among traits were due to differences among them in scale, we took the among-line (*V_L_*) estimates from model (5) and regressed them on the corresponding estimates of environmental variance or on the squared trait mean. These regressions were applied to the REML parameter estimates, and to each of 10,000 samples of these parameters to determine statistical significance (95% CI of slope did not include zero).

## RESULTS

### Meta-analysis of published mutational variance estimates

Our final meta-analysis data set consisted of 154 estimates of 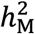 and 148 estimates of *CV_M_*. These estimates (excluding extreme outliers; Table S1.B) of 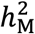 ranged from 2.50 x 10^-5^ to 1.02 x 10^-2^, while *CV*_M_ ranged from 0.13 to 7.32. We predicted that differences in genome size and/or genomic mutation rate would cause differences in the magnitude of mutational variance among taxa. However, there was no statistical support for a difference in mutational variance among the taxon categories (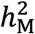: *F*_3,24.2_ = 2.28, *P* = 0.1044; *CV*_M_: *F*_3,17.5_ = 1.24, *P* = 0.3261), although 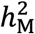 estimates from Plants were markedly lower than estimates from Daphnia and Drosophila (Figure 3A). We predicted that differences among traits in the number of contributing loci would cause differences in the magnitude of mutational variance but that due to differences among trait categories in environmental sensitivity, the rank of trait types would differ between 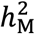 and *CV_M_*. These predictions for trait type were partially met as the pattern did differ between the two standardisations, but trait categories only differed in the magnitude of *CV*_M_ (*F*_5,37.1_ = 3.86, *P* = 0.0064), not 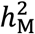 (*F*_5,85.9_ = 0.40, *P* = 0.8497) (Figure 4). Overall, these factors (taxon and trait category) accounted for 1.64% of the variation in estimates of 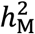 and 9.88% of variation among *CV_M_* estimates.

**Figure 3.**
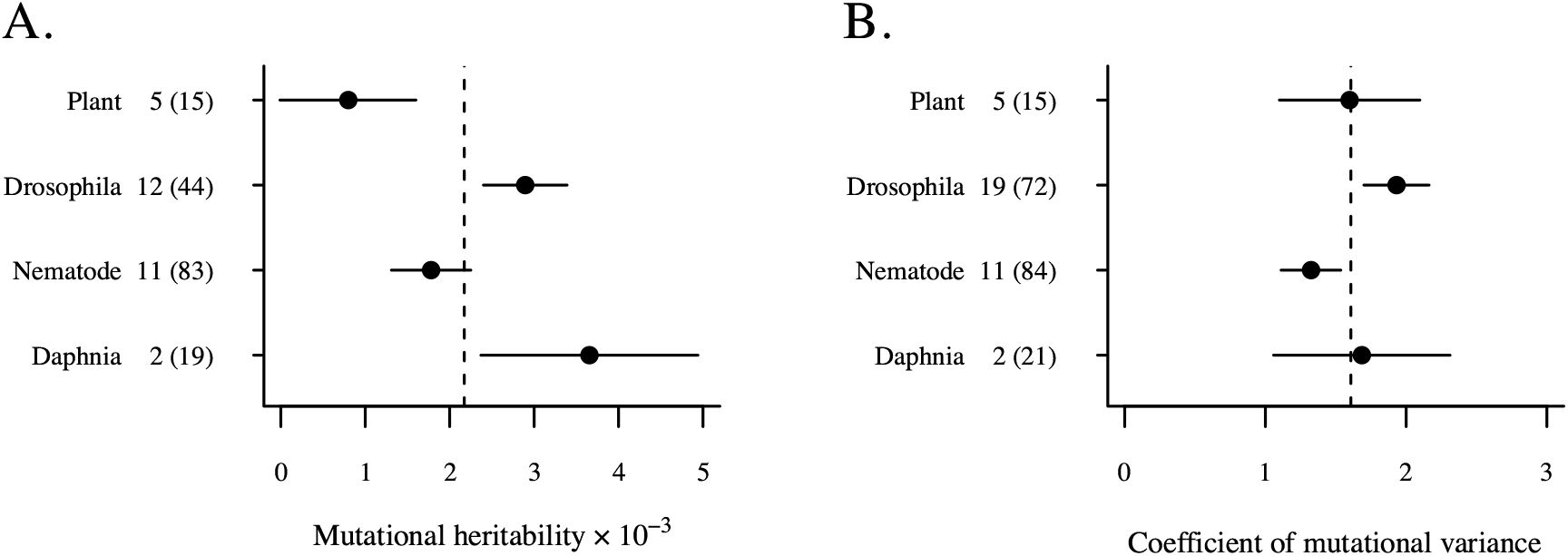
Variation in estimates of (A) mutational heritability and (B) coefficient of mutational variance across taxon categories. Plotted are the least-squares mean estimate (± SE) from the analysis of model (1). The number of studies (and estimates) analysed for each category are shown. The dashed line indicates the global mean value.

**Figure 4.**
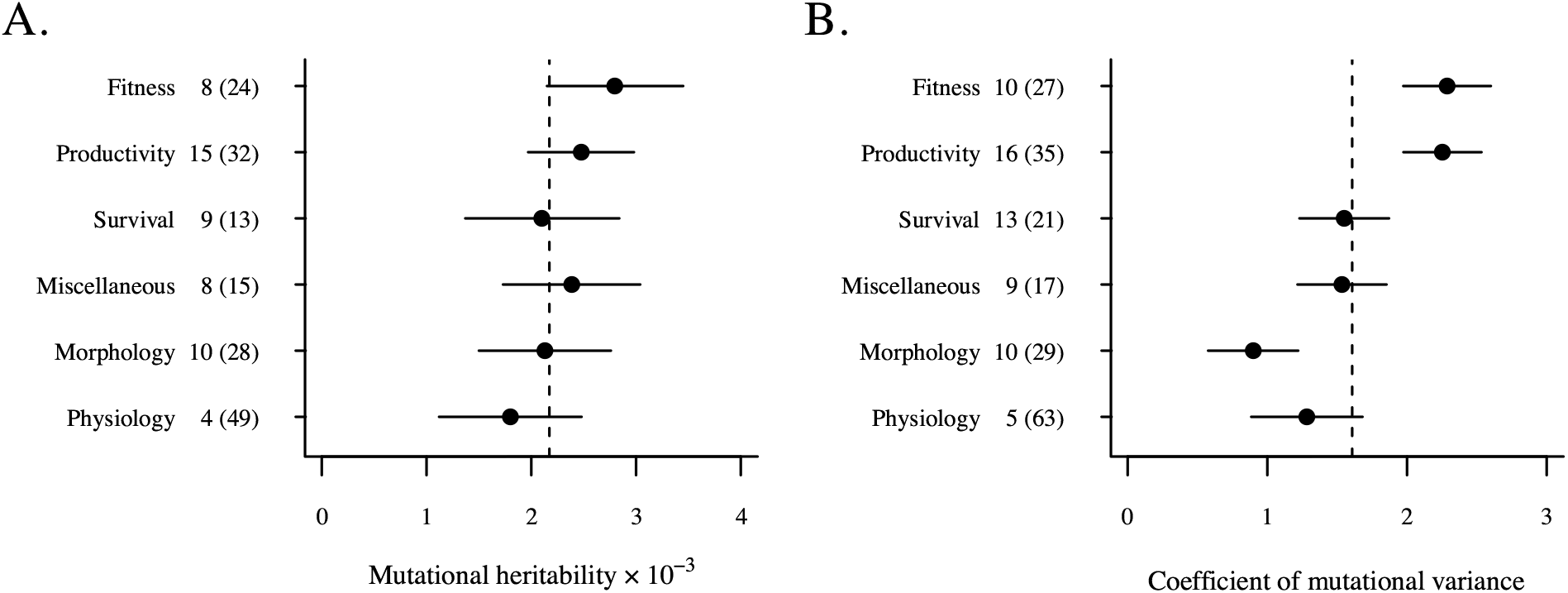
Variation in estimates of (A) mutational heritability and (B) coefficient of mutational variance across trait categories. Plotted are the least-squares mean estimate (± SE) from the analysis of model (1). The number of studies (and estimates) analysed for each category are shown. The dashed line indicates the global mean value

Although not statistically supported, it is notable that the among-trait trend did not follow predictions for 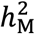: fitness traits had the largest average 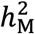, not the lowest as expected (Figure 4A). Following Houle *et al.* (1996), we also analysed *V_E_* (fit model (1) to 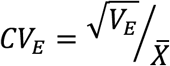). There was no statistical support for a difference among traits in the magnitude of *CV_E_* (*F*_5,25.2_ = 1.32, *P* = 0.2887); morphology (average *CV_E_* = 7.4) and physiology (71.6) differed the most, with life history traits having intermediate values (e.g., fitness = 42.6) (Figure S2A). For *CV_M_*, the statistically supported differences did follow the predicted pattern, with morphological traits having the smallest *CV_M_* and fitness the largest (Figure 4B). Survival notably had lower *CV*_M_ than productivity and fitness (Figure 4B), although surviving to reproduce was a component of fitness. Physiological traits had similar magnitude of *CV*_M_ to morphological traits, and lower than any life history trait category (Figure 4B).

While the lack of observed difference in scaled estimates of *V_M_* among taxa may reflect a true commonality among species in this important evolutionary parameter, aspects of the MA design also differed markedly among taxa. Estimates from Plants were derived from MA experiments that were of short duration (maximum 25 generations) relative to other taxa (median 44, 75 and 214 for Drosophila, Daphnia and Nematode, respectively) (Figure S2B). As mutations arise independently in each MA line, the number of MA lines maintained may also influence the number of mutations that similar duration MA could sample, and the potential for sampling of rarer mutational effects (i.e., from the tails of the distribution of effect sizes) to influence estimates; while the median number of MA lines was similar in Nematode (43) Plant (50) and Drosophila (52), it was substantially lower in Daphnia (8) (Figure S2C). While mutational variance was estimated for multiple types of traits in every taxon, most data from Plants was for fitness, while most data from Daphnia was for morphological traits, and Drosophila and Nematode were the only two taxon categories that contributed estimates for physiological traits (Figure S2D).

### Variation in estimates of among-line variance within the same taxon and trait type

Within the *D. serrata* experiment, we first determined the heterogeneity in among-line variance (*V_L_*) under the assumption that mutation number and effects were constant, treating the 12 *V_L_* estimates per trait as independent mutation accumulation experiments. There was substantial variation among the 12 estimates per trait, with some differences of over an order of magnitude (Figure 5; Table S4). Notably, there was a fourfold difference between the smallest and largest estimates of *V_L_* for size (CS), comparable to the fourfold differences among reported estimates of 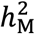 for body size in *Caenorhabditis elegans* (Azevedo *et al.* 2002; Estes *et al.* 2005; Ostrow *et al.* 2007). Predictably given this heterogeneity in magnitude of effect (i.e., in *V_L_*), there was also inconsistent statistical support for the presence of *V_L_* for most traits, despite consistent sample sizes in each of the 12 “experiments” (Figure 5; Table S4). Thus, we might draw very different conclusions about the magnitude of *V_L_* for a trait, depending on which “experiment” we had conducted (Figure 5).

**Figure 5.**
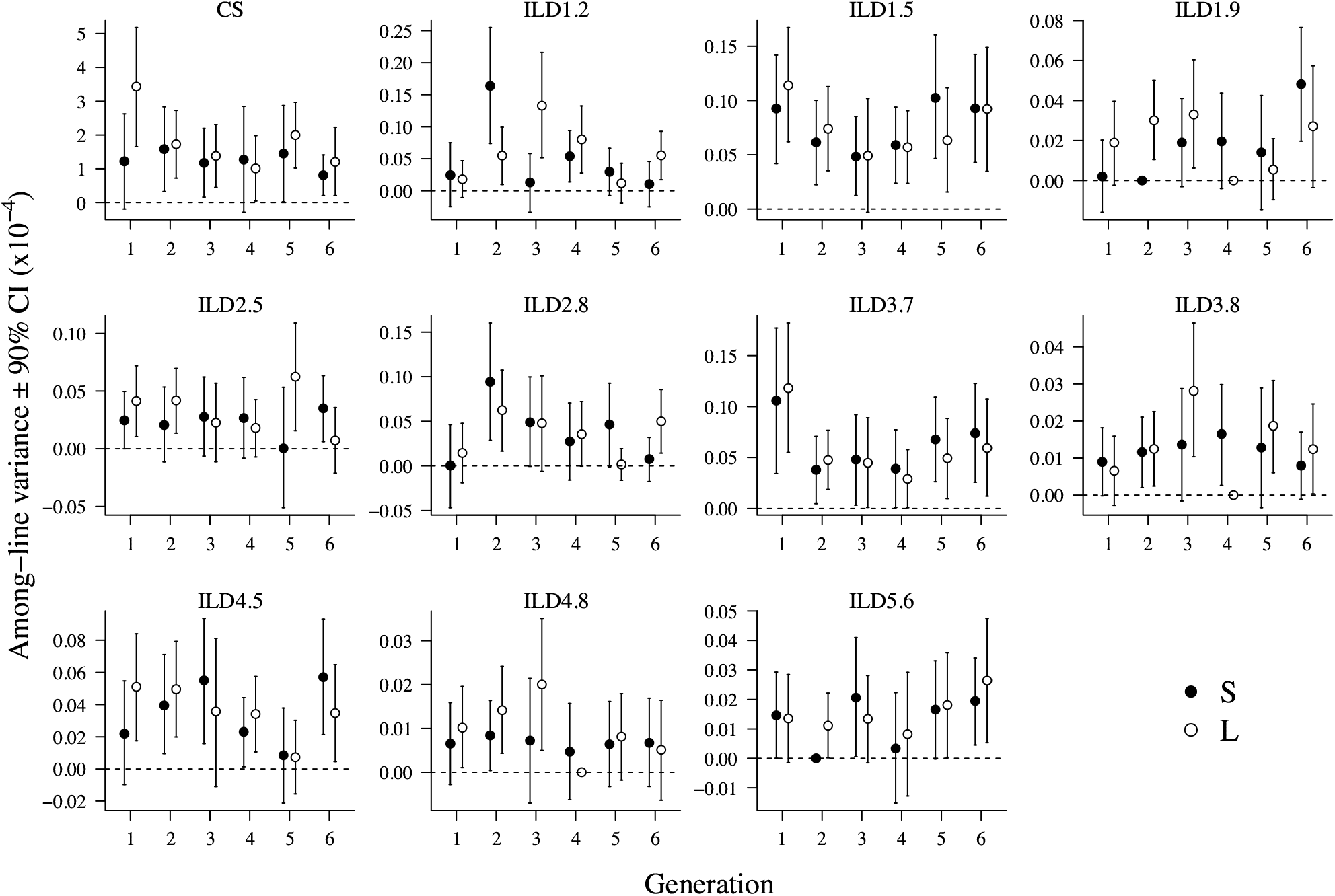
Among-line variance estimates across six generations in an experiment in *Drosophila serrata.* Variances were estimated independently for each trait (panel; see Figure 2B for trait definitions) in each generation (x-axis) for each of the two population size treatments (Small: solid circles; Large: open circles). Plotted are the REML point estimate, and the REML-MVN 90% confidence intervals (CI). The dashed horizontal line indicates zero; estimates for which the lower CI did not overlap zero were interpreted as statistically supported. Where REML estimates of among-line variance were zero, no CI are plotted.

Due to the changes in *N_e_* within this experiment, we do not place these *V_L_* estimates on a per-generation scale (i.e., do not calculate *V_M_*). However, there is no trend for *V_L_* to increase through time (i.e., no signal of ongoing divergence through fixation of mutations), or to diverge between the different population size (*N_e_*) treatments (Figure 5) (addressed further below). Therefore, calculating *V_M_* is not expected to eliminate the heterogeneity in estimates.

Reporting mutational variance estimates as 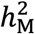or *CV_M_* facilitates comparison among estimates by accounting for inherent differences in scale. Although here the 12 estimates come from the same trait, scale differences may still arise through typical effects of any unintended variation in culture conditions (occurring among generations or between the replicate S and L sublines) on non-genetic trait variance (*V_E_*) or mean. Both *V_E_* and the trait mean varied substantially among the 12 repeated estimates for all traits (Figures S3, S4; Table S4). However, this variation in *V_E_* and trait mean was independent of the observed variation in *V_L_*; regressing the 12 estimates of *V_L_* on their corresponding estimate of *V_E_* or trait mean supported only one slope (ILD3.7, *V_L_* on mean) as statistically different from zero (although this did not remain significant following FDR correction) (Figure 6; Table S5). Consistent with this pervasive independence of *V_L_* from the scaling factors for these repeated measures of the same trait, when the 12 estimates were placed on either a heritability (Figure S5; Table S4) or coefficient of variance scale (Figure S6; Table S4), the variation among them was of a similar magnitude to that observed for *V_L_* itself (i.e., the variation plotted in Figure 5). Plotting the scaled estimates (*h*^2^ or *CV*) against their respective numerator (*V_L_* or 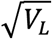) and denominator (*V_E_* or mean) illustrates the predominant contribution from variation in *V_L_* to variation in the scaled estimates (Figure S7). Overall, the 12 estimates of *V_L_* are more variable than the corresponding estimates of *V_E_* or trait mean, with variability of the scaled estimates (*h*^2^ or *CV*) more similar to *V_L_* than to their respective scaling factor (Figure S8).

**Figure 6.**
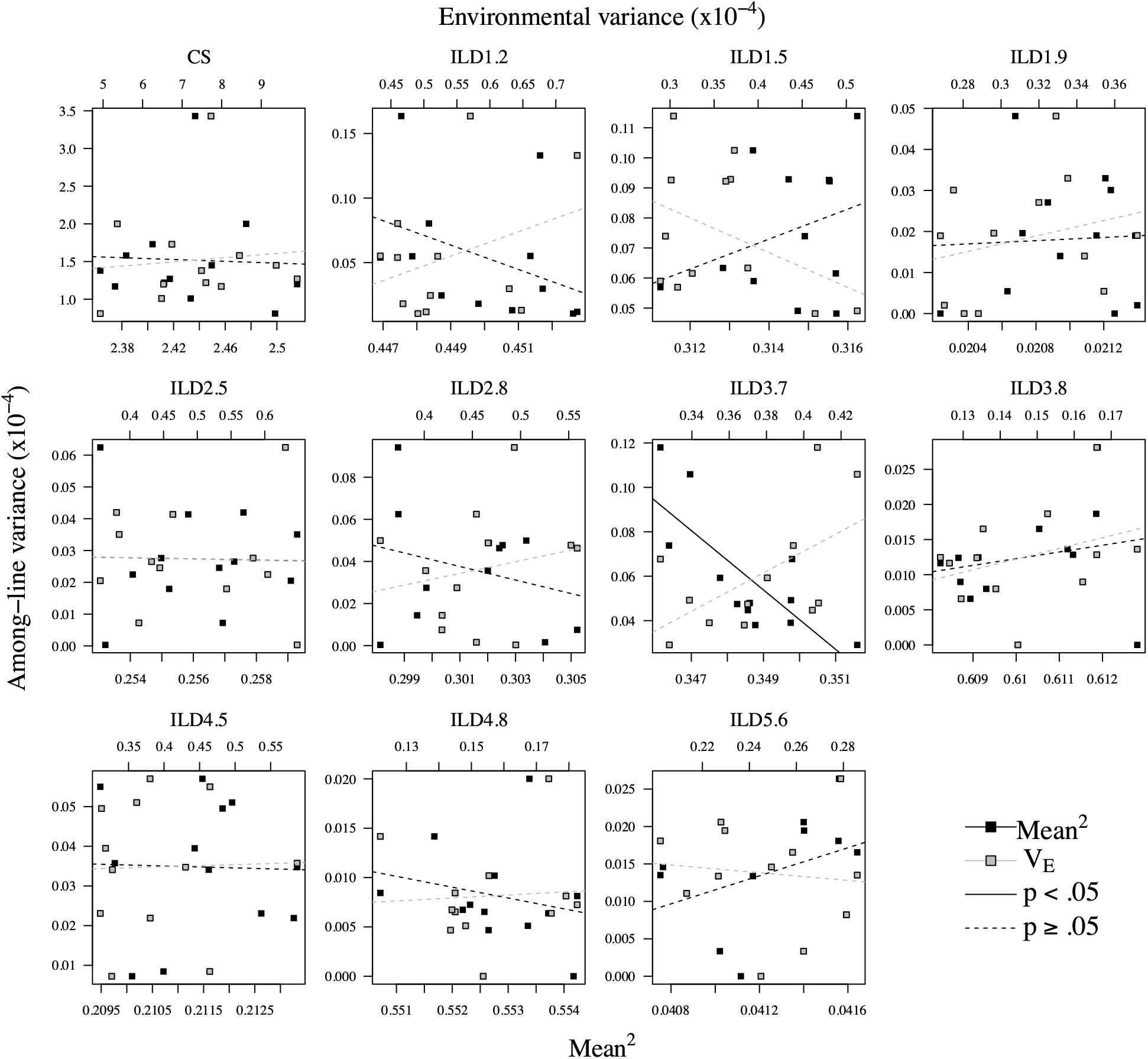
Among-line variance estimates for *Drosophila serrata* wing traits plotted as a function of trait mean or variance. The 12 estimates of among-line variances for each of the eleven wing traits (panels; see Figure 2B for trait definitions) are plotted against the corresponding (i.e., same generation and treatment) squared trait mean (bottom x-axis, black symbols) or environmental variance (summed among and within vial variances; top x-axis, grey symbols). All regression statistics are reported in Table S4; only the effect of mean^2^ on *V_L_* of ILD3.7 was significant at *P* < 0.05, although it does not remain significant after applying a 5% FDR correction.

To compare the variability among published estimates of 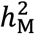 and *CV_M_* in similar morphological traits (excluding bristle traits: 19 estimates, eight each from Daphnia and Nematode, three Drosophila and one Plant) to the variability among the *V_L_* estimates for *D. serrata* wing trait traits, we calculated the coefficient of variance (cv = standard deviation / mean) for each set of estimates. The cv of all 19 published 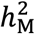 (0.80) and *CV_M_* (0.82) estimates was above the median cv of the 11 *D. serrata* traits on the observed *V_L_* scale (0.55), *V_E_*-scale (*h*^2^: 0.54) or mean-scale (*CV*: 0.39), but nonetheless within the same range: cv of *V_L_* (*V_E_*-scale; mean-scaled) ranged from 0.30 (0.37; 0.15) up to 0.92 (0.85; 0.62) across the 11 traits (Figure S8). Within taxa, cv of published 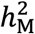 (*CV_M_*) estimates ranged from 0.29 (0.08) for the three Drosophila estimates up to 0.97 (0.38) in Daphnia, with a median of the three within-taxon (Drosophila, Daphnia and Nematode) cv’s of 0.64 (0.24). Thus, overall, the heterogeneity among repeated *V_L_* estimates in *D. serrata* is of a similar magnitude to the variation among published estimates of the same trait type.

Having established that variation in the magnitude of *V_L_* is not a simple consequence of varying scale (*V_E_* or mean), we investigated other putative causes. In addition to the general effects on scale, unintended differences in culture conditions among generations could also affect *V_L_* if mutations had context-dependent effects on the trait (i.e., GxE), as characterised by generation by among-line variance. GxE was statistically supported in only four cases, with only two remaining significant at a 5% FDR (LRT for CS, in L: 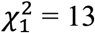, *P* = 0.0001; LRT for ILD2.5 in L: 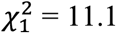, *P* = 0.0005) (Figure 7A). Thus, for these two traits, the analysis suggests that mutational effects, and the magnitude of among-line variance, may depend on the specific conditions under which the traits were assayed.

**Figure 7.**
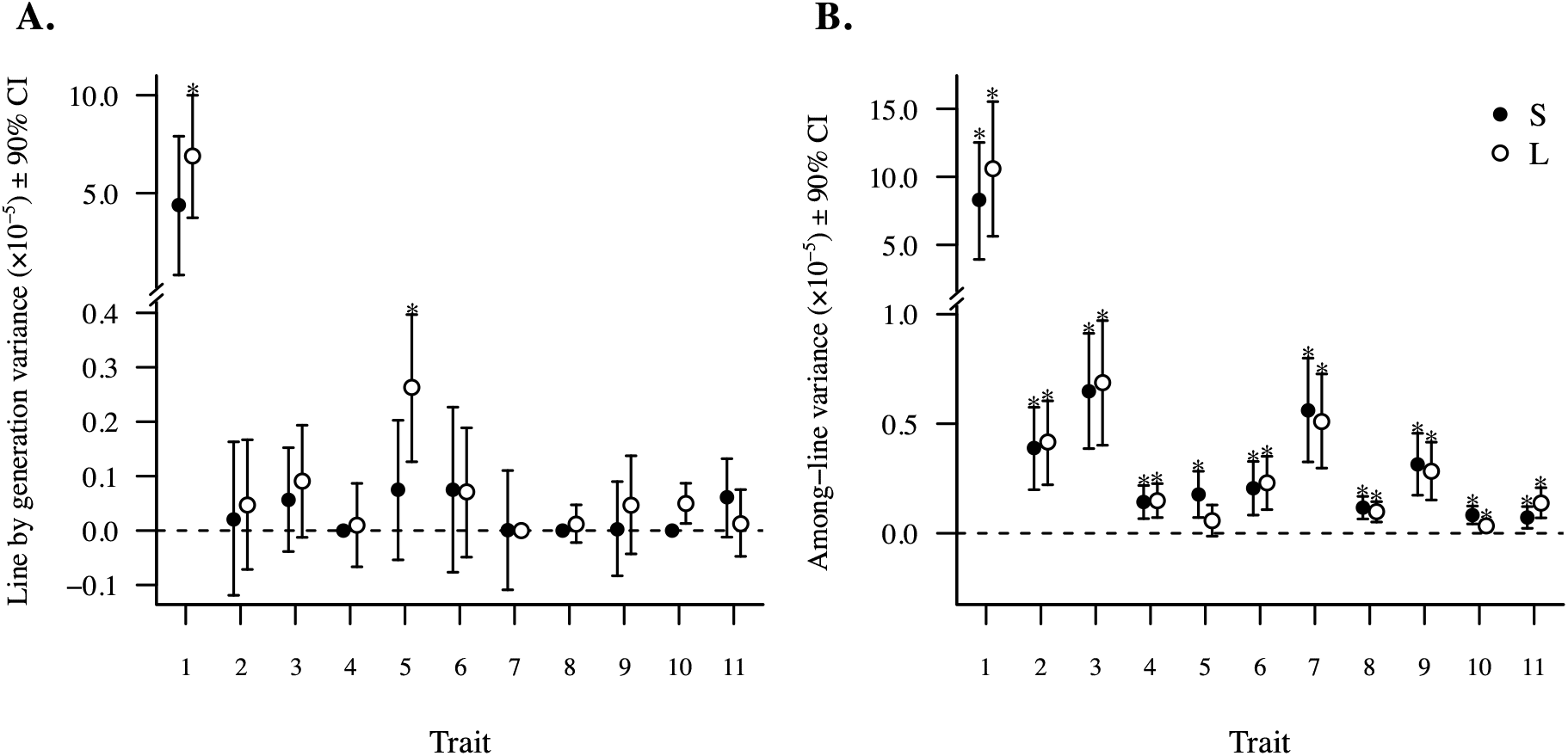
Estimates of variance from an experiment in *Drosophila serrata.* (A) Among-line by generation (GxE) variance and (B) among-line variance estimated for 11 *D. serrata* wing traits (x-axis), in two different population size treatments (Small: solid circles; Large: open circles). Plotted are the REML point estimates (from model 3) and the REML-MVN 90% confidence intervals (CI) around this. The dashed horizontal line indicates zero; statistical significance was inferred where the lower 5% CI did not overlap zero. After applying a conservative 5% FDR correction, two estimates in (A) and 21 in (B) remained significant (asterisk above CI).

Ongoing mutation-drift-selection processes could contribute to variation among the 12 estimates, where the S and L treatments are expected differ in the potential effects of these processes on both within and among-line variance. Segregating variation within a line will contribute to the estimate of *V_E_*, and we determined whether the S and L treatments differed in the magnitude of *V_E_*, analysing each of the 11 traits within each of the six generations separately. Eight of the 66 estimates of *V_E_* differed significantly between S and L at *P* < 0.05, but only one remained significant at 5% FDR (CS in generation 5; Table S6). There was no statistical support for the S and L sub-lines founded by each of the 42 original MA to have diverged from one another in the mutations they carried, either through initial sampling when lines were founded, or through (near) fixation of mutations arising after establishment of the sub-lines. Specifically, the among-line correlation between S and L sub-lines was not statistically distinguishable from 1.0 for any trait in any generation (Table S6).

Finally, we obtained a single estimate of among-line variance for each trait to determine whether the magnitude of *V_L_* was consistent among the 10 wing shape traits, which are expected to share a genetic basis, and developmental pathways (e.g., Mezey *et al.* 2005; Neto-Silva *et al.* 2009). The among-trait heterogeneity (i.e., non-overlapping confidence intervals: Figure 8A; cv = 0.77) was larger than the median variability among the repeated estimates per trait (0.55; see above), and comparable to the variability among published estimates of mutational variance in morphological traits (0.80, detailed above). Variation among shape traits (excluding size) in *V_E_* accounted for ~18% (95% CI: 0.049 - 0.383) of this variation (*β* = 0.064 [95% CI: 0.050 - 0.106]), but variation in trait mean did not account for any (*β* = 5.3 x 10^-7^ [-0.74 x 10^-8^ - 1.86 x 10^-6^]; *R*^2^ = 0.003 [95% CI: <0.001 - 0.027]) (Figure 8B). Establishing whether these differences are informative of the inherent genetic architecture, or are a manifestation of the stochastic nature of mutation, requires repeating the estimation using either the same or a different genetic background to determine if consistent differences among traits persist. When wing size was also considered, overall scale influenced *V_L_*, with much of the variation among estimates accounted from by the scaling factors (*V_E_*: *R*^2^ = 0.995 [0.978 - 0.998]; Mean^2^: *R*^2^ = 0.921 [0.903 - 0.925]), suggesting that there was little difference in the magnitude of underlying mutational variance between wing size and the shape traits (Figure 8B).

**Figure 8.**
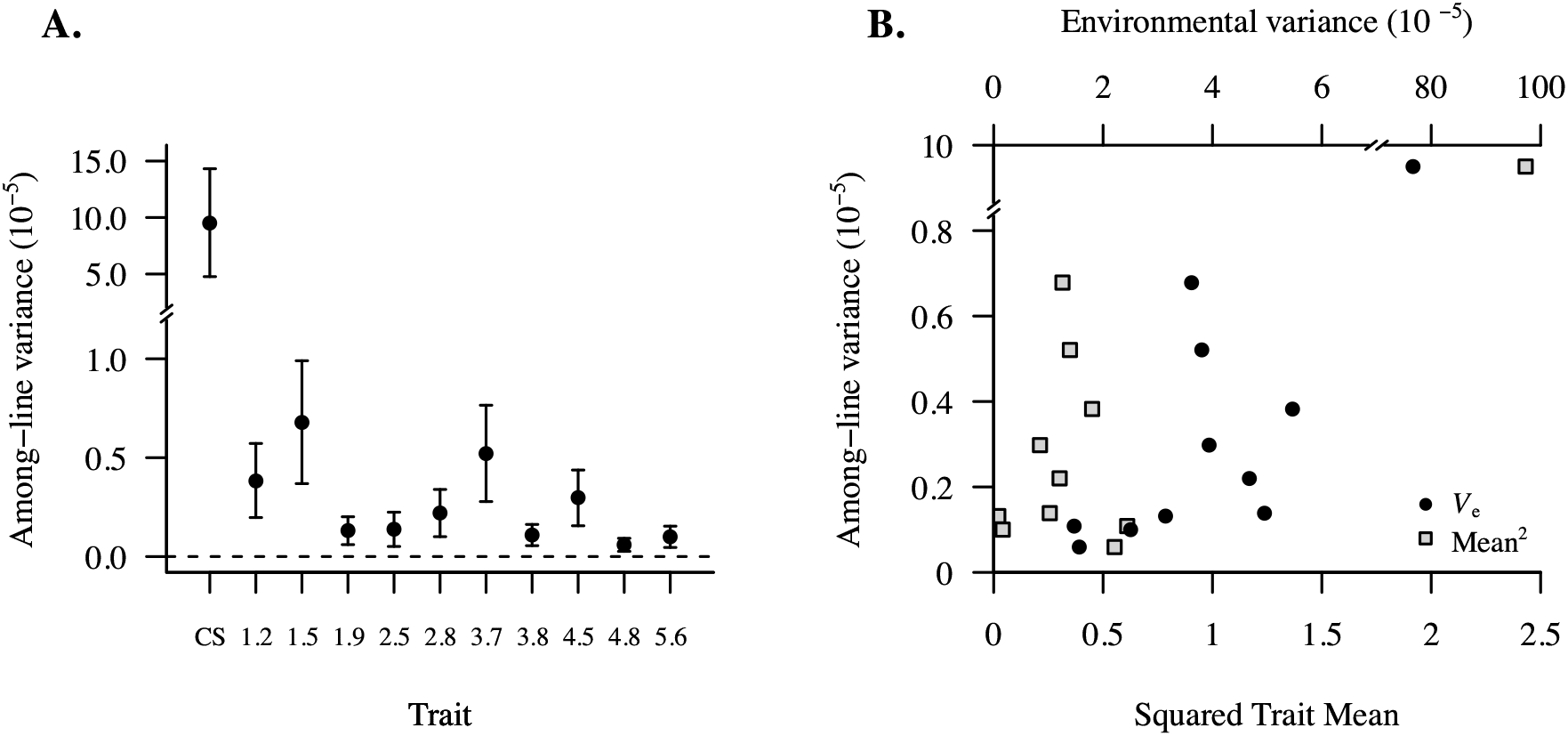
Among-line variances for eleven wing traits in *Drosophila serrata.* (A) Among-line variance, *V_L_* REML estimates (and 90% CI) (model 5; see Figure 2B for trait definitions) are plotted. Dashed line indicates zero. (B) REML estimates of among line variance (points in panel A) were plotted against the corresponding estimate of environmental variance (black circles, top x-axis) or mean squared (grey squares, bottom x-axis) of the trait. Regression results are reported in text.

## DISCUSSION

Although numerous estimates of mutational variance have been published, it remains unclear what contributes to the ~two orders of magnitude difference among these estimates. Our meta-analytic investigation provided some support for a difference among trait types in the magnitude of mutational variance, but also revealed substantial confoundment between potential causal factors. Analyses of data from a manipulative experiment in *D. serrata* suggests that, for the morphological traits under consideration, factors such as unintended heterogeneity in environmental conditions or transient segregation of mutations within MA lines may contribute little to the variation among estimates. Given this experimental design, and the evidence that mutation number and effect did not typically cause differences among repeated estimates, we conclude that substantial variability among repeated estimates of the among-line variance must reflect sampling error. Below we discuss the specific outcomes and limitations of both approaches, and the implications our analyses have for future work characterising mutational input to quantitative genetic variation.

### Effects of taxon and trait type on the magnitude of mutational variance

Given the ~four-fold higher per site mutation rate (Katju and Bergthorsson 2019), and slightly larger genome of *A. thalania* relative to *C. elegans* we predict (assuming the same mutational effect sizes) ~five times more mutational variance in the Plant than Nematode taxon categories. However, the meta-analysis did not support a difference among taxa in the magnitude of mutational variance, and the observed (strong but non-significant) pattern in 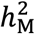 contradicted this rank prediction, with Plants (to which *Arabidopsis* contributed most estimates) having substantially smaller 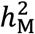 than other taxa (Figure 3A). Taxon categories differed substantially in the number of generations (Figure S2B), and the number of genomes (MA lines) (Figure S2C) sampled. However, scaling predicted genomic mutation by this opportunity for mutation also fails to predict the trend, with Daphnia MA experiments predicted to sample the fewest mutations but observed to have the largest 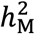 (Figure 3B). We suggest that further MA experiments, decoupling the confounded effects of MA duration and trait type from taxon, are warranted to determine whether *V*_M_ does vary among taxa. Advances in accessibility of genome data provides substantial scope for such experiments to explicitly estimate relevant genomic parameters (e.g., frequency spectra for different types of mutations across putatively causal genes) alongside the phenotypic variation generated by those mutations (Katju and Bergthorsson 2019). Furthermore, given evidence that epigenetic mutations arise more frequently than genetic mutations (e.g., van der Graaf *et al.* 2015; Beltran *et al.* 2020), we suggest that the potential contribution of epimutations to patterns of heterogeneity of *V_M_* should be explicitly assessed in future studies.

Houle *et al.* (1996); (see also Lynch and Walsh 1998; Lynch *et al.* 1999) concluded that life history traits had lower 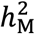 and higher *CV_M_* than morphological traits, a pattern that is also observed in standing genetic variation (e.g., Houle 1992; Hansen *et al.* 2011; but see Hoffmann *et al.* 2016). However, this conclusion was not supported by our analysis, where the trend was for fitness and productivity to have the highest 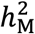 (non-significant) as well as highest *CV_M_* (Figure 4). As expected given this shared pattern, 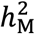 and *CV_M_* estimates were positively correlated (Spearman’s correlation coefficient: 0.309, *N*=117, *P* = 0.0007). Although with reverse rank (life history traits having lowest values), standing genetic variation estimates have also been reported to be positively correlated between the two scales (*h*^2^ and *CV*) when biologically uninformative *CV* estimates were excluded (Hoffmann *et al.* 2016). Garcia-Gonzalez *et al.* (2012) highlight the potential for skewed data distributions to inflate (deflate) *CV*, an issue that may be particularly relevant to estimates of *CV_M_*. While strong bias toward mutations that decrease mean fitness has been reported (Halligan and Keightley 2009), bias in other traits is less well-established. If trait types differ in the magnitude of directional bias of mutational effects, this may also result in differences in skew, and exaggerate differences between trait types on the *CV_M_* scale. Again, resolution of the key question of whether differences among traits in 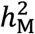 and *CV_M_* reflect differences in mutation number and/or effect size may depend on further genomic data.

### The contributions of unintended environmental variation, mutation-drift-selection processes, and sampling error to variation in the magnitude of mutational variance

We observed substantial variation among repeated estimates of *V_L_* each of 11 wing traits measured in *D. serrata* (Figure 5), resulting in variation among scaled (*h*^2^ or *CV*) estimates that was of comparable magnitude to the differences observed among published estimates. Although both *V_E_* and trait mean also varied among repeated measures (Figures S3, S4), this heterogeneity was substantially less than that observed for *V_L_* (or *h*^2^ or *CV*) (Figure S8). Given the evidence that mutational effects can vary among environments (Kondrashov and Houle 1994; Martin and Lenormand 2006), we tested the effects on the magnitude of *V_L_* resulting from unintentional and undocumented minor changes in culture conditions (e.g., density, humidity or temperature), such as may occur among phenotype assays conducted at different times or in different laboratories. Variation in mutational effects among phenotypic assays (generations) was supported in only two cases (Figure 7A). GarcÍa-Dorado *et al.* (2000) also found evidence of GxE among consecutive generations for one (sternopleural bristle count) of four traits investigated. Notably, in *D. serrata,* wing size, which might be particularly sensitive to variation in energy availability (or competing energetic demands) (Cavicchi *et al.* 1985; Bitner-MathÉ and Klaczko 1999), exhibited the strongest GxE (Figure 7A). Our results, and those of GarcÍa-Dorado *et al.* (2000) suggest that changes in mutational effects with environment may contribute to heterogeneity among published estimates of some traits, which may reflect differences in trait environmental sensitivity, or potentially in the covariation of environmental sensitivity and mutational effect size (Lynch *et al.* 1999; GarcÍa-Dorado *et al.* 2000).

The mutation-drift process itself may also contribute to variability among published estimates due to effects on both the within-line variance (transient inflation leading to increased magnitude of *V_E_* but not *V_L_*) and among-line variance (transient contribution to *V_L_* of additive or dominant mutations that are subsequently lost via random sampling). We introduced an ~order of magnitude difference in census size in paired sets of MA sub-lines to manipulate the mutation-drift-selection processes. However, analyses did not support an effect of population size on either within- or among-line variation, with wing size again an exception, where there was some evidence that relaxed selection allowed the S treatment to accumulate greater within-line variance (Table S6). The effect of segregating variation can be expected to be greater at smaller population sizes (e.g., when *N* = 2, mutations can reach within-line frequency of 75% before being lost by drift) than considered here, and so may play a greater role in explaining variation among estimates from classical MA breeding designs. But, nonetheless, this factor did not account for the substantial heterogeneity that was observed among the 12 estimates per trait within the current study.

Rejecting general contributions from environmental variation and transient segregation of mutations as explanations of the heterogeneity among the 12 repeated estimates of *V_L_* for the wing shape traits, we conclude that the observed variability is largely the consequence of sampling error. Lynch *et al.* (1999) suggested that a substantial part of the order of magnitude range of 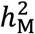 reported for *D. melanogaster* may be due to sampling error. Here, we observed the magnitude of heterogeneity among the 12 repeated estimates of *V_L_* to be similar to the heterogeneity among published, scaled estimates of mutational variance in morphological traits, consistent with their prediction. We observed that *V_L_* estimates varied markedly more than the other estimated parameters (Figure S8), as expected given that quantitative genetic parameters are associated with relatively large sampling errors. Notably, *V_E_* was more variable among the 12 estimates than trait mean was, which may lead to greater variability among estimates of 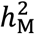 than *CV_M_* (Figure S8). We designed this experiment to mimic an average MA sample size, considered a trait expected to have relatively low experimental noise (residual variation), but expected small effect size relative to many MA (due to the relatively few generations; see e.g., Vassilieva *et al.* 2000). While traits and MA panels will vary in their vulnerability to sampling error, we nonetheless suggest that greater consideration must be given to the consequences of this error when designing experiments. The heterogeneity among repeated estimates resulted in the total confidence range for each trait spanning a far greater region than suggested by the error estimated for each repeated estimation of *V_L_* (Figure 5), indicating that within-study estimates of error do not fully capture the uncertainty in estimates.

The sequential repeated-measures experimental design provided greater statistical control over the experimental noise, allowing us to consistently detect statistically significant mutational variance in all traits (Figure 8A), including in traits for which very few of the 12 estimates were distinguishable from zero (e.g., ILD2.8; Figure 5). While increasing sample sizes within a generation is likely to have similarly improved estimate precision, this can be logistically prohibitive in some systems. Given these limits, our analysis highlights the potential benefits of short-term repeated measures (sequential generations) to improve estimate precision, and power to detect small effects. Repeated measures of lines at relatively large generation intervals have also been utilised to estimate *V_M_* as the slope of the regression of among-line variance on generation (Vassilieva *et al.* 2000; Houle and Nuzhdin 2004; McGuigan *et al.* 2011), which may also improve estimation.

Understanding the contribution that mutations make to evolutionary and genetic phenomena relies on accurate estimates of the phenotypic variance generated by new mutation. Our meta-analysis of empirical estimates of mutational variance was unsuccessful in clearly resolving causes of variation due to confounding of predictors, and inconsistent patterns. Our manipulative experiment suggested that sampling error may contribute substantially to estimate variability, and demonstrated that repeated measures over few (e.g., sequential) generations provides a simple but effective approach to address this and improve inference. Overall, further empirical studies are needed to fully assess how both general and study specific factors influence *V*_M_ estimates, where improved precision and replicability in estimates will consequently advance broader evolutionary questions such as those addressing the maintenance of quantitative genetic variance (Barton and Turelli 1989; Johnson and Barton 2005; Walsh and Lynch 2018).

## Data Availability

Both analysed datasets are available at doi 10.6084/m9.figshare.14913051

## Acknowledgments

We thank Stephen F. Chenoweth for providing us with the *Drosophila serrata* MA lines, and Adam Reddiex, Nicholas Appleton, Jack Price and Derek Sun for their help with maintaining the flies and collecting the data. We also thank Jan Engelstädter for suggesting the combinational approach, and Emma Hine for contributions to preparing figures. This manuscript was improved by the input of three anonymous reviewers and the AE.

## SUPPLEMENTARY TABLES AND FIGURES

Supplementary Table S1 **Information on mutational variance estimates obtained from the literature.**

Supplementary Table S2 **Broad and narrow trait categories.**

Supplementary Table S3 **Mutation accumulation (MA) lines shared across studies.**

Supplementary Table S4 **Estimates of mean and variances within each trait, generation and treatment in *D. serrata***

Supplementary Table S5 **Results of regression of the among-line variance estimates on either the trait mean or on the environmental variance in *D. serrata*.**

Supplementary Table S6 **Statistical support for the effect of population size treatment on (A) among-line and (B) among-line by generation variance estimates.**

Supplementary Figure S1 **Diagram of systematic literature search for the meta-analysis of mutational variance estimates from mutation accumulation (MA) experiments.**

Supplementary Figure S2. **Distribution of the (log-transformed) number of generations (A) and the (log-transformed) number of lines (B) of mutation accumulation experiments in different taxon categories.**

Supplementary Figure S3 **Variation in wing trait mean across six generations of an experiment in *Drosophila serrata.***

Supplementary Figure S4 **Variation in *V_E_* across six generations of an experiment in *Drosophila serrata.***

Supplementary Figure S5 **Contribution of variability in phenotypic variance to estimates of among-line variance in *D. serrata.***

Supplementary Figure S6 **Contribution of variability in trait mean to variability of among-line variance estimates in *D. serrata.***

Supplementary Figure S7 **Relative contributions of heterogeneity in the scaling parameter and among-line variance to the variation in the scaled estimates from *D. serrata*.**

Supplementary Figure S8 **Variation in estimates of each genetic variation parameter in *D. serrata.***

**Table S1.**
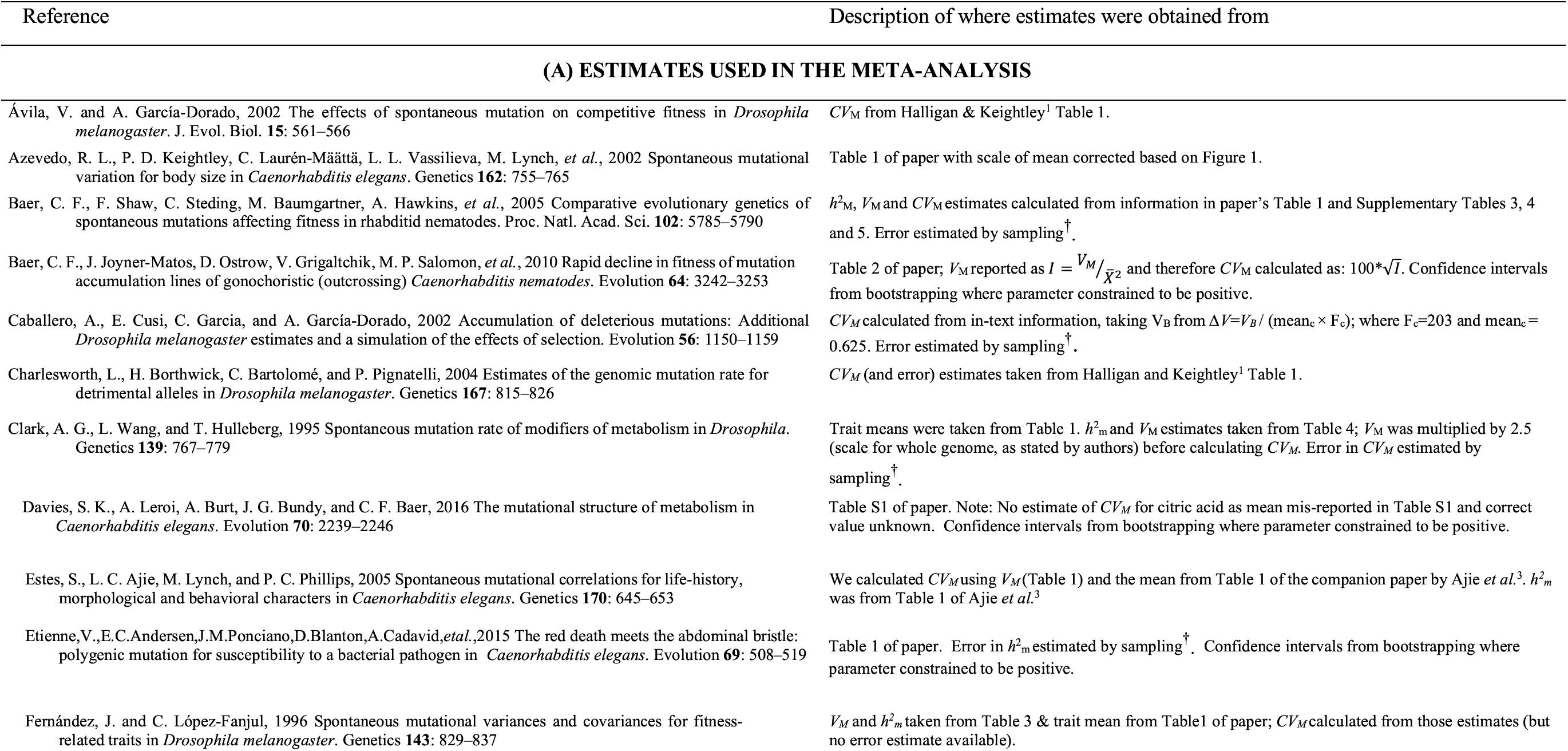
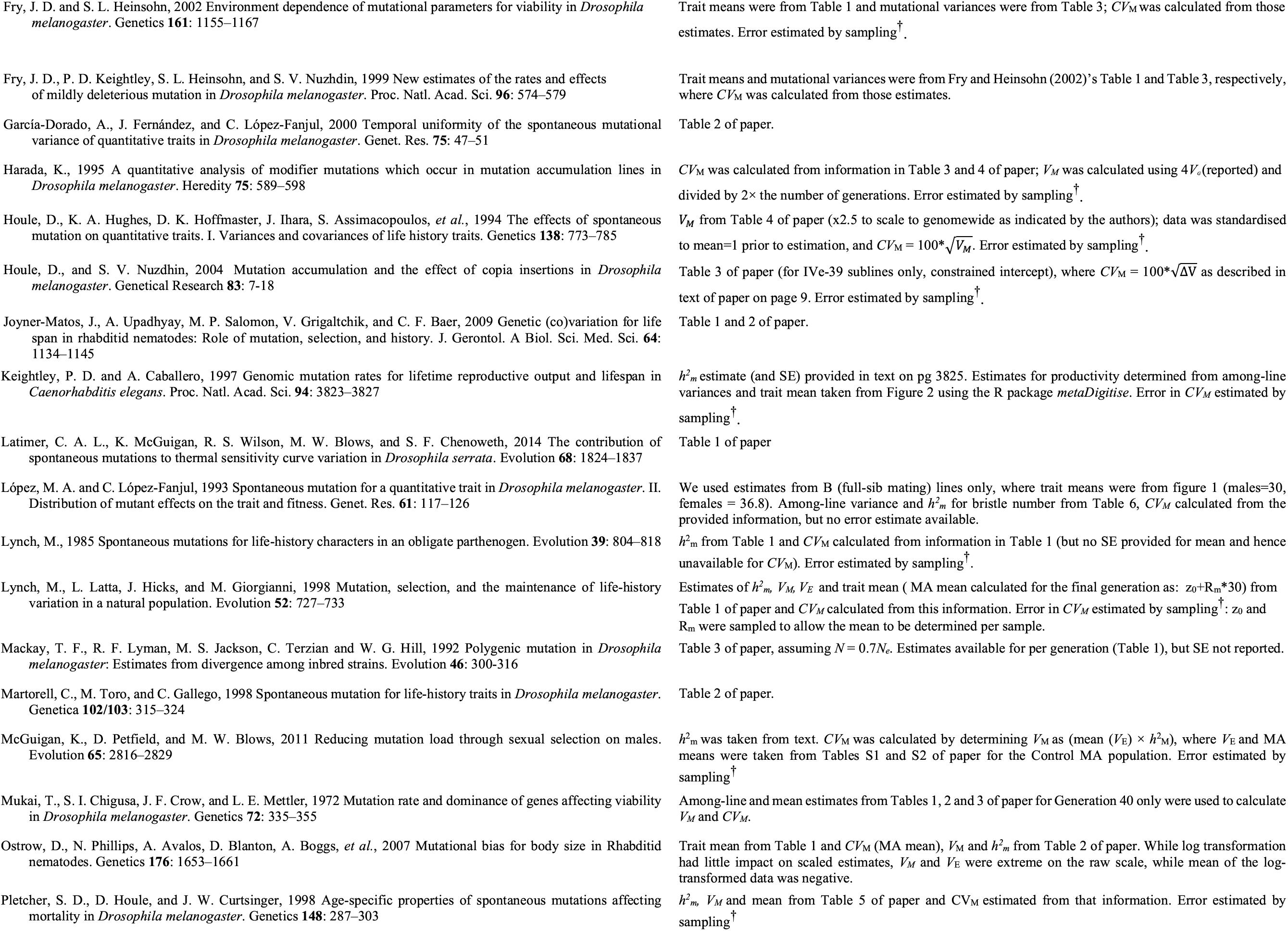
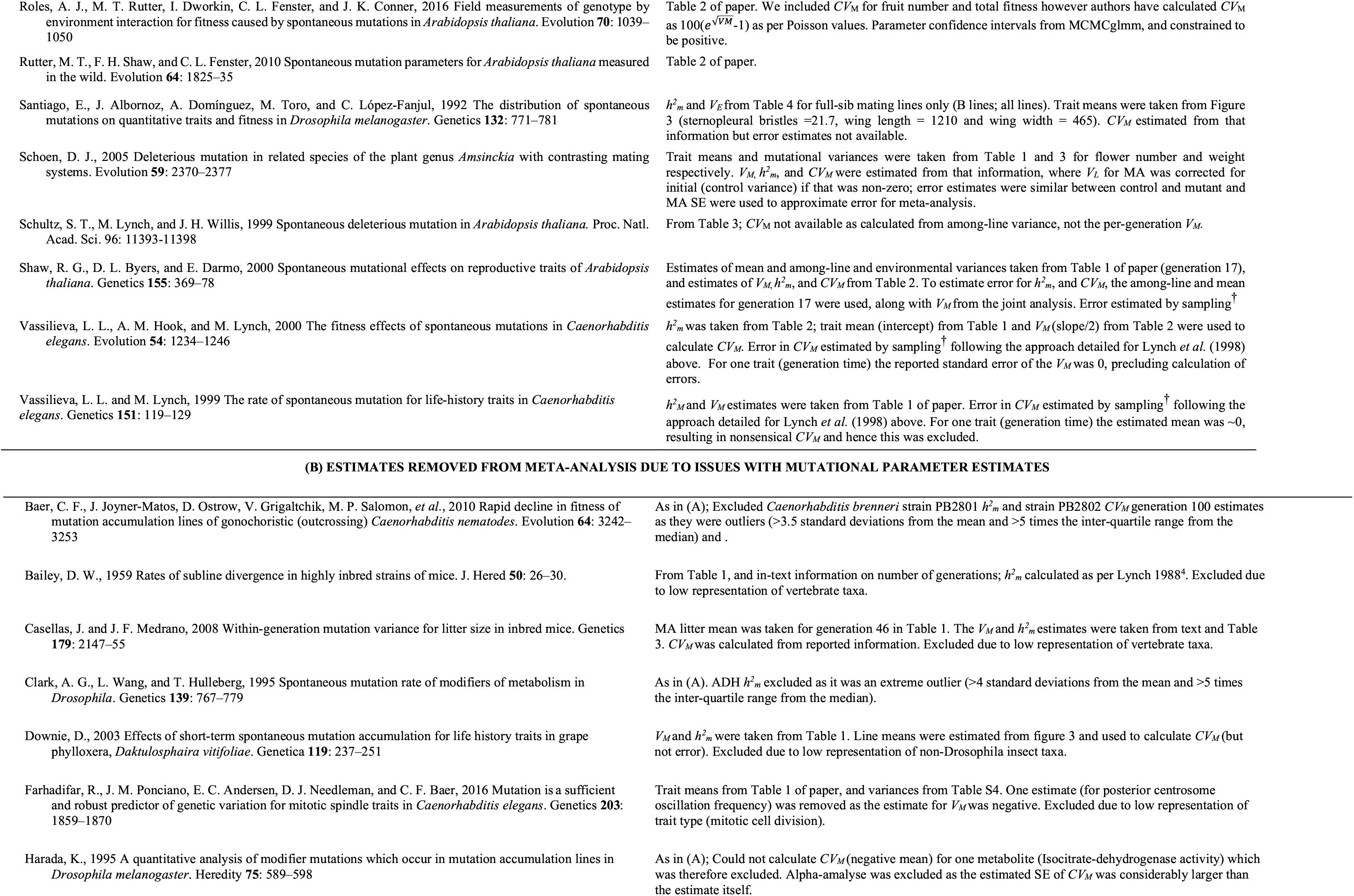
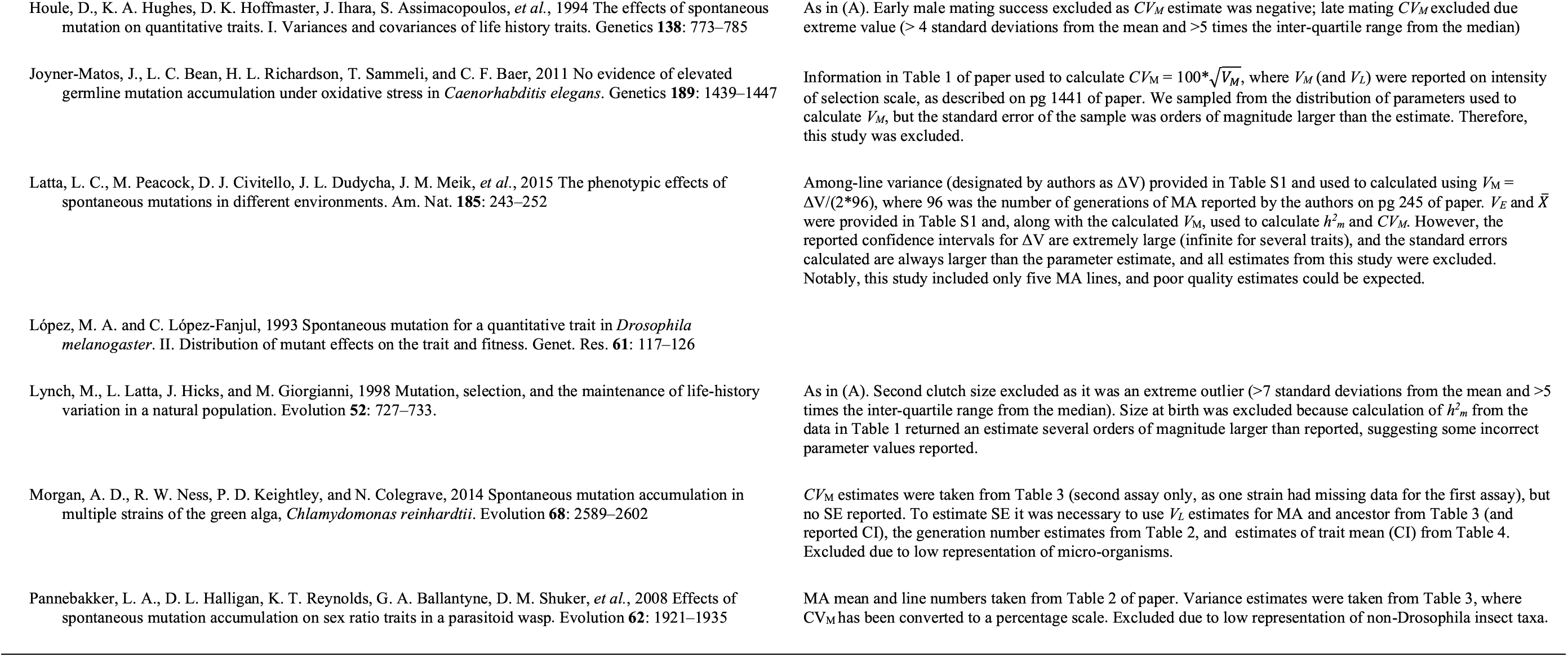
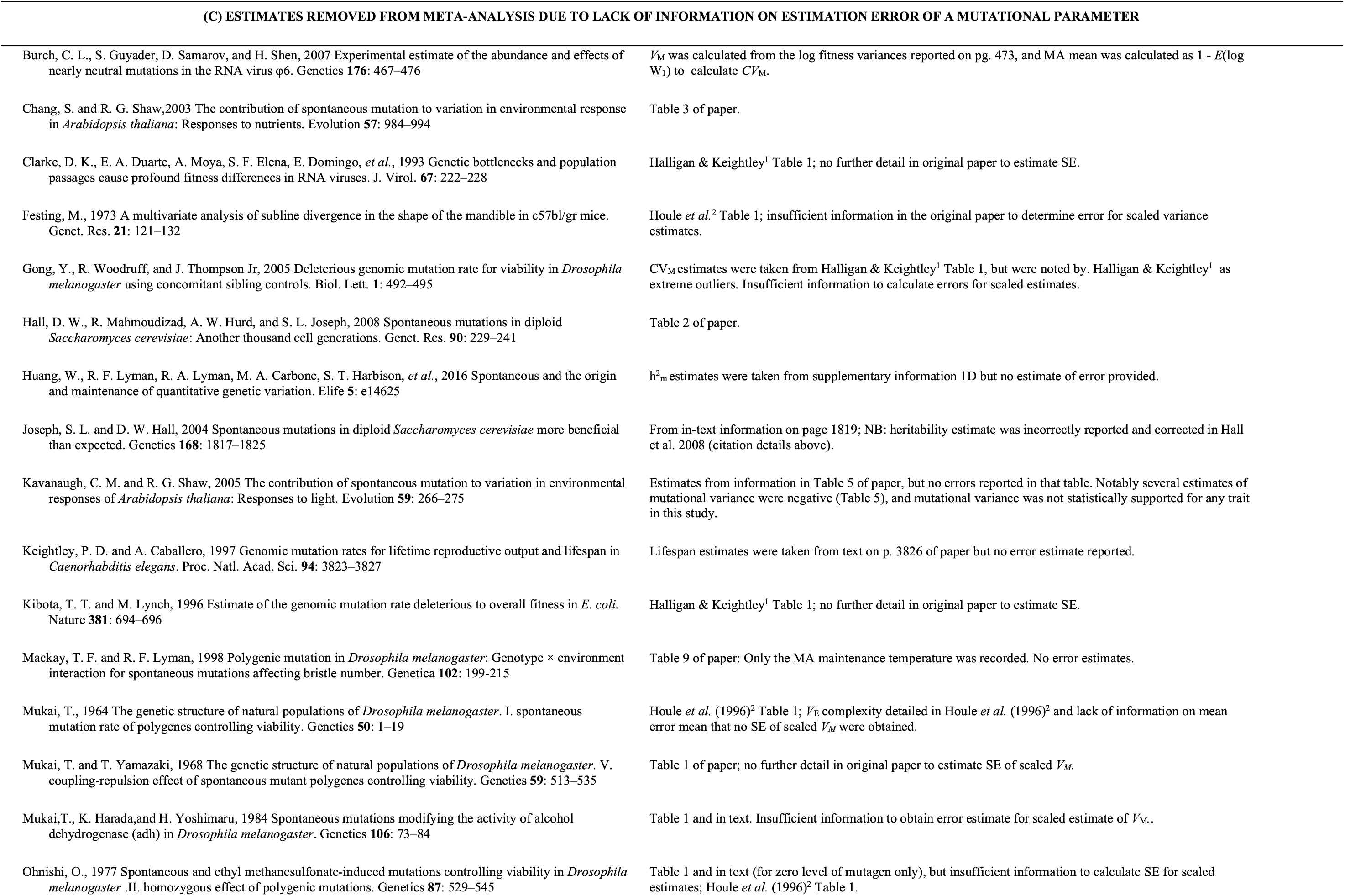
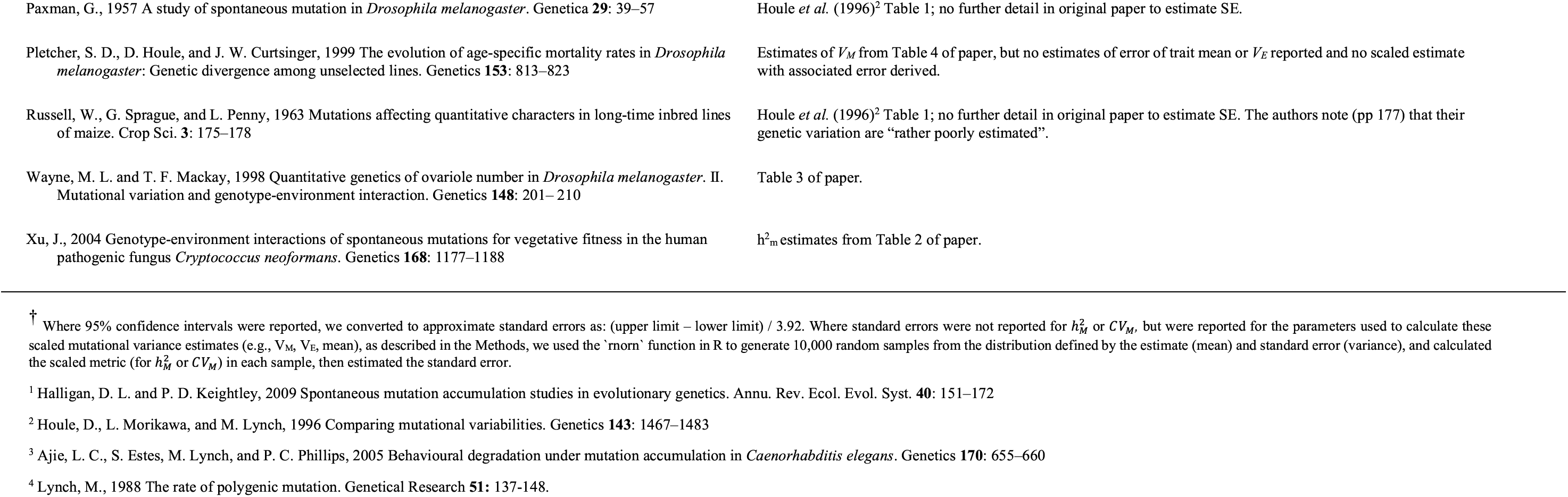
Information on mutational variance estimates obtained from the literature. We detail estimates that were included in the meta-analysis (A) and those that were disqualified (B, C). For each paper, we detail the source of the estimates of scaled mutational variance (mutational heritability, 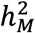 and, coefficient of mutational variance, *CV_M_*) and associated error estimates^†^. Studies (or individual estimates within studies) with scaled estimate (and error) required for the meta-analysis, but which were excluded on based on some additional data criteria are listed (and exclusion justified) in (B). In C, we list papers (estimates) that were excluded due to insufficient information to obtain standard error estimates for the scaled mutational parameters. NB: Some papers appear in (A) and (B) or (C). Only estimates identified in (A) were included in the meta-analysis. Estimates are available at: https://figshare.com/articles/dataset/MA_dserrata_wings_csv/14913051.

| Reference | Description of where estimates were obtained from |
| --- | --- |
| **(A) ESTIMATES USED IN THE META-ANALYSIS** |  |
| Ávila, V. and A. García-Dorado, 2002 The effects of spontaneous mutation on competitive fitness in *Drosophila melanogaster*. J. Evol. Biol. **15**: 561–566 | $CV_M$ from Halligan & Keightley<sup>1</sup> Table 1. |
| Azevedo, R. L., P. D. Keightley, C. Laurén-Määttä, L. L. Vassilieva, M. Lynch, *et al.*, 2002 Spontaneous mutational variation for body size in *Caenorhabditis elegans*. Genetics **162**: 755–765 | Table 1 of paper with scale of mean corrected based on Figure 1. |
| Baer, C. F., F. Shaw, C. Steding, M. Baumgartner, A. Hawkins, *et al.*, 2005 Comparative evolutionary genetics of spontaneous mutations affecting fitness in rhabditid nematodes. Proc. Natl. Acad. Sci. **102**: 5785–5790 | $h_M^2$, $V_M$ and $CV_M$ estimates calculated from information in paper's Table 1 and Supplementary Tables 3, 4 and 5. Error estimated by sampling<sup>†</sup>. |
| Baer, C. F., J. Joyner-Matos, D. Ostrow, V. Grigaltchik, M. P. Salomon, *et al.*, 2010 Rapid decline in fitness of mutation accumulation lines of gonochoristic (outcrossing) *Caenorhabditis nematodes*. Evolution **64**: 3242–3253 | Table 2 of paper; $V_M$ reported as $I = V_M/\bar{x}^2$ and therefore $CV_M$ calculated as: $100*\sqrt{I}$. Confidence intervals from bootstrapping where parameter constrained to be positive. |
| Caballero, A., E. Cusi, C. Garcia, and A. Garcia-Dorado, 2002 Accumulation of deleterious mutations: Additional *Drosophila melanogaster* estimates and a simulation of the effects of selection. Evolution **56**: 1150–1159 | $CV_M$ calculated from in-text information, taking $V_B$ from $\Delta V = V_B / (\text{mean}_c \times F_c)$; where $F_c = 203$ and $\text{mean}_c = 0.625$. Error estimated by sampling<sup>†</sup>. |
| Charlesworth, L., H. Borthwick, C. Bartolomé, and P. Pignatelli, 2004 Estimates of the genomic mutation rate for detrimental alleles in *Drosophila melanogaster*. Genetics **167**: 815–826 | $CV_M$ (and error) estimates taken from Halligan and Keightley<sup>1</sup> Table 1. |
| Clark, A. G., L. Wang, and T. Hulleberg, 1995 Spontaneous mutation rate of modifiers of metabolism in *Drosophila*. Genetics **139**: 767–779 | Trait means were taken from Table 1. $h_m^2$ and $V_M$ estimates taken from Table 4; $V_M$ was multiplied by 2.5 (scale for whole genome, as stated by authors) before calculating $CV_M$. Error in $CV_M$ estimated by sampling<sup>†</sup>. |
| Davies, S. K., A. Leroi, A. Burt, J. G. Bundy, and C. F. Baer, 2016 The mutational structure of metabolism in *Caenorhabditis elegans*. Evolution **70**: 2239–2246 | Table S1 of paper. Note: No estimate of $CV_M$ for citric acid as mean mis-reported in Table S1 and correct value unknown. Confidence intervals from bootstrapping where parameter constrained to be positive. |
| Estes, S., L. C. Ajie, M. Lynch, and P. C. Phillips, 2005 Spontaneous mutational correlations for life-history, morphological and behavioral characters in *Caenorhabditis elegans*. Genetics **170**: 645–653 | We calculated $CV_M$ using $V_M$ (Table 1) and the mean from Table 1 of the companion paper by Ajie *et al.*<sup>3</sup>. $h_m^2$ was from Table 1 of Ajie *et al.*<sup>3</sup> |
| Etienne, V., E. C. Andersen, J. M. Ponciano, D. Blanton, A. Cadavid, *et al.*, 2015 The red death meets the abdominal bristle: polygenic mutation for susceptibility to a bacterial pathogen in *Caenorhabditis elegans*. Evolution **69**: 508–519 | Table 1 of paper. Error in $h_m^2$ estimated by sampling<sup>†</sup>. Confidence intervals from bootstrapping where parameter constrained to be positive. |
| Fernández, J. and C. López-Fanjul, 1996 Spontaneous mutational variances and covariances for fitness-related traits in *Drosophila melanogaster*. Genetics **143**: 829–837 | $V_M$ and $h_m^2$ taken from Table 3 & trait mean from Table 1 of paper; $CV_M$ calculated from those estimates (but no error estimate available). |
| Fry, J. D. and S. L. Heinsohn, 2002 Environment dependence of mutational parameters for viability in *Drosophila melanogaster*. *Genetics* **161**: 1155–1167 | Trait means were from Table 1 and mutational variances were from Table 3; $CV_M$ was calculated from those estimates. Error estimated by sampling<sup>†</sup>. |
| Fry, J. D., P. D. Keightley, S. L. Heinsohn, and S. V. Nuzhdin, 1999 New estimates of the rates and effects of mildly deleterious mutation in *Drosophila melanogaster*. *Proc. Natl. Acad. Sci.* **96**: 574–579 | Trait means and mutational variances were from Fry and Heinsohn (2002)'s Table 1 and Table 3, respectively, where $CV_M$ was calculated from those estimates. |
| García-Dorado, A., J. Fernández, and C. López-Fanjul, 2000 Temporal uniformity of the spontaneous mutational variance of quantitative traits in *Drosophila melanogaster*. *Genet. Res.* **75**: 47–51 | Table 2 of paper. |
| Harada, K., 1995 A quantitative analysis of modifier mutations which occur in mutation accumulation lines in *Drosophila melanogaster*. *Heredity* **75**: 589–598 | $CV_M$ was calculated from information in Table 3 and 4 of paper; $V_M$ was calculated using $4V_e$ (reported) and divided by $2 \times$ the number of generations. Error estimated by sampling<sup>†</sup>. |
| Houle, D., K. A. Hughes, D. K. Hoffmaster, J. Ihara, S. Assimacopoulos, *et al.*, 1994 The effects of spontaneous mutation on quantitative traits. I. Variances and covariances of life history traits. *Genetics* **138**: 773–785 | $V_M$ from Table 4 of paper ($\times 2.5$ to scale to genome-wide as indicated by the authors); data was standardised to mean=1 prior to estimation, and $CV_M = 100 * \sqrt{V_M}$. Error estimated by sampling<sup>†</sup>. |
| Houle, D., and S. V. Nuzhdin, 2004 Mutation accumulation and the effect of copia insertions in *Drosophila melanogaster*. *Genetical Research* **83**: 7–18 | Table 3 of paper (for IVE-39 sublines only, constrained intercept), where $CV_M = 100 * \sqrt{\Delta V}$ as described in text of paper on page 9. Error estimated by sampling<sup>†</sup>. |
| Joyner-Matos, J., A. Upadhyay, M. P. Salomon, V. Grigaltchik, and C. F. Baer, 2009 Genetic (co)variation for life span in rhabditid nematodes: Role of mutation, selection, and history. *J. Gerontol. A Biol. Sci. Med. Sci.* **64**: 1134–1145 | Table 1 and 2 of paper. |
| Keightley, P. D. and A. Caballero, 1997 Genomic mutation rates for lifetime reproductive output and lifespan in *Caenorhabditis elegans*. *Proc. Natl. Acad. Sci.* **94**: 3823–3827 | $h_m^2$ estimate (and SE) provided in text on pg 3825. Estimates for productivity determined from among-line variances and trait mean taken from Figure 2 using the R package *metaDigitise*. Error in $CV_M$ estimated by sampling<sup>†</sup>. |
| Latimer, C. A. L., K. McGuigan, R. S. Wilson, M. W. Blows, and S. F. Chenoweth, 2014 The contribution of spontaneous mutations to thermal sensitivity curve variation in *Drosophila serrata*. *Evolution* **68**: 1824–1837 | Table 1 of paper |
| López, M. A. and C. López-Fanjul, 1993 Spontaneous mutation for a quantitative trait in *Drosophila melanogaster*. II. Distribution of mutant effects on the trait and fitness. *Genet. Res.* **61**: 117–126 | We used estimates from B (full-sib mating) lines only, where trait means were from figure 1 (males=30, females = 36.8). Among-line variance and $h_m^2$ for bristle number from Table 6, $CV_M$ calculated from the provided information, but no error estimate available. |
| Lynch, M., 1985 Spontaneous mutations for life-history characters in an obligate parthenogen. *Evolution* **39**: 804–818 | $h_m^2$ from Table 1 and $CV_M$ calculated from information in Table 1 (but no SE provided for mean and hence unavailable for $CV_M$). Error estimated by sampling<sup>†</sup>. |
| Lynch, M., L. Latta, J. Hicks, and M. Giorgianni, 1998 Mutation, selection, and the maintenance of life-history variation in a natural population. *Evolution* **52**: 727–733 | Estimates of $h_m^2$, $V_M$, $V_E$ and trait mean (MA mean calculated for the final generation as: $z_0 + R_m * 30$) from Table 1 of paper and $CV_M$ calculated from this information. Error in $CV_M$ estimated by sampling<sup>†</sup>; $z_0$ and $R_m$ were sampled to allow the mean to be determined per sample. |
| Mackay, T. F., R. F. Lyman, M. S. Jackson, C. Terzian and W. G. Hill, 1992 Polygenic mutation in *Drosophila melanogaster*: Estimates from divergence among inbred strains. *Evolution* **46**: 300–316 | Table 3 of paper, assuming $N = 0.7N_e$. Estimates available for per generation (Table 1), but SE not reported. |
| Martorell, C., M. Toro, and C. Gallego, 1998 Spontaneous mutation for life-history traits in *Drosophila melanogaster*. *Genetica* **102/103**: 315–324 | Table 2 of paper. |
| McGuigan, K., D. Petfield, and M. W. Blows, 2011 Reducing mutation load through sexual selection on males. *Evolution* **65**: 2816–2829 | $h_m^2$ was taken from text. $CV_M$ was calculated by determining $V_M$ as (mean ($V_E$) × $h_m^2$), where $V_E$ and MA means were taken from Tables S1 and S2 of paper for the Control MA population. Error estimated by sampling<sup>†</sup>. |
| Mukai, T., S. I. Chigusa, J. F. Crow, and L. E. Mettler, 1972 Mutation rate and dominance of genes affecting viability in *Drosophila melanogaster*. *Genetics* **72**: 335–355 | Among-line and mean estimates from Tables 1, 2 and 3 of paper for Generation 40 only were used to calculate $V_M$ and $CV_M$. |
| Ostrow, D., N. Phillips, A. Avalos, D. Blanton, A. Boggs, *et al.*, 2007 Mutational bias for body size in Rhabditid nematodes. *Genetics* **176**: 1653–1661 | Trait mean from Table 1 and $CV_M$ (MA mean), $V_M$ and $h_m^2$ from Table 2 of paper. While log transformation had little impact on scaled estimates, $V_M$ and $V_E$ were extreme on the raw scale, while mean of the log-transformed data was negative. |
| Pletcher, S. D., D. Houle, and J. W. Curtsinger, 1998 Age-specific properties of spontaneous mutations affecting mortality in *Drosophila melanogaster*. *Genetics* **148**: 287–303 | $h_m^2$, $V_M$ and mean from Table 5 of paper and $CV_M$ estimated from that information. Error estimated by sampling<sup>†</sup>. |
| Roles, A. J., M. T. Rutter, I. Dworkin, C. L. Fenster, and J. K. Conner, 2016 Field measurements of genotype by environment interaction for fitness caused by spontaneous mutations in *Arabidopsis thaliana*. *Evolution* **70**: 1039–1050 | Table 2 of paper. We included $CV_M$ for fruit number and total fitness however authors have calculated $CV_M$ as $100(e^{\sqrt{V_M}}-1)$ as per Poisson values. Parameter confidence intervals from MCMCglmm, and constrained to be positive. |
| Rutter, M. T., F. H. Shaw, and C. L. Fenster, 2010 Spontaneous mutation parameters for *Arabidopsis thaliana* measured in the wild. *Evolution* **64**: 1825–35 | Table 2 of paper. |
| Santiago, E., J. Albornoz, A. Domínguez, M. Toro, and C. López-Fanjul, 1992 The distribution of spontaneous mutations on quantitative traits and fitness in *Drosophila melanogaster*. *Genetics* **132**: 771–781 | $h_m^2$ and $V_E$ from Table 4 for full-sib mating lines only (B lines; all lines). Trait means were taken from Figure 3 (sternopleural bristles = 21.7, wing length = 1210 and wing width = 465). $CV_M$ estimated from that information but error estimates not available. |
| Schoen, D. J., 2005 Deleterious mutation in related species of the plant genus *Amsinckia* with contrasting mating systems. *Evolution* **59**: 2370–2377 | Trait means and mutational variances were taken from Table 1 and 3 for flower number and weight respectively. $V_M$, $h_m^2$, and $CV_M$ were estimated from that information, where $V_L$ for MA was corrected for initial (control variance) if that was non-zero; error estimates were similar between control and mutant and MA SE were used to approximate error for meta-analysis. |
| Schultz, S. T., M. Lynch, and J. H. Willis, 1999 Spontaneous deleterious mutation in *Arabidopsis thaliana*. *Proc. Natl. Acad. Sci.* **96**: 11393–11398 | From Table 3; $CV_M$ not available as calculated from among-line variance, not the per-generation $V_M$. |
| Shaw, R. G., D. L. Byers, and E. Darnø, 2000 Spontaneous mutational effects on reproductive traits of *Arabidopsis thaliana*. *Genetics* **155**: 369–78 | Estimates of mean and among-line and environmental variances taken from Table 1 of paper (generation 17), and estimates of $V_M$, $h_m^2$, and $CV_M$ from Table 2. To estimate error for $h_m^2$, and $CV_M$, the among-line and mean estimates for generation 17 were used, along with $V_M$ from the joint analysis. Error estimated by sampling<sup>†</sup> |
| Vassilieva, L. L., A. M. Hook, and M. Lynch, 2000 The fitness effects of spontaneous mutations in *Caenorhabditis elegans*. *Evolution* **54**: 1234–1246 | $h_m^2$ was taken from Table 2; trait mean (intercept) from Table 1 and $V_M$ (slope/2) from Table 2 were used to calculate $CV_M$. Error in $CV_M$ estimated by sampling<sup>†</sup> following the approach detailed for Lynch *et al.* (1998) above. For one trait (generation time) the reported standard error of the $V_M$ was 0, precluding calculation of errors. |
| Vassilieva, L. L. and M. Lynch, 1999 The rate of spontaneous mutation for life-history traits in *Caenorhabditis elegans*. *Genetics* **151**: 119–129 | $h_m^2$ and $V_M$ estimates were taken from Table 1 of paper. Error in $CV_M$ estimated by sampling<sup>†</sup> following the approach detailed for Lynch *et al.* (1998) above. For one trait (generation time) the estimated mean was ~0, resulting in nonsensical $CV_M$ and hence this was excluded. |
| **(B) ESTIMATES REMOVED FROM META-ANALYSIS DUE TO ISSUES WITH MUTATIONAL PARAMETER ESTIMATES** |  |
| Baer, C. F., J. Joyner-Matos, D. Ostrow, V. Grigaltchik, M. P. Salomon, *et al.*, 2010 Rapid decline in fitness of mutation accumulation lines of gonochoristic (outcrossing) *Caenorhabditis nematodes*. *Evolution* **64**: 3242–3253 | As in (A); Excluded *Caenorhabditis brenneri* strain PB2801 $h_m^2$ and strain PB2802 $CV_M$ generation 100 estimates as they were outliers (>3.5 standard deviations from the mean and >5 times the inter-quartile range from the median) and . |
| Bailey, D. W., 1959 Rates of subline divergence in highly inbred strains of mice. *J. Hered* **50**: 26–30. | From Table 1, and in-text information on number of generations; $h_m^2$ calculated as per Lynch 1988<sup>4</sup>. Excluded due to low representation of vertebrate taxa. |
| Casellas, J. and J. F. Medrano, 2008 Within-generation mutation variance for litter size in inbred mice. *Genetics* **179**: 2147–55 | MA litter mean was taken for generation 46 in Table 1. The $V_M$ and $h_m^2$ estimates were taken from text and Table 3. $CV_M$ was calculated from reported information. Excluded due to low representation of vertebrate taxa. |
| Clark, A. G., L. Wang, and T. Hulleberg, 1995 Spontaneous mutation rate of modifiers of metabolism in *Drosophila*. *Genetics* **139**: 767–779 | As in (A). ADH $h_m^2$ excluded as it was an extreme outlier (>4 standard deviations from the mean and >5 times the inter-quartile range from the median). |
| Downie, D., 2003 Effects of short-term spontaneous mutation accumulation for life history traits in grape phylloxera, *Daktulosphaira vitifoliae*. *Genetica* **119**: 237–251 | $V_M$ and $h_m^2$ were taken from Table 1. Line means were estimated from figure 3 and used to calculate $CV_M$ (but not error). Excluded due to low representation of non-Drosophila insect taxa. |
| Farhadifar, R., J. M. Ponciano, E. C. Andersen, D. J. Needleman, and C. F. Baer, 2016 Mutation is a sufficient and robust predictor of genetic variation for mitotic spindle traits in *Caenorhabditis elegans*. *Genetics* **203**: 1859–1870 | Trait means from Table 1 of paper, and variances from Table S4. One estimate (for posterior centrosome oscillation frequency) was removed as the estimate for $V_M$ was negative. Excluded due to low representation of trait type (mitotic cell division). |
| Harada, K., 1995 A quantitative analysis of modifier mutations which occur in mutation accumulation lines in *Drosophila melanogaster*. *Heredity* **75**: 589–598 | As in (A); Could not calculate $CV_M$ (negative mean) for one metabolite (Isocitrate-dehydrogenase activity) which was therefore excluded. Alpha-analyse was excluded as the estimated SE of $CV_M$ was considerably larger than the estimate itself. |
| Houle, D., K. A. Hughes, D. K. Hoffmaster, J. Ihara, S. Assimakopoulos, *et al.*, 1994 The effects of spontaneous mutation on quantitative traits. I. Variances and covariances of life history traits. *Genetics* **138**: 773–785 | As in (A). Early male mating success excluded as $CV_M$ estimate was negative; late mating $CV_M$ excluded due extreme value ($> 4$ standard deviations from the mean and $>5$ times the inter-quartile range from the median) |
| Joyner-Matos, J., L. C. Bean, H. L. Richardson, T. Sammeli, and C. F. Baer, 2011 No evidence of elevated germline mutation accumulation under oxidative stress in *Caenorhabditis elegans*. *Genetics* **189**: 1439–1447 | Information in Table 1 of paper used to calculate $CV_M = 100 * \sqrt{V_M}$, where $V_M$ (and $V_L$) were reported on intensity of selection scale, as described on pg 1441 of paper. We sampled from the distribution of parameters used to calculate $V_M$, but the standard error of the sample was orders of magnitude larger than the estimate. Therefore, this study was excluded. |
| Latta, L. C., M. Peacock, D. J. Civitello, J. L. Dudyche, J. M. Meik, *et al.*, 2015 The phenotypic effects of spontaneous mutations in different environments. *Am. Nat.* **185**: 243–252 | Among-line variance (designated by authors as $\Delta V$) provided in Table S1 and used to calculated using $V_M = \Delta V / (2 * 96)$, where 96 was the number of generations of MA reported by the authors on pg 245 of paper. $V_E$ and $\bar{X}$ were provided in Table S1 and, along with the calculated $V_M$, used to calculate $h_m^2$ and $CV_M$. However, the reported confidence intervals for $\Delta V$ are extremely large (infinite for several traits), and the standard errors calculated are always larger than the parameter estimate, and all estimates from this study were excluded. Notably, this study included only five MA lines, and poor quality estimates could be expected. |
| López, M. A. and C. López-Fanjul, 1993 Spontaneous mutation for a quantitative trait in *Drosophila melanogaster*. II. Distribution of mutant effects on the trait and fitness. *Genet. Res.* **61**: 117–126 |  |
| Lynch, M., L. Latta, J. Hicks, and M. Giorgianni, 1998 Mutation, selection, and the maintenance of life-history variation in a natural population. *Evolution* **52**: 727–733. | As in (A). Second clutch size excluded as it was an extreme outlier ($>7$ standard deviations from the mean and $>5$ times the inter-quartile range from the median). Size at birth was excluded because calculation of $h_m^2$ from the data in Table 1 returned an estimate several orders of magnitude larger than reported, suggesting some incorrect parameter values reported. |
| Morgan, A. D., R. W. Ness, P. D. Keightley, and N. Colegrave, 2014 Spontaneous mutation accumulation in multiple strains of the green alga, *Chlamydomonas reinhardtii*. *Evolution* **68**: 2589–2602 | $CV_M$ estimates were taken from Table 3 (second assay only, as one strain had missing data for the first assay), but no SE reported. To estimate SE it was necessary to use $V_L$ estimates for MA and ancestor from Table 3 (and reported CI), the generation number estimates from Table 2, and estimates of trait mean (CI) from Table 4. Excluded due to low representation of micro-organisms. |
| Pannebakker, L. A., D. L. Halligan, K. T. Reynolds, G. A. Ballantyne, D. M. Shuker, *et al.*, 2008 Effects of spontaneous mutation accumulation on sex ratio traits in a parasitoid wasp. *Evolution* **62**: 1921–1935 | MA mean and line numbers taken from Table 2 of paper. Variance estimates were taken from Table 3, where $CV_M$ has been converted to a percentage scale. Excluded due to low representation of non-Drosophila insect taxa. |
| **(C) ESTIMATES REMOVED FROM META-ANALYSIS DUE TO LACK OF INFORMATION ON ESTIMATION ERROR OF A MUTATIONAL PARAMETER** |  |
| Burch, C. L., S. Guyader, D. Samarov, and H. Shen, 2007 Experimental estimate of the abundance and effects of nearly neutral mutations in the RNA virus $\phi 6$. *Genetics* **176**: 467–476 | $V_M$ was calculated from the log fitness variances reported on pg. 473, and MA mean was calculated as $1 - E(\log W_i)$ to calculate $CV_M$. |
| Chang, S. and R. G. Shaw, 2003 The contribution of spontaneous mutation to variation in environmental response in *Arabidopsis thaliana*: Responses to nutrients. *Evolution* **57**: 984–994 | Table 3 of paper. |
| Clarke, D. K., E. A. Duarte, A. Moya, S. F. Elena, E. Domingo, *et al.*, 1993 Genetic bottlenecks and population passages cause profound fitness differences in RNA viruses. *J. Virol.* **67**: 222–228 | Halligan & Keightley<sup>1</sup> Table 1; no further detail in original paper to estimate SE. |
| Festing, M., 1973 A multivariate analysis of subline divergence in the shape of the mandible in c57bl/gr mice. *Genet. Res.* **21**: 121–132 | Houle *et al.*<sup>2</sup> Table 1; insufficient information in the original paper to determine error for scaled variance estimates. |
| Gong, Y., R. Woodruff, and J. Thompson Jr, 2005 Deleterious genomic mutation rate for viability in *Drosophila melanogaster* using concomitant sibling controls. *Biol. Lett.* **1**: 492–495 | $CV_M$ estimates were taken from Halligan & Keightley<sup>1</sup> Table 1, but were noted by. Halligan & Keightley<sup>1</sup> as extreme outliers. Insufficient information to calculate errors for scaled estimates. |
| Hall, D. W., R. Mahmoudizad, A. W. Hurd, and S. L. Joseph, 2008 Spontaneous mutations in diploid *Saccharomyces cerevisiae*: Another thousand cell generations. *Genet. Res.* **90**: 229–241 | Table 2 of paper. |
| Huang, W., R. F. Lyman, R. A. Lyman, M. A. Carbone, S. T. Harbison, *et al.*, 2016 Spontaneous and the origin and maintenance of quantitative genetic variation. *Elife* **5**: e14625 | $h^2_m$ estimates were taken from supplementary information 1D but no estimate of error provided. |
| Joseph, S. L. and D. W. Hall, 2004 Spontaneous mutations in diploid *Saccharomyces cerevisiae* more beneficial than expected. *Genetics* **168**: 1817–1825 | From in-text information on page 1819; NB: heritability estimate was incorrectly reported and corrected in Hall *et al.* 2008 (citation details above). |
| Kavanaugh, C. M. and R. G. Shaw, 2005 The contribution of spontaneous mutation to variation in environmental responses of *Arabidopsis thaliana*: Responses to light. *Evolution* **59**: 266–275 | Estimates from information in Table 5 of paper, but no errors reported in that table. Notably several estimates of mutational variance were negative (Table 5), and mutational variance was not statistically supported for any trait in this study. |
| Keightley, P. D. and A. Caballero, 1997 Genomic mutation rates for lifetime reproductive output and lifespan in *Caenorhabditis elegans*. *Proc. Natl. Acad. Sci.* **94**: 3823–3827 | Lifespan estimates were taken from text on p. 3826 of paper but no error estimate reported. |
| Kibota, T. T. and M. Lynch, 1996 Estimate of the genomic mutation rate deleterious to overall fitness in *E. coli*. *Nature* **381**: 694–696 | Halligan & Keightley<sup>1</sup> Table 1; no further detail in original paper to estimate SE. |
| Mackay, T. F. and R. F. Lyman, 1998 Polygenic mutation in *Drosophila melanogaster*: Genotype × environment interaction for spontaneous mutations affecting bristle number. *Genetica* **102**: 199–215 | Table 9 of paper: Only the MA maintenance temperature was recorded. No error estimates. |
| Mukai, T., 1964 The genetic structure of natural populations of *Drosophila melanogaster*. I. spontaneous mutation rate of polygenes controlling viability. *Genetics* **50**: 1–19 | Houle *et al.* (1996)<sup>2</sup> Table 1; $V_E$ complexity detailed in Houle *et al.* (1996)<sup>2</sup> and lack of information on mean error mean that no SE of scaled $V_M$ were obtained. |
| Mukai, T. and T. Yamazaki, 1968 The genetic structure of natural populations of *Drosophila melanogaster*. V. coupling-repulsion effect of spontaneous mutant polygenes controlling viability. *Genetics* **59**: 513–535 | Table 1 of paper; no further detail in original paper to estimate SE of scaled $V_M$. |
| Mukai, T., K. Harada, and H. Yoshimaru, 1984 Spontaneous mutations modifying the activity of alcohol dehydrogenase (adh) in *Drosophila melanogaster*. *Genetics* **106**: 73–84 | Table 1 and in text. Insufficient information to obtain error estimate for scaled estimate of $V_M$. |
| Ohnishi, O., 1977 Spontaneous and ethyl methanesulfonate-induced mutations controlling viability in *Drosophila melanogaster*. II. homozygous effect of polygenic mutations. *Genetics* **87**: 529–545 | Table 1 and in text (for zero level of mutagen only), but insufficient information to calculate SE for scaled estimates; Houle *et al.* (1996)<sup>2</sup> Table 1. |
| Paxman, G., 1957 A study of spontaneous mutation in *Drosophila melanogaster*. *Genetica* **29**: 39–57 | Houle *et al.* (1996)<sup>2</sup> Table 1; no further detail in original paper to estimate SE. |
| Pletcher, S. D., D. Houle, and J. W. Curtsinger, 1999 The evolution of age-specific mortality rates in *Drosophila melanogaster*: Genetic divergence among unselected lines. *Genetics* **153**: 813–823 | Estimates of $V_M$ from Table 4 of paper, but no estimates of error of trait mean or $V_E$ reported and no scaled estimate with associated error derived. |
| Russell, W., G. Sprague, and L. Penny, 1963 Mutations affecting quantitative characters in long-time inbred lines of maize. *Crop Sci.* **3**: 175–178 | Houle *et al.* (1996)<sup>2</sup> Table 1; no further detail in original paper to estimate SE. The authors note (pp 177) that their genetic variation are “rather poorly estimated”. |
| Wayne, M. L. and T. F. Mackay, 1998 Quantitative genetics of ovariole number in *Drosophila melanogaster*. II. Mutational variation and genotype-environment interaction. *Genetics* **148**: 201–210 | Table 3 of paper. |
| Xu, J., 2004 Genotype-environment interactions of spontaneous mutations for vegetative fitness in the human pathogenic fungus *Cryptococcus neoformans*. *Genetics* **168**: 1177–1188 | $h^2_m$ estimates from Table 2 of paper. |

<sup>†</sup> Where 95% confidence intervals were reported, we converted to approximate standard errors as: (upper limit – lower limit) / 3.92. Where standard errors were not reported for $h^2_M$ or $CV_M$ , but were reported for the parameters used to calculate these scaled mutational variance estimates (e.g., $V_M$ , $V_E$ , mean), as described in the Methods, we used the ‘rnorm’ function in R to generate 10,000 random samples from the distribution defined by the estimate (mean) and standard error (variance), and calculated the scaled metric (for $h^2_M$ or $CV_M$ ) in each sample, then estimated the standard error.
<sup>1</sup> Halligan, D. L. and P. D. Keightley, 2009 Spontaneous mutation accumulation studies in evolutionary genetics. *Annu. Rev. Ecol. Evol. Syst.* **40**: 151–172
<sup>2</sup> Houle, D., L. Morikawa, and M. Lynch, 1996 Comparing mutational variabilities. *Genetics* **143**: 1467–1483
<sup>3</sup> Ajie, L. C., S. Estes, M. Lynch, and P. C. Phillips, 2005 Behavioural degradation under mutation accumulation in *Caenorhabditis elegans*. *Genetics* **170**: 655–660
<sup>4</sup> Lynch, M., 1988 The rate of polygenic mutation. *Genetical Research* **51**: 137–148.

**Table S2.** Trait types included in the broad categories and within the more narrowly defined life-history trait categories. The trait description is a list of traits included within each category.

| Broad Category | Narrow Category | Trait Description |
| --- | --- | --- |
| Life-history | Fitness | Fitness; Intrinsic rate of increase; Competitive mating success; Rate of convergence to the stable age distribution. |
|  | Miscellaneous | Longevity, Lifespan, Male mating success, Age at maturity, Age at reproduction, Growth rate, Instar duration, The reciprocal of the mean age at reproduction, Time to maturity. |
|  | Productivity | Clutch size, Fecundity, Litter size, Number of flowers, Seed count, Productivity, Reproductive mass |
|  | Survival | Survival, Age-specific survivorship or viability, Germination, Pathogen susceptibility. |
| Morphology | - | Abdominal and sternopleural bristles; Body length, width or size; Limb/Organ/Spine length or width, Weight. |
| Physiology | - | Enzyme activity, Metabolites, Locomotor activity, Velocity. |

**Table S3.** Mutation accumulation (MA) lines shared across studies. The first published description of a set of MA lines is indicated in bold; the following non-bolded entries represent subsequent investigations of the same set of MA lines. See Table S1 for citation details.

| Species name | Reference | Generation | Trait Description |
| --- | --- | --- | --- |
| *Arabidopsis thaliana* | **Shaw *et al.* (2000)** | 17 | Fecundity, Reproductive mass |
|  | Rutter *et al.* (2010) | 25 | Fecundity |
|  | Roles *et al.* (2016) | 25 | Fecundity, Germination, Age-specific survivorship, Fitness |
| *Caenorhabditis briggsae* | **Baer *et al.* (2005)** | 200 | Productivity |
|  | Ostrow *et al.* (2007) | 200 | Body size |
|  | Joyner-Matos *et al.* (2009) | 250 | Life span |
| *Caenorhabditis elegans* | **Baer *et al.* (2005)** | 200 | Productivity |
|  | Ostrow *et al.* (2007) | 200 | Body size |
|  | Joyner-Matos *et al.* (2009) | 250 | Life span |
|  | Etienne *et al.* (2015) | 250 | Pathogen susceptibility |
|  | Davies *et al.* (2016) | 250 | Metabolites |
|  | **Keightley & Caballero (1997)** | 60 | Lifespan, Productivity, Population growth |
|  | Azevedo *et al.* (2002) | 60 | Body size |
|  | **Vassilieva *et al.* (1999)** | 49.4 | Longevity, Productivity, Survivorship, Population growth |
|  | Vassilieva *et al.* (2000) | 214 | Longevity, Productivity, Survivorship, Population growth |
|  | Azevedo *et al.* (2002) | 152 | Body size |
|  | Estes *et al.* and Ajie *et al.* (2005) | 370 | Population growth parameter, Body width Locomotor activity, Productivity, Velocity |
| *Drosophila melanogaster* | **Houle *et al.* (1994)** | 44 | Fecundity, Fitness, Longevity, Male mating, Productivity |
|  | Pletcher *et al.* (1998) | 19 | Female longevity, Male longevity |
|  | **Santiago *et al.* (1992)** | 46 | Sternopleural bristles, Wing dimensions |
|  | Fernandez & Lopez-Fanjul (1996) | 106 | Viability, Fecundity |
|  | Martorell *et al.* (1998) | 82 | Productivity, Longevity |
|  | Garcia-Dorado *et al.* (2000) | 170 | Wing length |
|  |  | 204 | Abdominal bristles, Sternopleural bristles |
|  |  | 209 | Viability |
|  | Caballero *et al.* (2002) | 210 | Viability |
|  | Avila & Garca-Dorado (2002) | 288 | Fitness |
| *Drosophila serrata* | **McGuigan *et al.* (2011a)** | 23 | Mating Success |
|  |  | 26 | Productivity |
|  | Latimer *et al.* (2014) | 32 | Locomotor activity |
| *Oscheius myriophila* | **Baer *et al.* (2005)** | 200 | Productivity |
|  | Ostrow *et al.* (2007) | 200 | Body size |

**Table S4.**
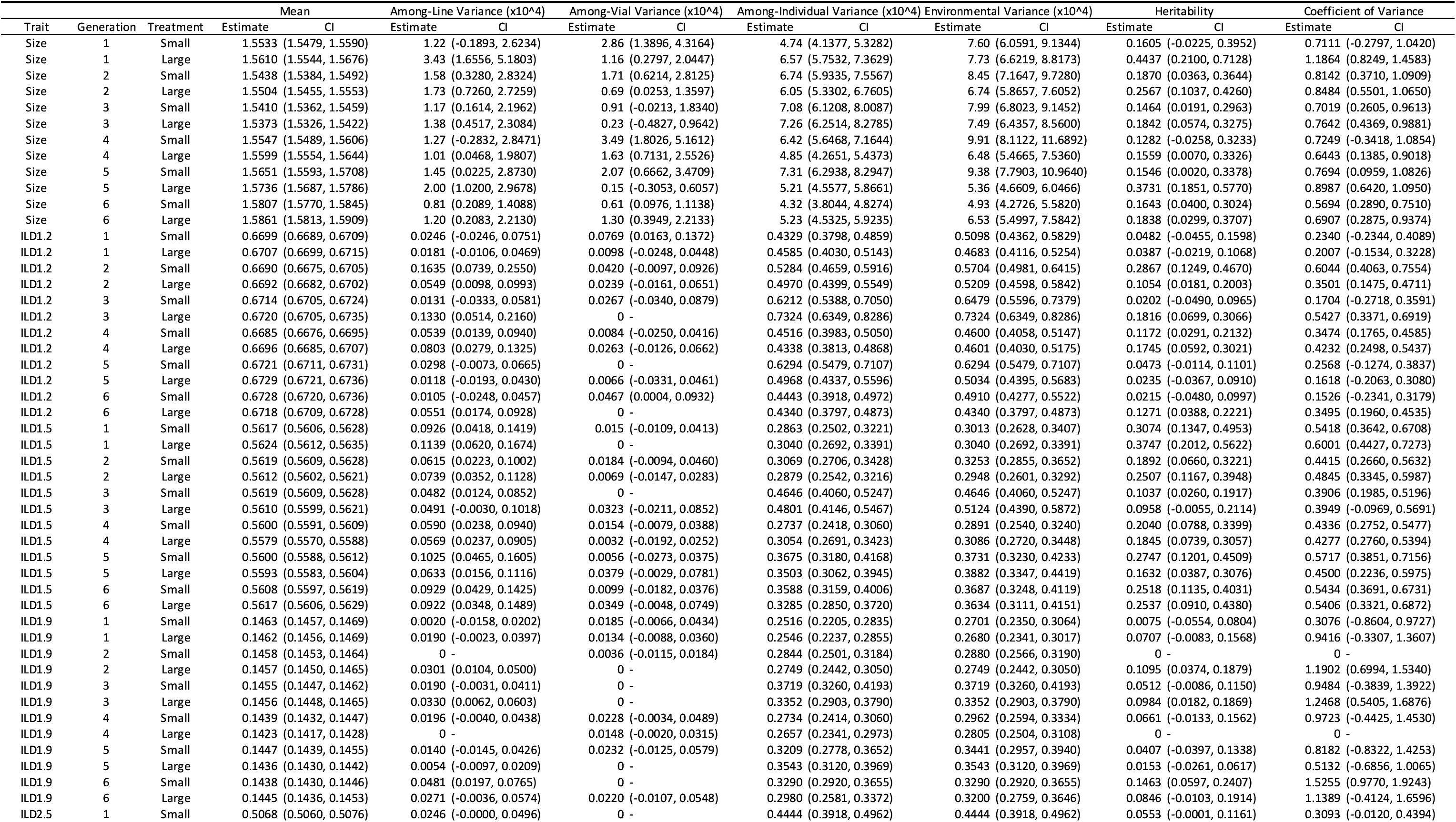
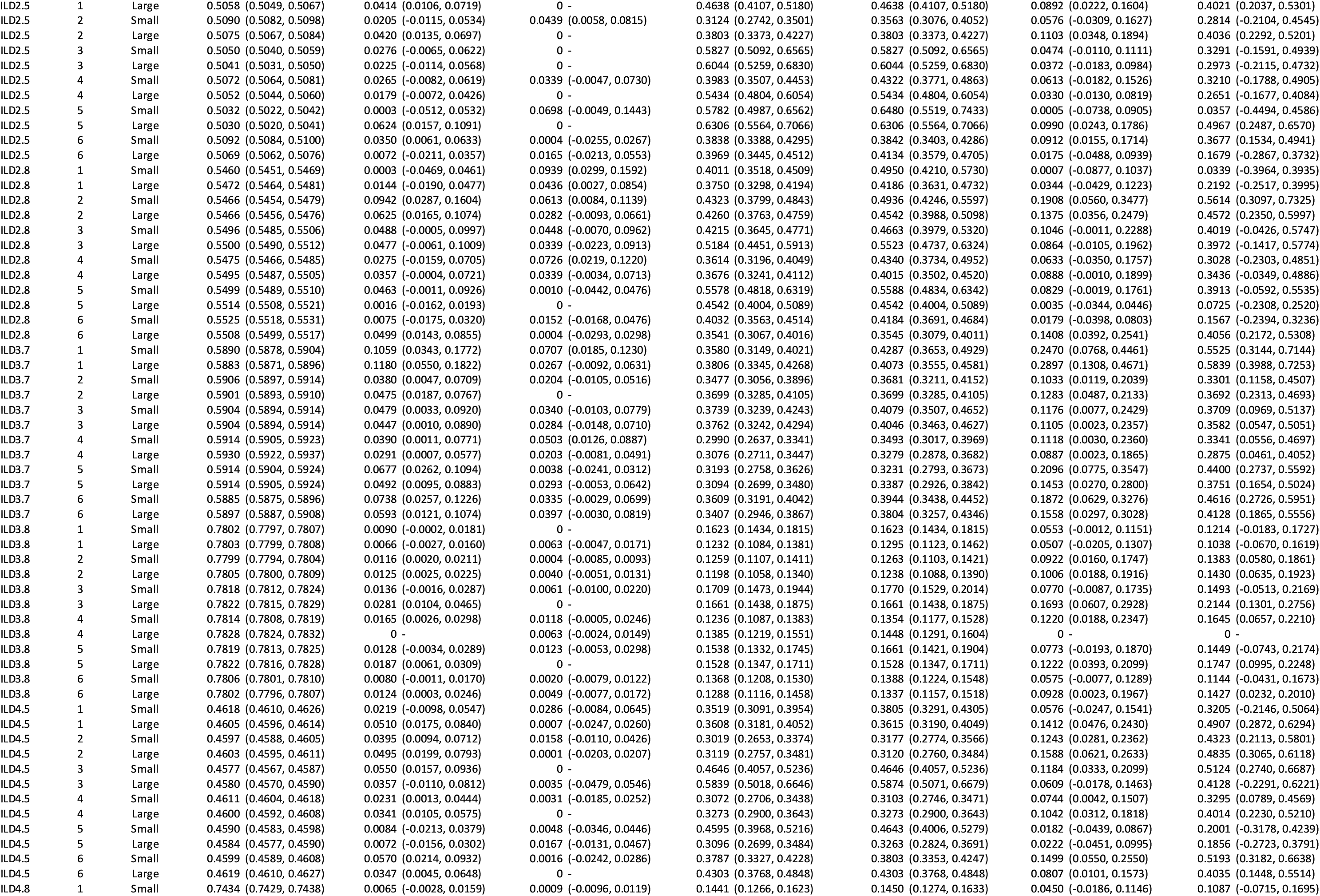
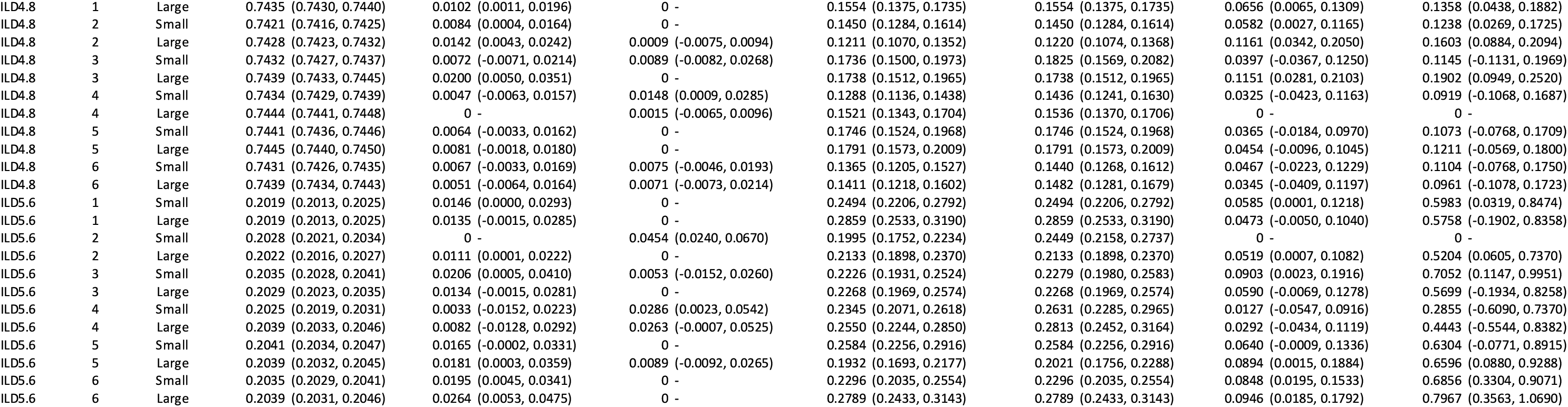
Parameter estimates from analyses of data for each of the 11 traits in each generation for each treatment in the Drosophila serrata experiment. Model (2) was fit to data for each trait in each generation for each population size treatment. The least squares trait mean was estimated, and the phenotypic variance was partitioned to among-line (due to mutation), among the two replicate rearing vials nested within each line, and to the residual (among individuals within each vial) variances. We estimated the environmental variance as the sum of among-vial and among-individual (residual) variances. We placed the estimates of among-line variance on both a heritability (among-line variance / environmental variance) and coefficient of variance (100* √Among-line / Mean) scales but note that these should not be interpreted as mutational heritability or coefficient of variance because they are calculated from the total accumulated mutational variance (among-line variance) not the per generation rate at which that accrued. For each parameter, confidence intervals were estimated by drawing 10,000 random samples from a normal distribution with a mean equal to the REML model estimate and variance equal to either the inverse of the Fisher information matrix (for variance parameters) or the standard deviation (for the trait mean), as described in the Methods. From the 10,000 samples, 95% (Mean) or 90% (all other parameters, which are constrained to be positive) confidence intervals (CI) are reported. The environmental variance, heritability and coefficient of variance) were calculated for each of the 10,000 samples to estimate the CI of these calculated parameters. Statistical support for non-zero estimates was inferred where the lower CI did not include zero. Where REML parameter estimates were zero, no CI were estimated.

**Table S5.** Results of regressing the *D. serrata* among-line variance estimates on either the squared trait mean or on the environmental variance. For each trait (defined in Figure 2B) the slope, *R*^2^, *P*-value and 5% FDR adjusted *P*-value are reported. Only one relationship (ILD3.7 on mean, in bold) was significant at *P* < 0.05, but not following FDR multiple test correction. The analysis was conducted on the squared trait mean, rather than the square root of *V_L_* (i.e., matching evolvability *I,* not *CV*) to allow visualization of both scalar relationships on the same graph (Figure 6).

| Trait | Relationship | Slope (Beta) | $R^2$ | $P$-value | FDR $P$ |
| --- | --- | --- | --- | --- | --- |
| Size | Mean<sup>2</sup> | -0.0001 | 0.2% | 0.8952 | 0.9440 |
| | $V_E$ | 0.0428 | 0.9% | 0.7735 | 0.9440 |
| ILD1.2 | Mean<sup>2</sup> | -0.0009 | 15.5% | 0.2052 | 0.8333 |
| | $V_E$ | 0.1842 | 11.5% | 0.2810 | 0.8333 |
| ILD1.5 | Mean<sup>2</sup> | 0.0005 | 10.5% | 0.3030 | 0.8333 |
| | $V_E$ | -0.1296 | 16.8% | 0.1857 | 0.8333 |
| ILD1.9 | Mean<sup>2</sup> | 0.0002 | 0.2% | 0.8836 | 0.9440 |
| | $V_E$ | 0.1050 | 6.4% | 0.4278 | 0.8766 |
| ILD2.5 | Mean<sup>2</sup> | 0.0000 | 0.1% | 0.9440 | 0.9440 |
| | $V_E$ | -0.0043 | 0.1% | 0.9318 | 0.9440 |
| ILD2.8 | Mean<sup>2</sup> | -0.0003 | 7.2% | 0.3980 | 0.8766 |
| | $V_E$ | 0.0962 | 4.3% | 0.5180 | 0.8766 |
| ILD3.7 | Mean<sup>2</sup> | -0.0013 | 59.5% | **0.0033** | 0.0726 |
| | $V_E$ | 0.4493 | 32.4% | 0.0533 | 0.5863 |
| ILD3.8 | Mean<sup>2</sup> | 0.0001 | 4.7% | 0.5006 | 0.8766 |
| | $V_E$ | 0.1302 | 11.5% | 0.2812 | 0.8333 |
| ILD4.5 | Mean<sup>2</sup> | <0.0000 | 0.1% | 0.9357 | 0.9440 |
| | $V_E$ | 0.0054 | 0.1% | 0.9342 | 0.9440 |
| ILD4.8 | Mean<sup>2</sup> | -0.0001 | 5.1% | 0.4808 | 0.8766 |
| | $V_E$ | 0.0179 | 0.4% | 0.8412 | 0.9440 |
| ILD5.6 | Mean<sup>2</sup> | 0.0009 | 17.6% | 0.1741 | 0.8333 |
| | $V_E$ | -0.0273 | 1.0% | 0.7533 | 0.9440 |

**Table S6.**
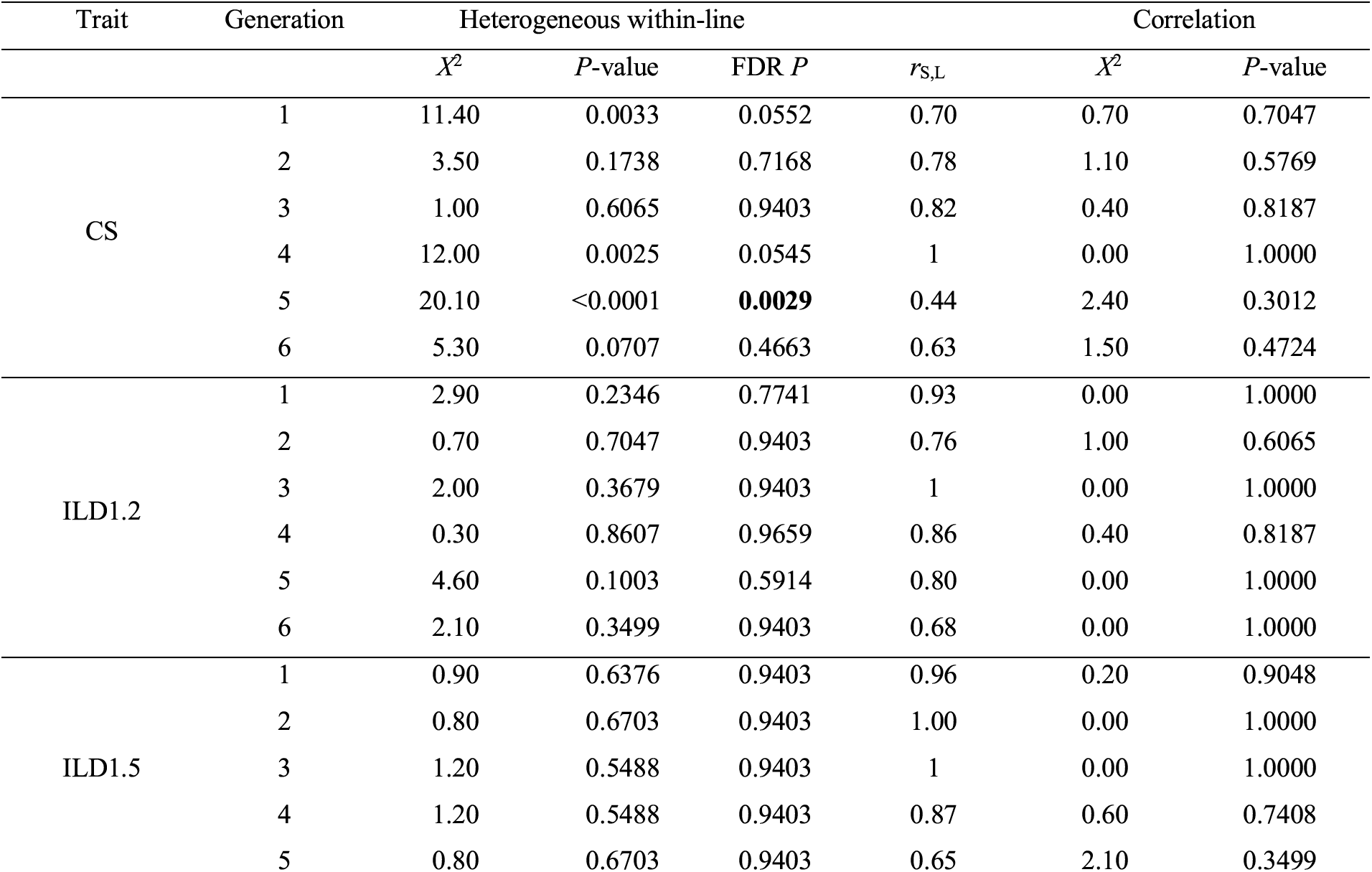
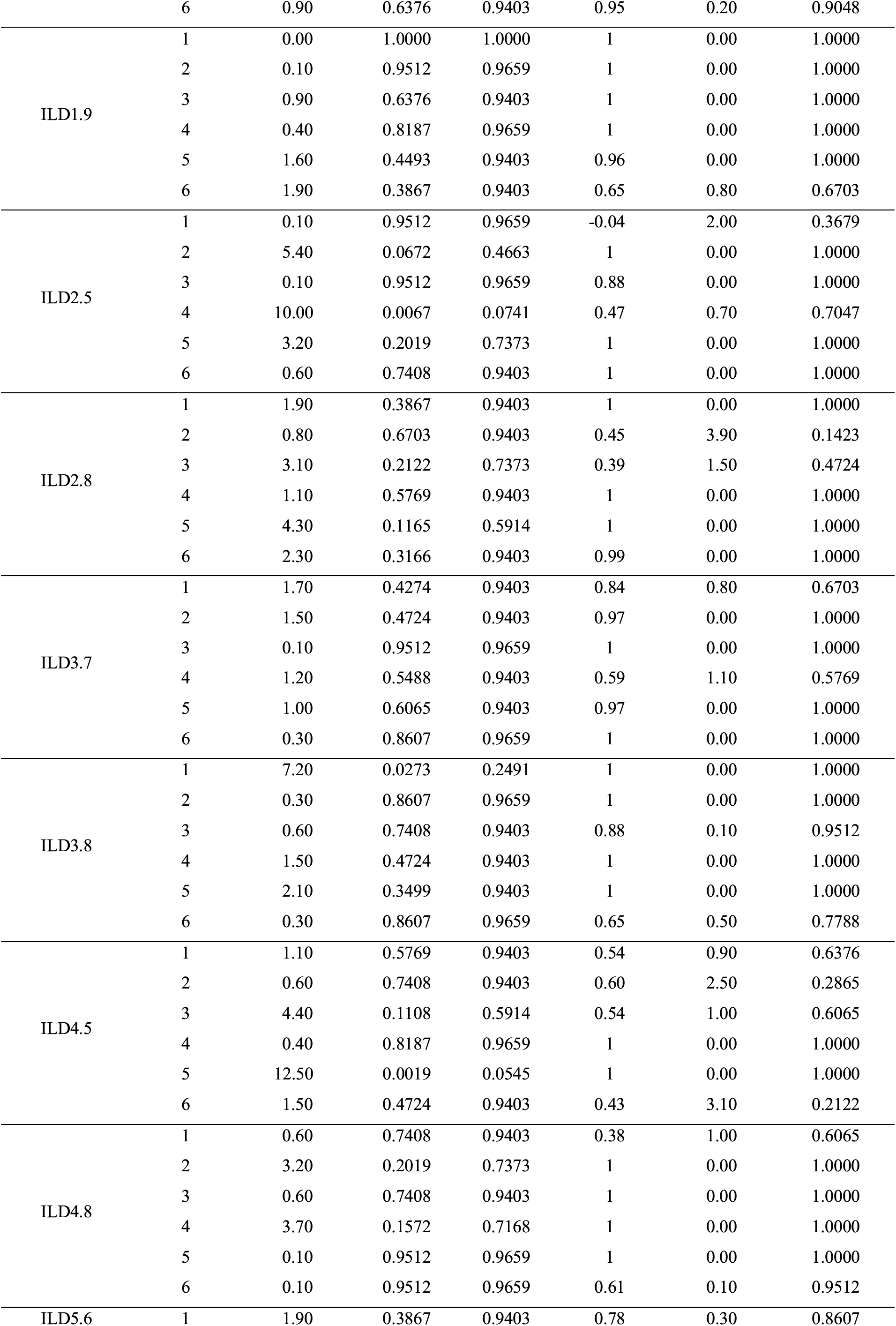
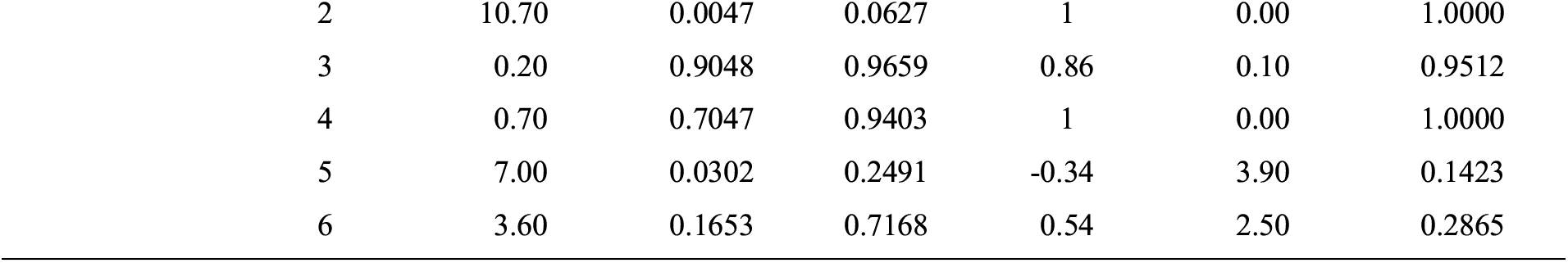
Statistical support for the effect of *Drosophila serrata* population size treatment on the mutations segregating within and among sub-lines. Log-likelihood ratio tests were used to test two hypotheses: whether the within-line variance differed in magnitude between the two population size treatments (Heterogeneous within-line) and; whether the mutational correlation between the treatments differed from 1 (Correlation). The test statistic (*X*^2^) follows a chi-square distribution with the degrees of freedom equal to the difference in the number of parameters between these models (Heterogeneous within-line: d.f. = 2; Correlation < 0.999: d.f. = 1). The *P*-value is shown, and for the Heterogeneous within-line test, the 5% FDR corrected *P*-value. Following FDR correction, there was only one instance of statistically supported difference between the treatments, highlighted in bold. No FDR correction was applied to Correlation < 0.999 as all *P* > 0.05. The among-line (mutational) correlation between the Small and Large treatments (*rs,L*) is shown and should be interpreted within the context of the corresponding variances (Table S4; Figure 5). The per-treatment estimates of *V_E_* from the model analysed here (5) were indistinguishable from the estimates from model (2) (Table S4) and are not re-reported here.

| Trait | Generation | Heterogeneous within-line | | | $r_{S,L}$ | Correlation | |
| --- | --- | --- | --- | --- | --- | --- | --- |
| | | $\chi^2$ | $P$-value | FDR $P$ | | $\chi^2$ | $P$-value |
| CS | 1 | 11.40 | 0.0033 | 0.0552 | 0.70 | 0.70 | 0.7047 |
|  | 2 | 3.50 | 0.1738 | 0.7168 | 0.78 | 1.10 | 0.5769 |
|  | 3 | 1.00 | 0.6065 | 0.9403 | 0.82 | 0.40 | 0.8187 |
|  | 4 | 12.00 | 0.0025 | 0.0545 | 1 | 0.00 | 1.0000 |
|  | 5 | 20.10 | <0.0001 | **0.0029** | 0.44 | 2.40 | 0.3012 |
|  | 6 | 5.30 | 0.0707 | 0.4663 | 0.63 | 1.50 | 0.4724 |
| ILD1.2 | 1 | 2.90 | 0.2346 | 0.7741 | 0.93 | 0.00 | 1.0000 |
|  | 2 | 0.70 | 0.7047 | 0.9403 | 0.76 | 1.00 | 0.6065 |
|  | 3 | 2.00 | 0.3679 | 0.9403 | 1 | 0.00 | 1.0000 |
|  | 4 | 0.30 | 0.8607 | 0.9659 | 0.86 | 0.40 | 0.8187 |
|  | 5 | 4.60 | 0.1003 | 0.5914 | 0.80 | 0.00 | 1.0000 |
|  | 6 | 2.10 | 0.3499 | 0.9403 | 0.68 | 0.00 | 1.0000 |
| ILD1.5 | 1 | 0.90 | 0.6376 | 0.9403 | 0.96 | 0.20 | 0.9048 |
|  | 2 | 0.80 | 0.6703 | 0.9403 | 1.00 | 0.00 | 1.0000 |
|  | 3 | 1.20 | 0.5488 | 0.9403 | 1 | 0.00 | 1.0000 |
|  | 4 | 1.20 | 0.5488 | 0.9403 | 0.87 | 0.60 | 0.7408 |
|  | 5 | 0.80 | 0.6703 | 0.9403 | 0.65 | 2.10 | 0.3499 |
|  | 6 | 0.90 | 0.6376 | 0.9403 | 0.95 | 0.20 | 0.9048 |
| ILD1.9 | 1 | 0.00 | 1.0000 | 1.0000 | 1 | 0.00 | 1.0000 |
|  | 2 | 0.10 | 0.9512 | 0.9659 | 1 | 0.00 | 1.0000 |
|  | 3 | 0.90 | 0.6376 | 0.9403 | 1 | 0.00 | 1.0000 |
|  | 4 | 0.40 | 0.8187 | 0.9659 | 1 | 0.00 | 1.0000 |
|  | 5 | 1.60 | 0.4493 | 0.9403 | 0.96 | 0.00 | 1.0000 |
|  | 6 | 1.90 | 0.3867 | 0.9403 | 0.65 | 0.80 | 0.6703 |
| ILD2.5 | 1 | 0.10 | 0.9512 | 0.9659 | -0.04 | 2.00 | 0.3679 |
|  | 2 | 5.40 | 0.0672 | 0.4663 | 1 | 0.00 | 1.0000 |
|  | 3 | 0.10 | 0.9512 | 0.9659 | 0.88 | 0.00 | 1.0000 |
|  | 4 | 10.00 | 0.0067 | 0.0741 | 0.47 | 0.70 | 0.7047 |
|  | 5 | 3.20 | 0.2019 | 0.7373 | 1 | 0.00 | 1.0000 |
|  | 6 | 0.60 | 0.7408 | 0.9403 | 1 | 0.00 | 1.0000 |
| ILD2.8 | 1 | 1.90 | 0.3867 | 0.9403 | 1 | 0.00 | 1.0000 |
|  | 2 | 0.80 | 0.6703 | 0.9403 | 0.45 | 3.90 | 0.1423 |
|  | 3 | 3.10 | 0.2122 | 0.7373 | 0.39 | 1.50 | 0.4724 |
|  | 4 | 1.10 | 0.5769 | 0.9403 | 1 | 0.00 | 1.0000 |
|  | 5 | 4.30 | 0.1165 | 0.5914 | 1 | 0.00 | 1.0000 |
|  | 6 | 2.30 | 0.3166 | 0.9403 | 0.99 | 0.00 | 1.0000 |
| ILD3.7 | 1 | 1.70 | 0.4274 | 0.9403 | 0.84 | 0.80 | 0.6703 |
|  | 2 | 1.50 | 0.4724 | 0.9403 | 0.97 | 0.00 | 1.0000 |
|  | 3 | 0.10 | 0.9512 | 0.9659 | 1 | 0.00 | 1.0000 |
|  | 4 | 1.20 | 0.5488 | 0.9403 | 0.59 | 1.10 | 0.5769 |
|  | 5 | 1.00 | 0.6065 | 0.9403 | 0.97 | 0.00 | 1.0000 |
|  | 6 | 0.30 | 0.8607 | 0.9659 | 1 | 0.00 | 1.0000 |
| ILD3.8 | 1 | 7.20 | 0.0273 | 0.2491 | 1 | 0.00 | 1.0000 |
|  | 2 | 0.30 | 0.8607 | 0.9659 | 1 | 0.00 | 1.0000 |
|  | 3 | 0.60 | 0.7408 | 0.9403 | 0.88 | 0.10 | 0.9512 |
|  | 4 | 1.50 | 0.4724 | 0.9403 | 1 | 0.00 | 1.0000 |
|  | 5 | 2.10 | 0.3499 | 0.9403 | 1 | 0.00 | 1.0000 |
|  | 6 | 0.30 | 0.8607 | 0.9659 | 0.65 | 0.50 | 0.7788 |
| ILD4.5 | 1 | 1.10 | 0.5769 | 0.9403 | 0.54 | 0.90 | 0.6376 |
|  | 2 | 0.60 | 0.7408 | 0.9403 | 0.60 | 2.50 | 0.2865 |
|  | 3 | 4.40 | 0.1108 | 0.5914 | 0.54 | 1.00 | 0.6065 |
|  | 4 | 0.40 | 0.8187 | 0.9659 | 1 | 0.00 | 1.0000 |
|  | 5 | 12.50 | 0.0019 | 0.0545 | 1 | 0.00 | 1.0000 |
|  | 6 | 1.50 | 0.4724 | 0.9403 | 0.43 | 3.10 | 0.2122 |
| ILD4.8 | 1 | 0.60 | 0.7408 | 0.9403 | 0.38 | 1.00 | 0.6065 |
|  | 2 | 3.20 | 0.2019 | 0.7373 | 1 | 0.00 | 1.0000 |
|  | 3 | 0.60 | 0.7408 | 0.9403 | 1 | 0.00 | 1.0000 |
|  | 4 | 3.70 | 0.1572 | 0.7168 | 1 | 0.00 | 1.0000 |
|  | 5 | 0.10 | 0.9512 | 0.9659 | 1 | 0.00 | 1.0000 |
|  | 6 | 0.10 | 0.9512 | 0.9659 | 0.61 | 0.10 | 0.9512 |
| ILD5.6 | 1 | 1.90 | 0.3867 | 0.9403 | 0.78 | 0.30 | 0.8607 |

|  |  |  |  |  |  |  |
| --- | --- | --- | --- | --- | --- | --- |
| 2 | 10.70 | 0.0047 | 0.0627 | 1 | 0.00 | 1.0000 |
| 3 | 0.20 | 0.9048 | 0.9659 | 0.86 | 0.10 | 0.9512 |
| 4 | 0.70 | 0.7047 | 0.9403 | 1 | 0.00 | 1.0000 |
| 5 | 7.00 | 0.0302 | 0.2491 | -0.34 | 3.90 | 0.1423 |
| 6 | 3.60 | 0.1653 | 0.7168 | 0.54 | 2.50 | 0.2865 |

**Figure S1.**
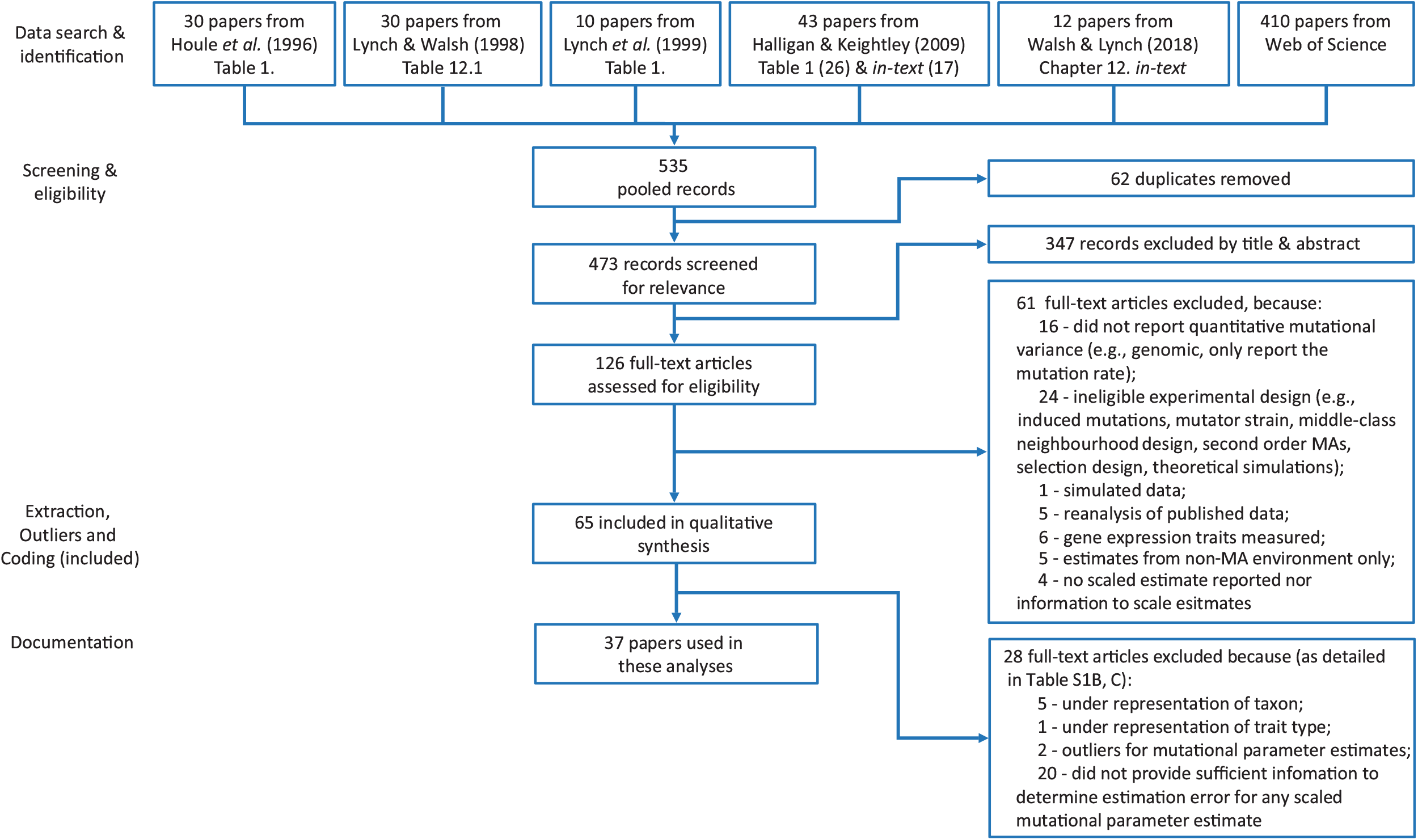
Diagram of systematic literature search for the meta-analysis of mutational variance estimates from mutation accumulation (MA) experiments. The schematic follows the PRISMA (Preferred Reporting Items for Systematic Reviews and Meta-Analysis)^1^

---

**Figure S2.**
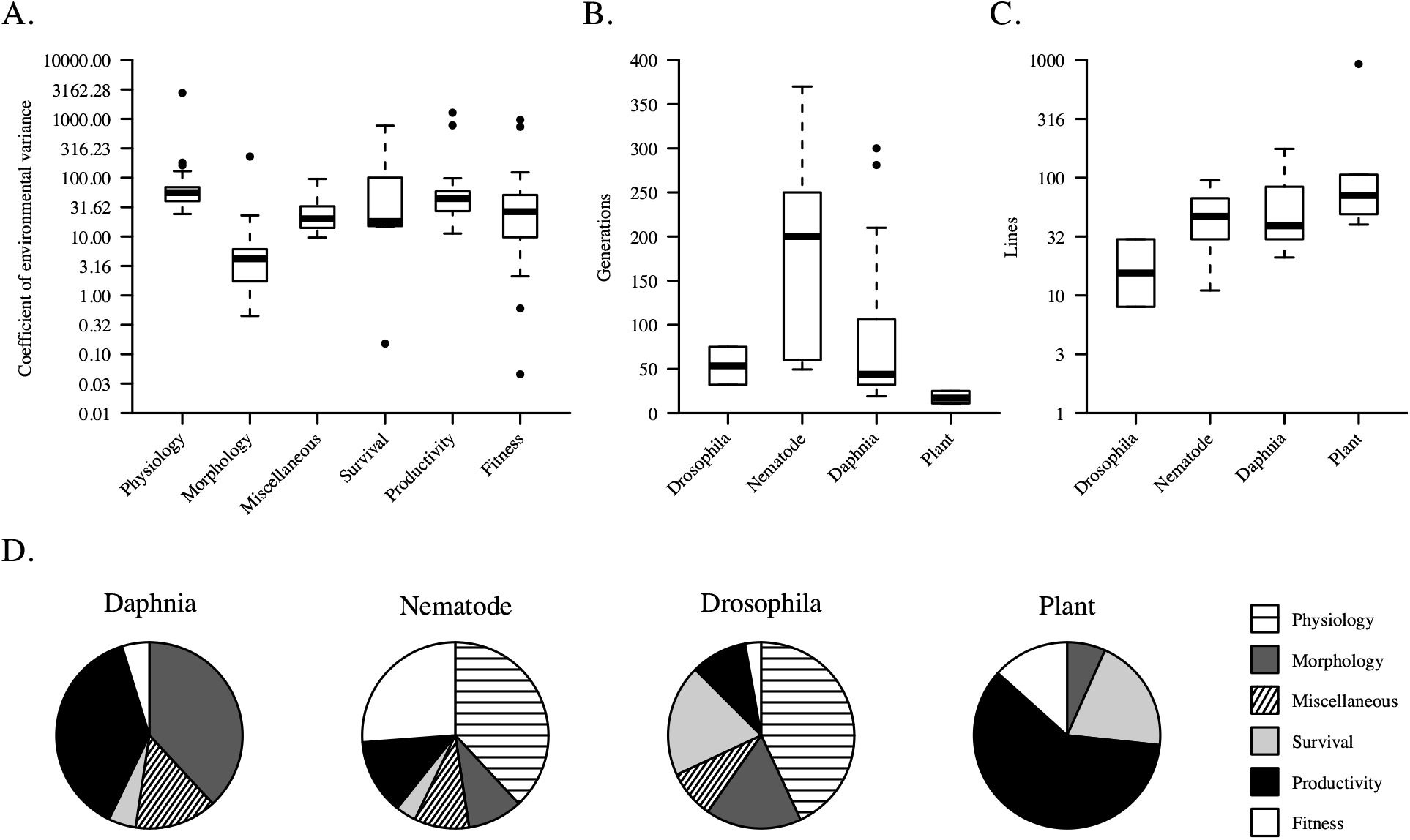
Variation of factors that may influence the magnitude of mutational variance estimates. (A) *CV_E_* was calculated as^1^: 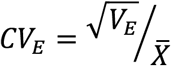 and log-transformed to improve resolution for plotting (NB: the values are shown in the y-axis were back-transformed to the original *CV_E_* scale). *CV_E_* was plotted per trait category, which are defined in Methods and Table S2. (B) the number of generations and (C) the (log-transformed; but as in (A) the y-axis is shown on the original scale) number of lines of mutation accumulation experiments (studies) in different taxon categories. Taxon categories are defined in Methods. Box plots represent the median (bar), interquartile range (IQR, box) and 3 IQR (whiskers) values of the published estimates, with more extreme values indicated by solid circles. (D) The distribution of estimate numbers across the different trait categories within each taxon group, where traits definitions as per Table S2.

**Figure S3.**
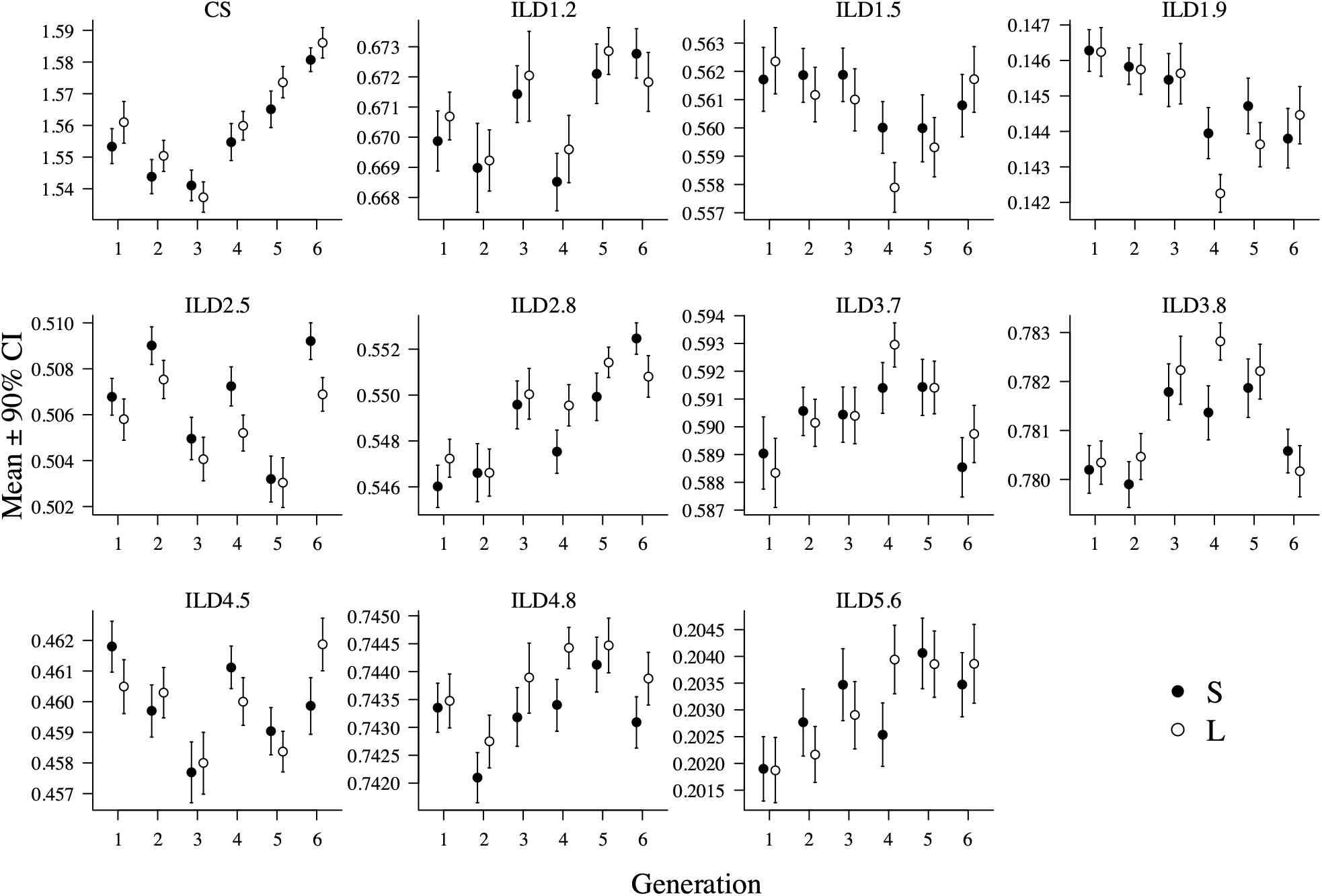
Variation in wing trait mean across six generations of an experiment in *Drosophila serrata.* For each of the 11 wing traits (panels), the least-squares mean (± 95% CI) from model (2) is plotted for each generation for the Small (solid circle) and Large (open circles) population size treatments. Centroid size (CS) is measured in millimetres; the ten inter-landmark distances are in units of centroid size (multiplied by 100; see Figure 2B for trait definitions). Plotted estimates are reported in Table S4.

**Figure S4.**
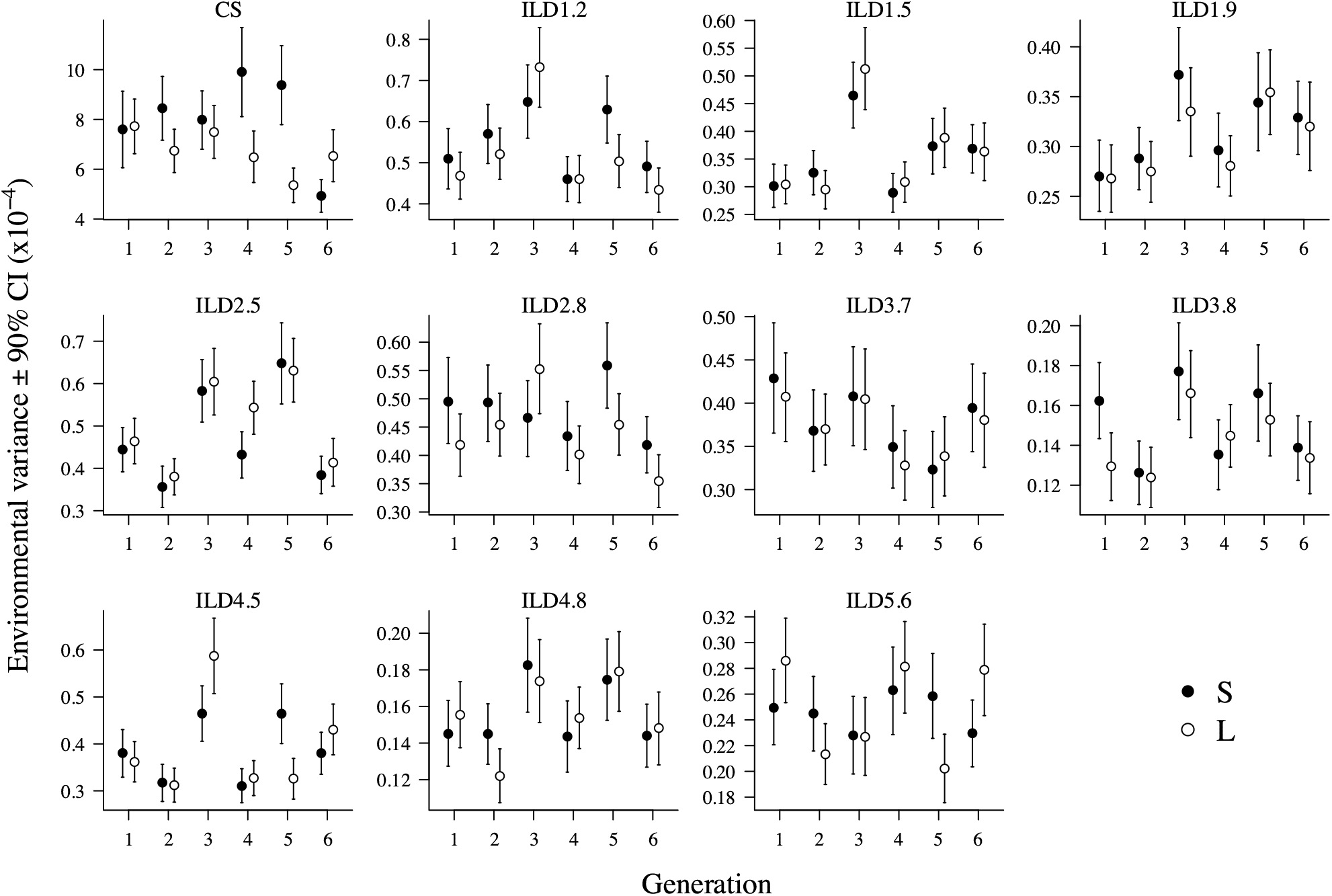
Variation in environmental variance across six generations of an experiment in *Drosophila serrata.* For each of the 11 wing traits (panels), *V_E_* (± 95% CI), estimated as the sum of among vial and residual variances from model (2), is plotted for each generation for the Small (solid circle) and Large (open circles) population size treatments. Plotted estimates are reported in Table S4.

**Figure S5.**
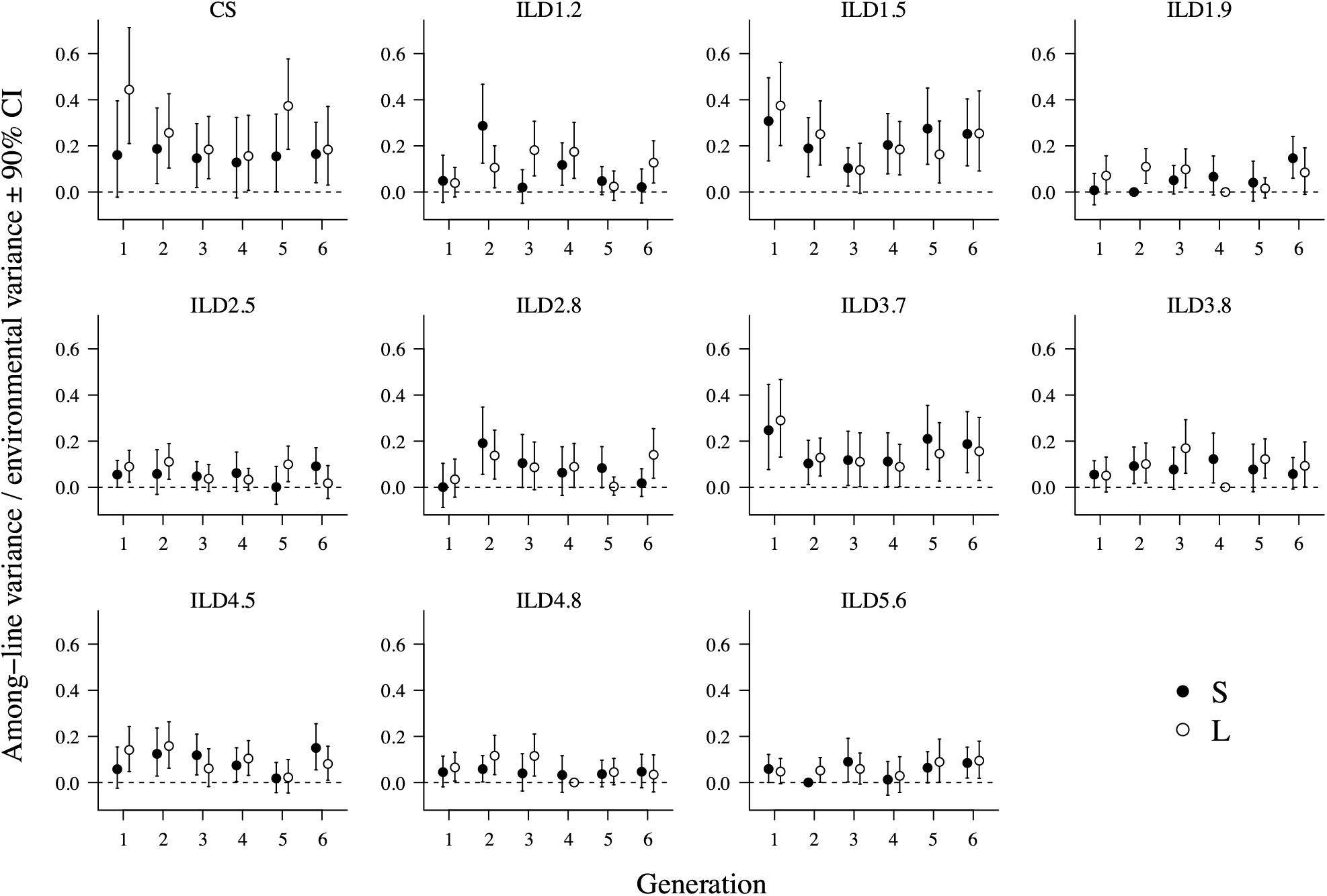
Variance scaled estimates of among-line variance estimates in *D. serrata.* Among-line variance estimates (*V_L_*, Figure 5) were placed on a heritability scale to account for the influence of trait variance on the magnitude of estimates, dividing *V_L_* by the corresponding estimate of environmental variance (summed among-vial and residual variance reported in Table S4). Plotted are these variance-scaled REML estimates (90% CI) for each trait (panel) (trait descriptions in Figure 2B) in each generation (x-axis) for each of the two population size treatments (Small: solid circles; Large: open circles). The dashed horizontal line indicates zero.

**Figure S6.**
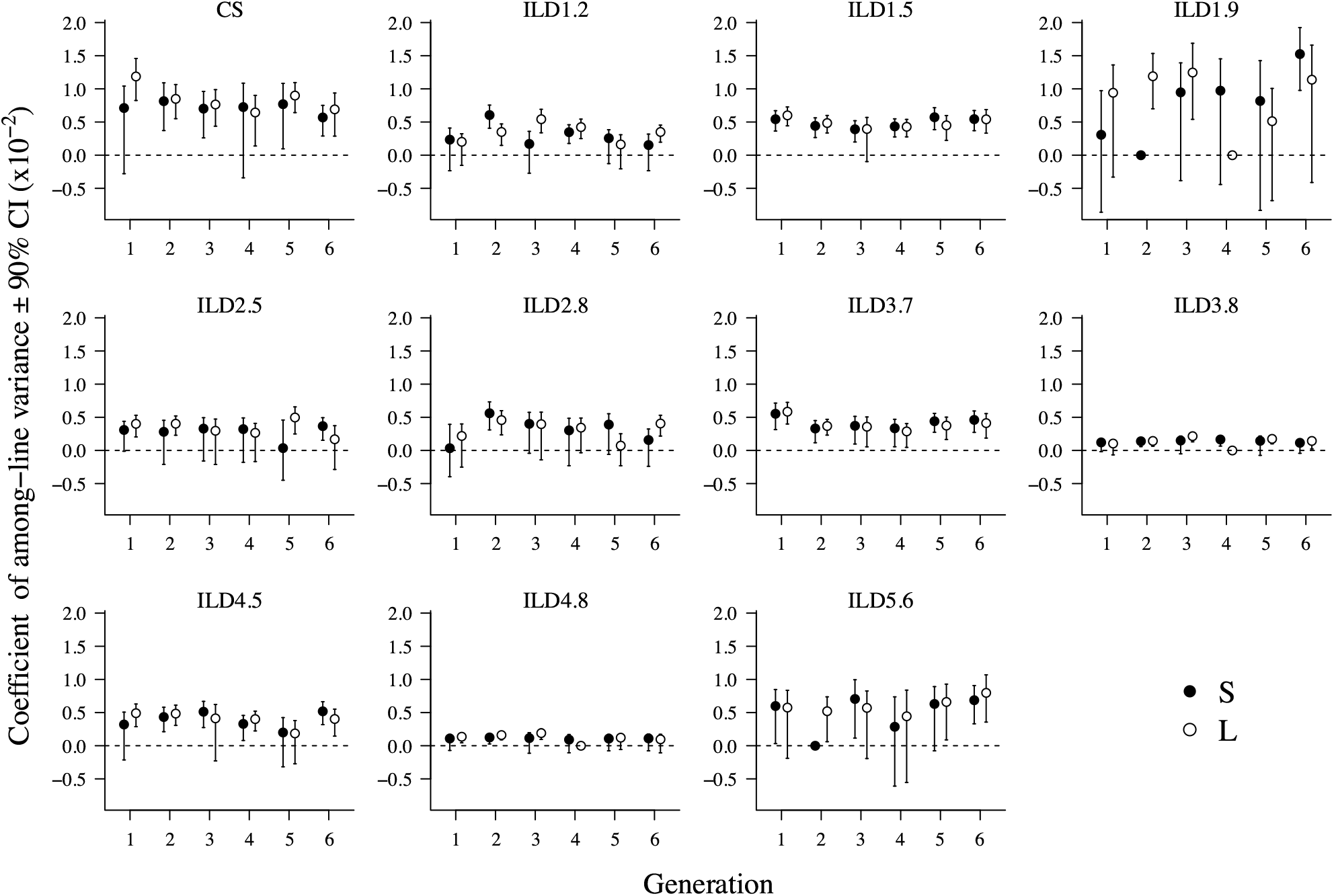
Mean-scaled estimates of among-line variance estimates in *D. serrata.* Among-line variance estimates (*V_L_*, Figure 5) were placed on a coefficient of variance scale to account for the influence of trait mean on magnitude of estimates, using the following formula: 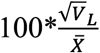 where 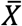 is the trait mean (Table S4). Plotted are the REML point estimates (and 90% CI) for each trait (panel) (see Figure 2B for trait descriptions) in each generation (x-axis) for each of the two population size treatments (Small: solid circles; Large: open circles). The dashed horizontal line indicates zero.

**Figure S7.**
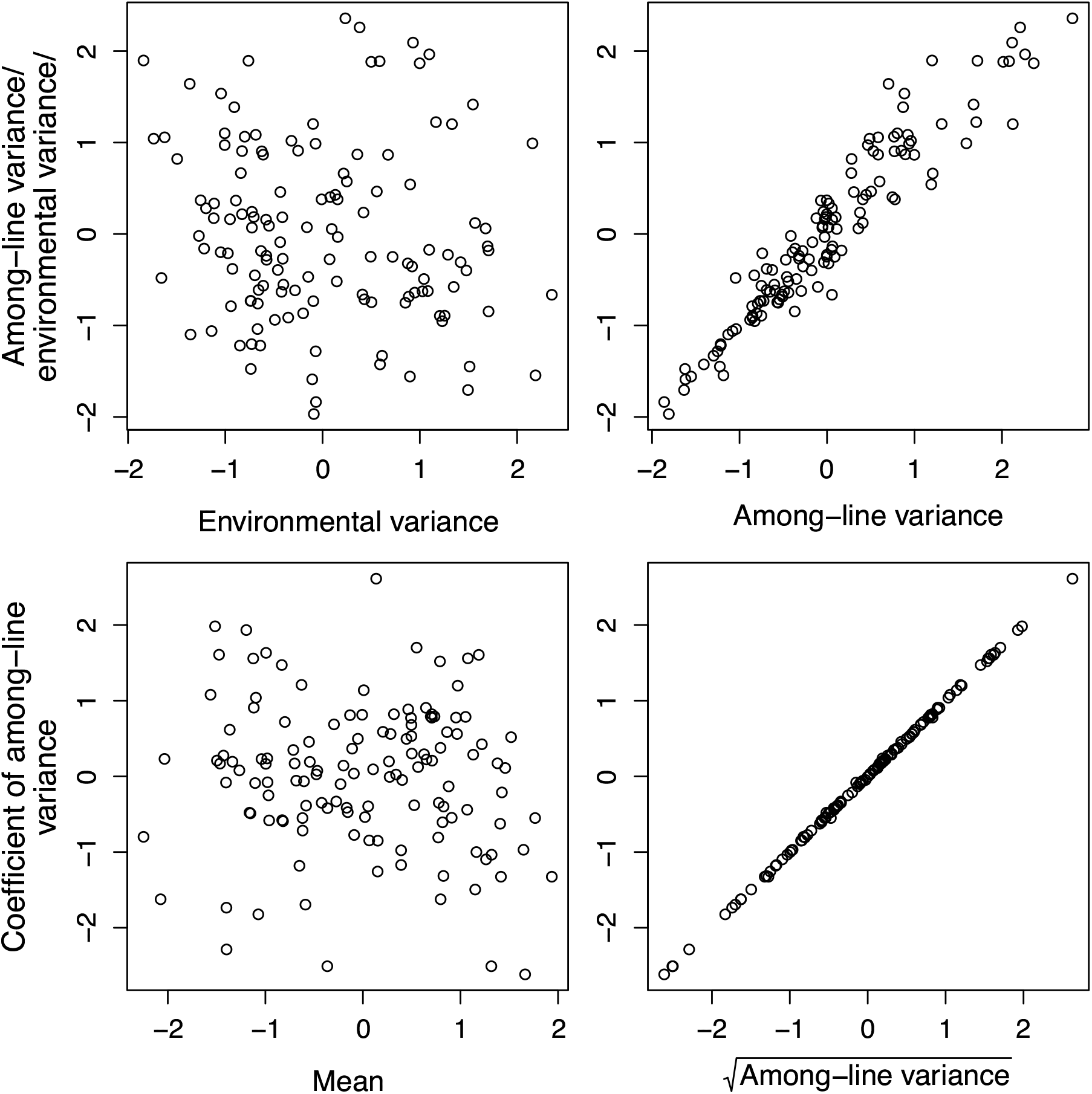
The relative contributions of heterogeneity in the scaling parameter (*V_E_* or trait mean; left panels) and the among-line variance (*V_L_*; right panels) to the variation observed in the variance-scaled (top panels) or mean-scaled (bottom panel) estimates of *Drosophila serrata* among-line variance. All estimates are presented in Table S4, *V_L_* is also presented in Figure 5; *V_E_* and mean in Figures S3 and S4, respectively; variance or mean scaled estimates of *V_L_* in Figures S5 and S6, respectively. The 12 estimates for each of the 11 traits for each parameter (*V_L_*, *V_E_*, mean, variance-scaled and mean-scaled) were first placed on a standardised scale by centring on the mean of the 12 estimates, and dividing by the standard deviation of the 12 estimates (i.e., z-scores were calculated), and all 132 estimates (12 estimates per 11 traits) are plotted.

**Figure S8.**
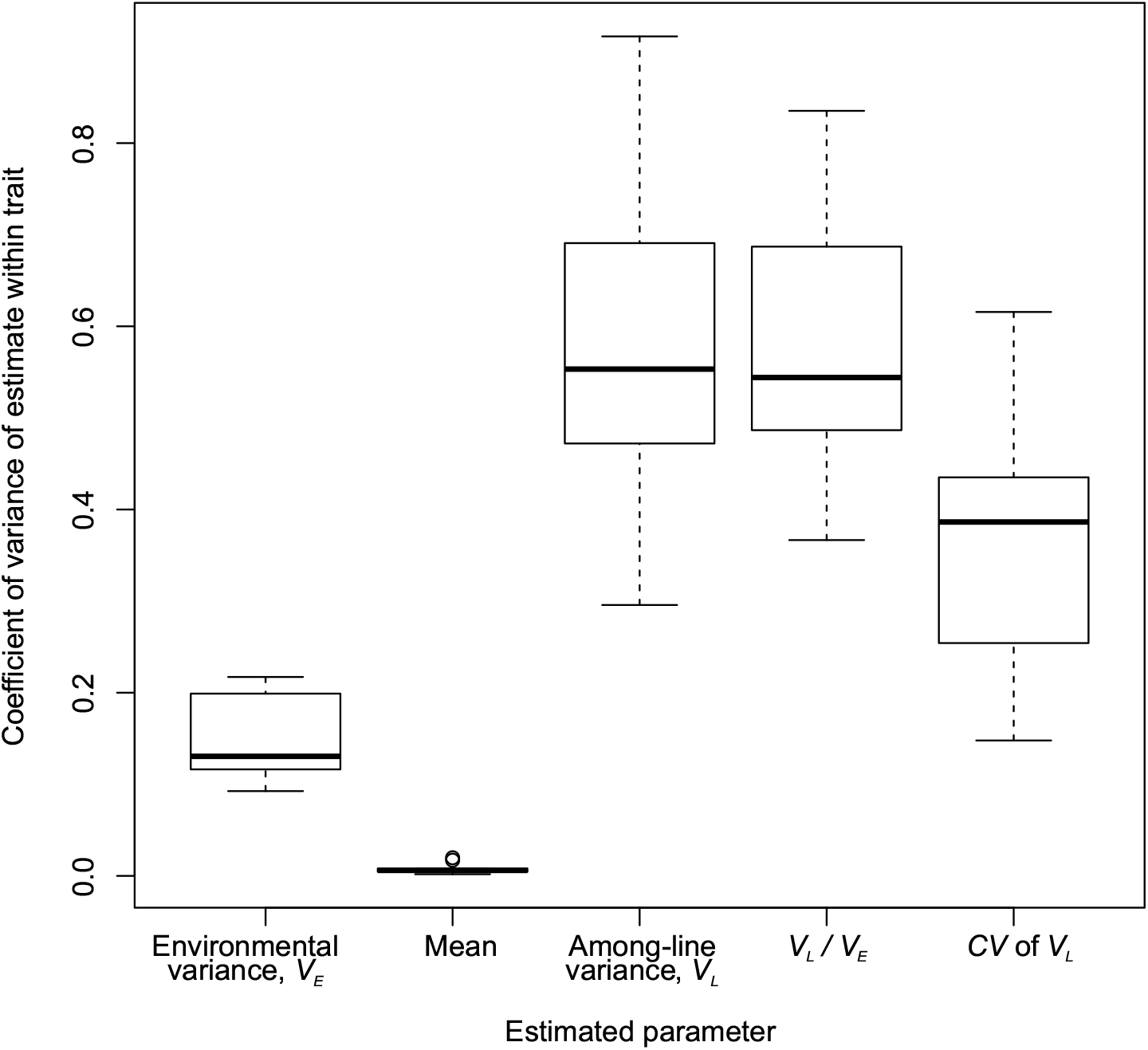
Variation in estimates of each genetic variation parameter in *D. serrata.* For each of these parameters, the 12 estimates per trait (Table S4) were analysed to calculate the coefficient of variance across the 12 observations as: standard deviation / mean. Box plots represent the median (bar), interquartile range (IQR, box) and 3 IQR (whiskers) values for the 11 traits.

## References

Azevedo, R. B., P. D. Keightley, C. Laurén-Määttä, L. L. Vassilieva, M. Lynch et al., 2002 Spontaneous mutational variation for body size in Caenorhabditis elegans. Genetics 162: 755–765.

Baer, C. F., 2008 Quantifying the decanalizing effects of spontaneous mutations in rhabditid nematodes. American Naturalist 172: 272–281.

Baer, C. F., J. Joyner-Matos, D. Ostrow, V. Grigaltchik, M. P. Salomon et al., 2010 Rapid decline in fitness of mutation accumulation lines of gonochoristic (outcrossing) Caenorhabditis nematodes. Evolution 64: 3242–3253.

Barton, N. H., and P. D. Keightley, 2002 Understanding quantitative genetic variation. Nature Reviews Genetics 3: 11–21.

Barton, N. H., and M. Turelli, 1989 Evolutionary quantitative genetics: how little do we know? Annual Review of Genetics 23: 337–370.

Beltran, T., V. Shahrezaei, V. Katju and P. Sarkies, 2020 Epimutations driven by small RNAs arise frequently but most have limited duration in *Caenorhabditis elegans*. Nature Ecology & Evolution 4: 1539–1548.

Benjamini, Y., and Y. Hochberg, 1995 Controlling the false discovery rate: a practical and powerful approach to multiple testing. Journal of the Royal Statistical Society: Series B (Statistical Methodology) 57: 289–300.

Besnard, F., J. Picao-Osorio, C. Dubois and M. A. Felix, 2020 A broad mutational target explains a fast rate of phenotypic evolution. Elife 9: e54928.

Bitner-Mathé, B. C., and L. B. Klaczko, 1999 Plasticity of *Drosophila melanogaster* wing morphology: effects of sex, temperature and density. Genetica 105: 203.

Boyle, E. A., Y. I. Li and J. K. Pritchard, 2017 An expanded view of complex traits: From polygenic to omnigenic. Cell 169: 1177–1186.

Braendle, C., C. F. Baer and M.-A. Félix, 2010 Bias and evolution of the mutationally accessible phenotypic space in a developmental system. PLoS Genetics 6: e1000877.

Cavicchi, S., D. Guerra, G. Giorgi and C. Pezzoli, 1985 Temperature-related divergence in experimental populations of *Drosophila melanogaster.* I. Genetic and developmental basis of wing size and shape variation. Genetics 109: 665–689.

Dugand, R. J., J. D. Aguirre, E. Hine, M. W. Blows and K. McGuigan, 2021 The contribution of mutation and selection to multivariate quantitative genetic variance in an outbred population of *Drosophila serrata*. Proceedings of the National Academy of Sciences, USA 118: e2026217118.

Estes, S., B. C. Ajie, M. Lynch and P. C. Phillips, 2005 Spontaneous mutational correlations for life-history, morphological and behavioral characters in *Caenorhabditis elegans*. Genetics 170: 645–653.

Estes, S., P. C. Phillips, D. R. Denver, W. K. Thomas and M. Lynch, 2004 Mutation accumulation in populations of varying size: The distribution of mutational effects for fitness correlates in *Caenorhabditis elegans*. Genetics 166: 1269–1279.

Fry, J. D., S. L. Heinsohn and T. F. Mackay, 1996 The contribution of new mutations to genotype-environment interaction for fitness in *Drosophila melanogaster*. Evolution 50: 2316–2327.

García-Dorado, A., J. Fernández and C. López-Fanjul, 2000 Temporal uniformity of the spontaneous mutational variance of quantitative traits in *Drosophila melanogaster*. Genetics Research 75: 47–51.

Garcia-Gonzalez, F., L. W. Simmons, J. L. Tomkins, J. S. Kotiaho and J. P. Evans, 2012 Comparing evolvabilities: Common errors surrounding the calculation and use of coefficients of additive genetic variation. Evolution 66: 2341–2349.

Hall, D. W., R. Mahmoudizad, A. W. Hurd and S. B. J. G. r. Joseph, 2008 Spontaneous mutations in diploid *Saccharomyces cerevisiae:* another thousand cell generations. Genetics Research 90: 229–241.

Halligan, D. L., and P. D. Keightley, 2009 Spontaneous mutation accumulation studies in evolutionary genetics. Annual Review of Ecology, Evolution and Systematics 40: 151–172.

Hansen, T. F., C. Pelabon and D. Houle, 2011 Heritability is not Evolvability. Evolutionary Biology 38: 258–277.

Hine, E., D. E. Runcie, K. McGuigan and M. W. Blows, 2018 Uneven distribution of mutational variance across the transcriptome of *Drosophila serrata* revealed by high-dimensional analysis of gene expression. Genetics 209: 1319–1328.

Ho, E. K. H., F. Macrae, L. C. Latta, P. Mcilroy, D. Ebert et al., 2020 High and highly variable spontaneous mutation rates in Daphnia. Molecular Biology and Evolution 37: 3258–3266.

Hoffmann, A. A., J. Merila and T. N. Kristensen, 2016 Heritability and evolvability of fitness and nonfitness traits: Lessons from livestock. Evolution 70: 1770–1779.

Houle, D., 1991 Genetic covariance of fitness correlates: what genetic correlations are made of and why it matters. Evolution 45: 630–648.

Houle, D., 1992 Comparing evolvability and variability of quantitative traits. Genetics 130: 195–204.

Houle, D., 1998 How should we explain variation in the genetic variance of traits? Genetica 102: 241–253.

Houle, D., G. H. Bolstad, K. van der Linde and T. F. Hansen, 2017 Mutation predicts 40 million years of fly wing evolution. Nature 548: 447–450.

Houle, D., and K. Meyer, 2015 Estimating sampling error of evolutionary statistics based on genetic covariance matrices using maximum likelihood. Journal of Evolutionary Biology 28: 1542–1549.

Houle, D., B. Morikawa and M. Lynch, 1996 Comparing mutational variabilities. Genetics 143: 1467–1483.

Houle, D., and S. V. Nuzhdin, 2004 Mutation accumulation and the effect of *copia* insertions in *Drosophila melanogaster*. Genetical Research 83: 7–18.

Huang, W., R. F. Lyman, R. A. Lyman, M. A. Carbone, S. T. Harbison et al., 2016 Spontaneous mutations and the origin and maintenance of quantitative genetic variation. Elife 5: e14625.

Johnson, T., and N. Barton, 2005 Theoretical models of selection and mutation on quantitative traits. Philosophical Transactions of the Royal Society of London, Series B: Biological Sciences 360: 1411–1425.

Katju, V., and U. Bergthorsson, 2019 Old trade, new tricks: insights into the spontaneous mutation process from the partnering of classical mutation accumulation experiments with high-throughput genomic approaches. Genome Biology and Evolution 11: 136–165.

Katju, V., L. B. Packard, L. Bu, P. D. Keightley and U. Bergthorsson, 2015 Fitness decline in spontaneous mutation accumulation lines of *Caenorhabditis elegans* with varying effective population sizes. Evolution 69: 104–116.

Kavanaugh, C. M., and R. G. Shaw, 2005 The contribution of spontaneous mutation to variation in environmental responses of *Arabidopsis thaliana:* Responses to light. Evolution 59: 266–275.

Keightley, P. D., T. F. Mackay and A. Caballero, 1993 Accounting for bias in estimates of the rate of polygenic mutation. Proceedings of the Royal Society of London, Series B: Biological Sciences 253: 291–296.

Kimura, M., 1983 The Neutral Theory of Molecular Evolution. Cambridge University Press, New York.

Klein, T. W., 1974 Heritability and genetic correlation: Statistical power, population comparisons and sample size. Behavior Genetics 4: 171–189.

Klein, T. W., J. C. De Vries and C. T. Finkbeiner, 1973 Heritability and genetic correlation: Standard errors of estimates and sample size. Behavior Genetics 3: 355–364.

Kondrashov, A. S., and D. Houle, 1994 Genotype-environment interactions and the estimation of the genomic mutation rate in *Drosophila melanogaster*. Proceedings of the Royal Society of London, Series B: Biological Sciences 258: 221–227.

Littell, R., G. Milliken, W. Stroup, R. Wolfinger and O. Schabenberger, 2006 SAS for Mixed Models. SAS Institute, Cary, NC.

Luijckx, P., E. K. H. Ho, A. Stanic and A. F. Agrawal, 2018 Mutation accumulation in populations of varying size: Large effect mutations cause most mutational decline in the rotifer *Brachionus calyciflorus* under UV-C radiation. Journal of Evolutionary Biology 31: 924–932.

Lynch, M., 1988 The rate of polygenic mutation. Genetical Research 51: 137–148.

Lynch, M., 2010 Evolution of the mutation rate. Trends in Genetics 26: 345–352.

Lynch, M., J. Blanchard, D. Houle, T. Kibota, S. Schultz et al., 1999 Perspective: spontaneous deleterious mutation. Evolution 53: 645–663.

Lynch, M., and W. G. Hill, 1986 Phenotypic evolution by neutral mutation. Evolution 40: 915–935.

Lynch, M., and B. Walsh, 1998 Genetics and analysis of quantitative traits. Sinauer, Sunderland, MA.

Mackay, T., R. F. Lyman and W. G. Hill, 1995 Polygenic mutation in *Drosophila melanogaster:* non-linear divergence among unselected strains. Genetics 139: 849–859.

Mackay, T. F. C., E. A. Stone and J. F. Ayroles, 2009 The genetics of quantitative traits: challenges and prospects. Nature Reviews Genetics 10: 565–577.

Martin, G., and T. Lenormand, 2006 The fitness effect of mutations across environments: a survey in light of fitness landscape models. Evolution 60: 2413–2427.

McGuigan, K., and M. W. Blows, 2013 Joint allelic effects on fitness and metric traits. Evolution 67: 1131–1142.

McGuigan, K., D. Petfield and M. W. Blows, 2011 Reducing mutation load through sexual selection on males. Evolution 65: 2816–2829.

Merilä, J., and B. C. Sheldon, 1999 Genetic architecture of fitness and nonfitness traits: empirical patterns and development of ideas. Heredity 83: 103–109.

Meyer, K., and D. Houle, 2013 Sampling based approximation of confidence intervals for functions of genetic covariance matrices. Proceedings of the Association for the Advancement of Animal Breeding and Genetics 20: 523–526.

Mezey, J. G., D. Houle and S. V. Nuzhdin, 2005 Naturally segregating quantitative trait loci affecting wing shape of *Drosophila melanogaster*. Genetics 169: 2101–2113.

Mukai, T., 1964 Genetic structure of natural populations of *Drosophila melanogaster.* I. Spontaneous mutation rate of polygenes controlling viability. Genetics 50: 1–19.

Ness, R. W., A. D. Morgan, N. Colegrave and P. D. Keightley, 2012 Estimate of the spontaneous mutation rate in *Chlamydomonas reinhardtii*. Genetics 192: 1447–1454.

Neto-Silva, R. M., B. S. Wells and L. A. Johnston, 2009 Mechanisms of growth and homeostasis in the *Drosophila* wing. Annual Review of Cell and Developmental Biology 25: 197–220.

Ostrow, D., N. Phillips, A. Avalos, D. Blanton, A. Boggs et al., 2007 Mutational bias for body size in Rhabditid nematodes. Genetics 176: 1653–1661.

Reddiex, A. J., S. L. Allen and S. F. Chenoweth, 2018 A genomic reference panel for Drosophila serrata. G3 8: 1335–1346.

Rockman, M. V., 2012 The QTN program and the alleles that matter for evolution: All that’s gold does not glitter. Evolution 66: 1–17.

Rohlf, F., 2007 tpsRelw version 1.45. Department of Ecology and Evolution, State University of New York, Stony Brook.

Rohlf, F. J., 1999 Shape statistics: Procrustes superimpositions and tangent spaces. Journal of Classifation 16: 197–223.

Rohlf, F. J., 2013 tpsDig, digitize landmarks and outlines, version 2.17. Department of Ecology and Evolution, State University of New York at Stony Brook.

Schrider, D. R., D. Houle, M. Lynch and M. W. J. G. Hahn, 2013 Rates and genomic consequences of spontaneous mutational events in Drosophila melanogaster. 194: 937–954.

Self, S. G., and K.-Y. Liang, 1987 Asymptotic properties of maximum likelihood estimators and likelihood ratio tests under nonstandard conditions. Journal of the American Statistical Association 82: 605–610.

Simons, Y. B., K. Bullaughey, R. R. Hudson and G. Sella, 2018 A population genetic interpretation of GWAS findings for human quantitative traits. Plos Biology 16: e2002985.

Sung, W., M. S. Ackerman, S. F. Miller, T. G. Doak and M. Lynch, 2012 Drift-barrier hypothesis and mutation-rate evolution. Proceedings of the National Academy of Sciences, USA 109: 18488–18492.

Sztepanacz, J. L., and M. W. Blows, 2017 Accounting for sampling error in genetic eigenvalues using random matrix theory. Genetics 206: 1271–1284.

van der Graaf, A., R. Wardenaar, D. A. Neumann, A. Taudt, R. G. Shaw et al., 2015 Rate, spectrum, and evolutionary dynamics of spontaneous epimutations. Proceedings of the National Academy of Sciences, USA 112: 6676–6681.

Vassilieva, L. L., A. M. Hook and M. Lynch, 2000 The fitness effects of spontaneous mutations in *Caenorhabditis elegans*. Evolution 54: 1234–1246.

Walsh, B., and M. Lynch, 2018 Evolution and Selection of Quantitative Traits. Oxford University Press.

Wayne, M. L., and T. F. Mackay, 1998 Quantitative genetics of ovariole number in *Drosophila melanogaster.* II. Mutational variation and genotype-environment interaction. Genetics 148: 201–210.

Wright, S., 1931 Evolution in Mendelian populations. Genetics 16: 97–159.

Yang, J., B. Benyamin, B. P. McEvoy, S. Gordon, A. K. Henders et al., 2010 Common SNPs explain a large proportion of the heritability for human height. Nature Genetics 42: 565–569.

## References

1 Moher, D., A. Liberati, J. Tetzlaff, D. G. Altman and P. Group, 2009 Preferred reporting items for systematic reviews and meta-analyses: the PRISMA statement. PLoS Medicine 6: e1000097.

## References

1 Houle, D., B. Morikawa and M. Lynch, 1996 Comparing mutational variabilities. Genetics 143: 1467–1483

